# Increased activity without changes in coupling between somata and apical dendrites during visual mismatches

**DOI:** 10.64898/2026.11.04.764139

**Authors:** Timothy M. Henley, Joshua Tindall, Jason E. Pina, Roman Pogodin, Shiella Caldejon, Natalia Orlova, Avalon Amaya, Ahad Bawany, Conor Grasso, Peter Groblewski, Vivian Ha, Warren Han, Tye Johnson, Robyn Naidoo, Katrina Nguyen, Sam Seid, Lucas Suarez, Jackie Swapp, Xana Waughman, Josh Wilkes, Ali Williford, Ahrial Young, Jérôme A. Lecoq, Joel Zylberberg, Blake A. Richards

## Abstract

Leading theories of cortical function posit that sensory processing involves a comparison between top-down predictions and bottom-up sensory data. Tests of these theories have been hampered by experimental challenges related to simultaneously recording top-down and bottom-up signals in response to sensory stimuli that either do or do not match internally-generated predictions. Here, we conduct dual-plane 2-photon calcium imaging near-simultaneously from distal apical dendrites and somata of pyramidal neurons in each of 4 different regions of mouse visual cortex while the mice are presented with visual sequences that can either match or mismatch previously habituated visual stimuli. Specifically, we examine the responses in the distal apical dendrites and cell bodies of pyramidal neurons while mice view either movie clips that reverse unexpectedly or Gabor patch sequences that have orientations that do not match habituated patterns. We find that in response to unexpected reversals in natural movies, both the apical dendrites and somata show increased activity that decreases over repeated exposures, consistent with a prediction error response. However, we do not find any clear evidence for increased activity in response to mismatches in Gabor sequences, beyond possible release from adaptation effects. We additionally use volumetric images to connect dendritic and somatic compartments and examine the coupling between connected dendrites and cell bodies in layer 5 pyramidal neurons. We do not find evidence for changes in coupling in response to mismatch stimuli, either for natural movies or Gabor sequences. Our results are consistent across four regions of visual cortex. We also observe that increased responses to reversals in natural movies are stronger in layer 2/3 than layer 5 pyramidal neurons. Altogether, these data suggest that visual cortex is sensitive to deviations from previously-experienced naturalistic inputs. Our data do not suggest that this mismatch sensitivity extends to artificial stimuli, nor do they support theories of prediction error causing changes in the coupling between apical dendrites and somata.

## 1 Introduction

A central hypothesis in systems neuroscience holds that the cortex does not passively relay signals about sensory inputs, but instead actively compares them to learned internal predictions, generating signals that reflect the mismatch between expectation and reality ^1,2,3,4^. This predictive processing framework has guided decades of experiments on the sensory cortex. However, within this broad framework, there remain many differing views as to where and by what physiological mechanisms such comparisons might be implemented.

Certain theories posit the existence of explicit error signals that represent the difference or mismatch between sensorimotor inputs and predictions being received by a given cortical area ^2^. Previous experimental studies suggest that the computation of these error signals might be implemented through the interaction of distinct subpopulations in the cortical microcircuit, or may involve dedicated error neurons ^5,6,7,8,9^. Other theoretical work proposes that mismatches are processed within single neurons, through the interaction between signals integrated in somatic and apical dendritic compartments ^10,11,12,13,14^.

To date, evidence from neural recordings is mixed. Some previous studies have reported increased activity in cortical neurons in response to mismatch stimuli ^15,16,17,18^. However, more recent work has argued that some of these responses may simply be an effect of release from adaptation, and that when adaptation is absent, unpredictable inputs may only drive increased activity in higher-order areas ^19,20^. Other research suggests that naturalistic stimuli are essential to studying vision and specifically to eliciting responses to visual mismatch ^21,22^. It is possible that purely visual mismatch responses are better characterized as stimulus-specific enhancement ^23^. Further, mounting evidence points to an active online aspect of visual perception, as investigated with closed-loop sensorimotor mismatch, which may be critical to eliciting certain elevated mismatch responses ^24,25,26,27^.

*In vivo* data regarding the potential role of apical dendrites in implementing predictive processing are extremely limited, since few studies have combined observation of dendritic activity with expectation-violating stimuli ^9^. One previous study ^17^ observed that responses of pyramidal cells’ apical dendrites and somata to unexpected stimuli appear to evolve differently over time. However, the evidence for this effect came from recordings that were not simultaneous and ultimately from different animals ^28^, leaving open the question of how responses of these compartments relate in the same population of neurons, and within single cells. Moreover, the mismatch stimuli changed both in location and orientation relative to the habituated stimuli, leaving open the possibility that their results were driven by release from adaptation due to repeated stimulation of specific receptive fields over a prolonged period of time ^17^.

Here, we conducted experiments using near-simultaneous 2-photon calcium imaging of apical dendritic and somatic compartments of pyramidal neurons in multiple visual areas. We habituated mice over several days to two types of sequential stimuli: natural movies and sequences of image frames containing sparse Gabor wavelets. After habituation, we included unpredictable changes in a subset of the sequences to introduce violations of expectations. Notably, all violation stimuli were carefully designed to control for the possibility of response differences arising from release from adaptation or stimulus differences. The natural movie stimuli provided complex and dynamic mixtures of lower- and higher-order features across spatial and temporal scales, matching more closely signals the visual system evolved to respond to ^29,30,31^.

We found that somata and apical dendrites of cortical pyramidal cells in layer 2/3 (L2/3), and partially in layer 5 (L5), respond more strongly to identical segments from natural movies when those segments violated expectations. However, we found no such increases in response to unexpected stimuli for less-naturalistic Gabor sequences. Strikingly, these responses evolved similarly in both cellular compartments and across four different visual areas of the mouse brain. When comparing somatic and dendritic activity, we did not see evidence for changes in somato-dendritic coupling, i.e., coactivation, in response to unexpected stimuli, whether natural or artificial.

Increased neural responses when habituated expectations are violated, which systematically decrease over days of exposure to violation, suggest that neocortical circuits are sensitive to deviations from learned visual regularities, consistent with predictive processing. That this sensitivity is present for natural movie reversals but absent for Gabor sequence violations indicates that the visual system has expectations grounded in the statistics of natural stimuli. Given the similarity of visual mismatch responses in somata and apical dendrites, a strong causal contribution from dissociated dendritic prediction-error computation is unlikely, at least under passive open-loop conditions.

## 2 Results

### 2.1 Simultaneous imaging of apical dendrite segments and cell bodies, in multiple visual areas, over multiple days, with volumetric reconstruction

Many prior studies found elevated responses to mismatch stimuli ^15,16,23^. However, in some cases, these elevated responses may reflect release from adaptation, caused by repeated presentations of the same stimulus during the expectation building portion(s) of the experiment ^19^. We designed two different sequential visual stimuli to probe the reaction of neural circuits to deviations from previously habituated stimuli, in a manner that controls for the effects of adaptation.

The first set of stimuli were short natural movies, such as behaving wild mice, and worms moving in soil (Fig. 1A). The mice were habituated over several days (E1-E3) to these clips played in the forward direction (“fwd”). We suspected that the movies’ naturalistic content would be salient to mice, and further, that the continuous structure of motion, surfaces, and causal sequences would align with both their learned and evolved visual processing, while the structural violations would be directly opposed to these ^32^. To generate violations of expectations in later experimental sessions, the animals were presented with versions of the same movies in which playback would reverse partway through the clip (“fwd-rev”). These were introduced in a random subset (25%) of sequences presented in the final three experimental sessions (U1-U3) (Fig. 1G). However, reversals flip the temporal order of the sequences. To perform a properly controlled comparison of responses, we therefore included a control condition: the full movies in reverse (“rev”). For each movie, we could thus compare the responses to *identical* movie segments in the mismatch condition, in which the segment appeared as a violation of the habituated sequence, and in the “control” condition, in which there should be no established expectation and no reversal in the sequence. We additionally included a secondary control stimulus: reversed movies with short forward sequential violations (“rev-fwd”, see Methods 4.2.1). These contained temporal reversals, but without strongly established expectations, since the segments preceding the reversal were not habituated.

**Figure 1.**
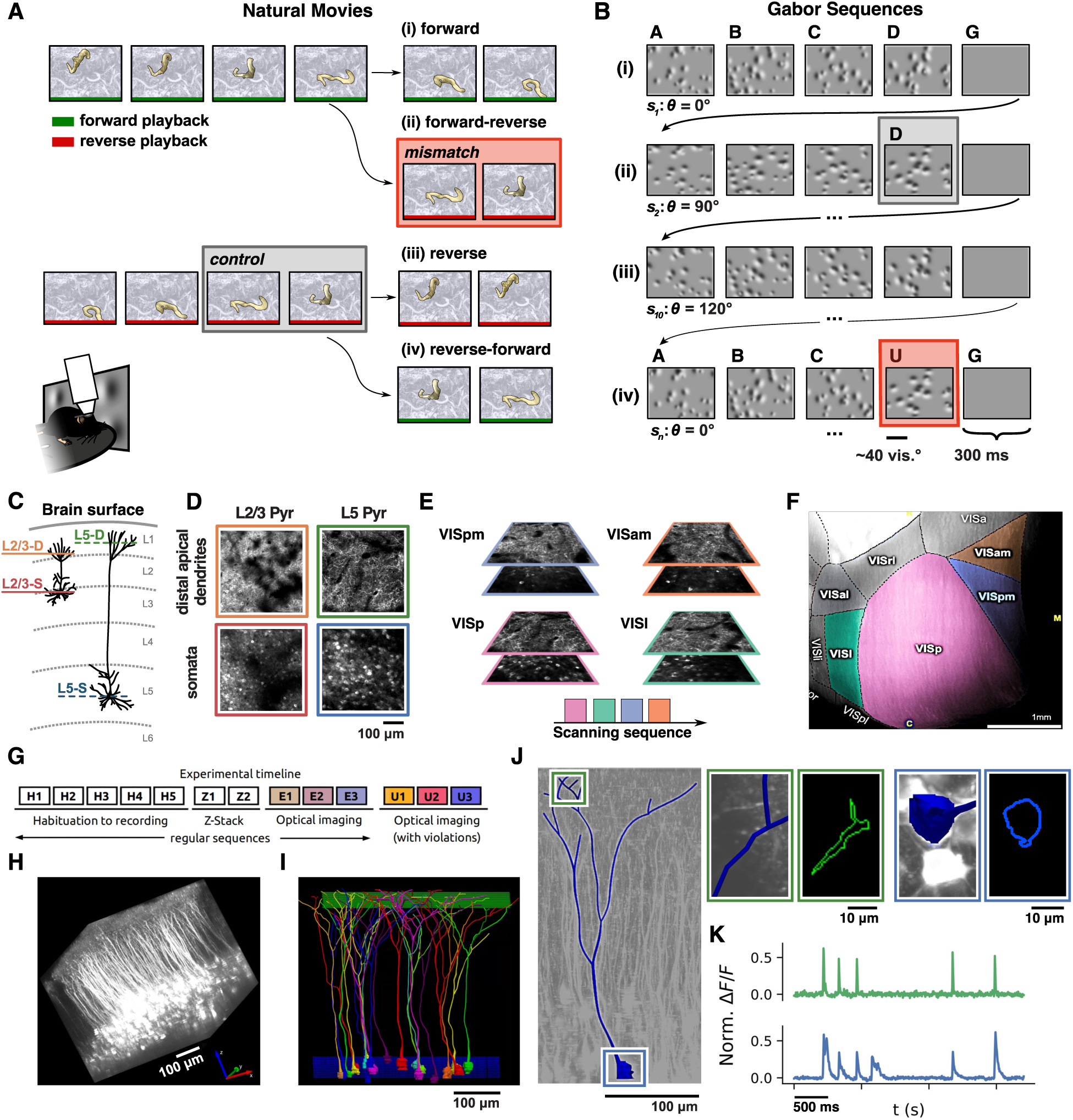
Experimental design. **(A)** Natural movie stimuli. Green bars indicate periods of forward playback, and red bars periods of reverse playback. Subjects were habituated to the forward version of the clip. A border and internal highlighting have been added to the worm in the example for visual clarity, and were not present in the actual stimulus. **(B)** Gabor sequence stimulus. **(C)** Activity was recorded in L1 for apical dendrites and L2/3 or L5 for somata, depending on the mouse line. **(D)** Example somatic and apical dendritic imaging planes for the two cell lines, for a single visual area. **(E)** Visualization of the scanning sequence of the multi-plane mesoscope, showing near-simultaneous acquisition of planes across compartments and areas of visual cortex (0.1 s per scanning sequence). **(F)** A map of the visual cortex, with the acquired areas indicated by coloring. **(G)** Experimental timeline (see Methods 4.1). **(H)** Example 2-photon calcium volumetric z-stack, acquired from VISp in an awake behaving mouse (see Methods 4.1). **(I)** Examples of cells traced from somatic to apical dendritic plane, with z-stack layers near the imaging planes marked in blue and green (see Methods 4.4). **(J)** A traced cell which was matched to both an apical dendritic and somatic ROI. **(K)** Time traces for an apical dendritic ROI (top, green) and somatic ROI (bottom, blue) matched to the same traced cell, with several visible coincident events.

The second set of stimuli were sequences of gray frames containing randomly located Gabor patches, similar to the stimuli used in other studies ^17,28,33^. The mice were habituated to five-frame sequences, A-B-C-D-G. For each frame in the sequence, the locations of the patches were the same in all presentations within a given session, with G frames containing no Gabor wavelets. Within each sequence, the orientations of the patches were sharply distributed about a common mean, shared across all frames (Fig. 1B). To avoid short-term adaptation responses, the mean orientation was randomly selected for each new sequence. To create inputs that violated the statistics of the habituated sequences, we then presented A-B-C-U-G sequences, in which the distribution of orientations was consistent for three frames, then rotated by 90 degrees for the U frame. To minimize the potential for release from adaptation as a result of the changing stimuli, the locations of the Gabor patches in the D and U frames were identical, such that any responses could not be a result of new receptive fields being stimulated by the U frames. This design also allowed us to compare responses to the same stimulus frames in the mismatch vs control conditions because the D frames from some trials had the same patch locations and mean orientations as the U frames from other trials. However, even with these additional controls, it should be noted that the Gabor stimuli could still induce release from adaptation when the mismatch was presented, since the mismatch stimuli had a very different orientation from that shared by the sequence preceding it. Thus, to provide an additional control for possible adaptation to the repetition of the same orientations within a sequence, we also designed a second Gabor sequence in which the mean orientation changed with each frame, while the mean of the mismatch frame violated expectations by *not* changing.

We used a multi-plane optical physiology platform to near-simultaneously observe activity in the distal apical dendrites (imaging in layer 1), and the cell bodies of GCaMP6f-expressing pyramidal neurons in the supragranular (L2/3, Cux2 mice) or subgranular (L5, Rbp4 mice) layers of the visual cortex in awake, behaving mice (Fig. 1C,D). We chose to sample supra- and subgranular layers because both have been implicated in predictive learning theories ^34^, but with possibly different functional roles, as pyramidal neurons in each exhibit differences in morphology, physiology, and local and long-range connectivity ^18,35,36,37,38^. The feedforward output of L2/3 pyramidal neurons and top-down feedback connections have made them a primary focus of many predictive processing studies; however, the extensive and more segregated apical dendrites in L5 pyramidal neurons are also often hypothesized to make it possible for bottom-up and top-down inputs to have distinct effects on these cells ^7,9,25,34,36,39,40^.

Our experimental recordings were segmented and matched across days, so that we could observe the evolution of responses of individual somatic or apical dendritic regions of interest (ROIs) (see Methods 4.1) ^41,42^. Combined with our simultaneous imaging, we could thus compare mismatch responses from somata and distal apical dendrites from the same populations of neurons, as they evolved over days of exposure to mismatch stimuli.

Motivated by theories of predictive learning based on feedback to apical dendrites ^10,11,12,13,14,43^, we further sought to study responses to mismatch stimuli in somatic and apical dendritic compartments belonging to the same cell. In order to match somatic and dendritic ROIs, we developed methods to produce high-resolution volumetric functional imaging of the same neural populations from our experimental recordings. From these we extracted z-stacks and used semi-automatic neural tracings to connect somatic and dendritic ROIs belonging to the same cell (see Fig. 1H-J, Methods 4.4). By combining these with our simultaneous recordings, we could observe the responses of connected somatic and dendritic compartments over days of exposure to mismatch stimuli, and in particular the degree of coactivation in these compartments across conditions (Fig. 1K).

We also imaged across multiple visual areas: VISp, VISl, VISpm and VISam. This allowed us to monitor somato-dendritic activity in both lower- and higher-order regions of the mouse visual cortex (Fig. 1E,F). In primates, evidence suggests the levels of the cortical visual hierarchy are selective to different stimulus properties, with incrementally more compound, mixed or abstract properties driving neural responses at each higher level ^2,3,20,44,45,46^. Although the mouse visual hierarchy may be comparatively simpler, shallower, more distributed, and parallel, it also shares many common features with primates ^22,46,47,48,49,50,51^. We investigated whether mismatch responses differ across hierarchical levels, as predicted by hierarchical predictive coding theories ^2,34^.

### 2.2 Responses to mismatch in natural movies are elevated in both somata and distal apical dendrites

To assess whether violations of visual expectations alone induce mismatch responses, we first studied the responses to our movie clips in VISp. We compared the activity of both somata and distal apical dendrites during the reversals in the natural movies (a mismatch condition) to their activity measured while the identical clip was played as part of the fully reversed movie (a control condition in which no expectation could have been established by habituation) (Fig. 2A). Responses to the reversed clip were consistently larger in the mismatch condition than in the control condition (Fig. 2B, C).

**Figure 2.**
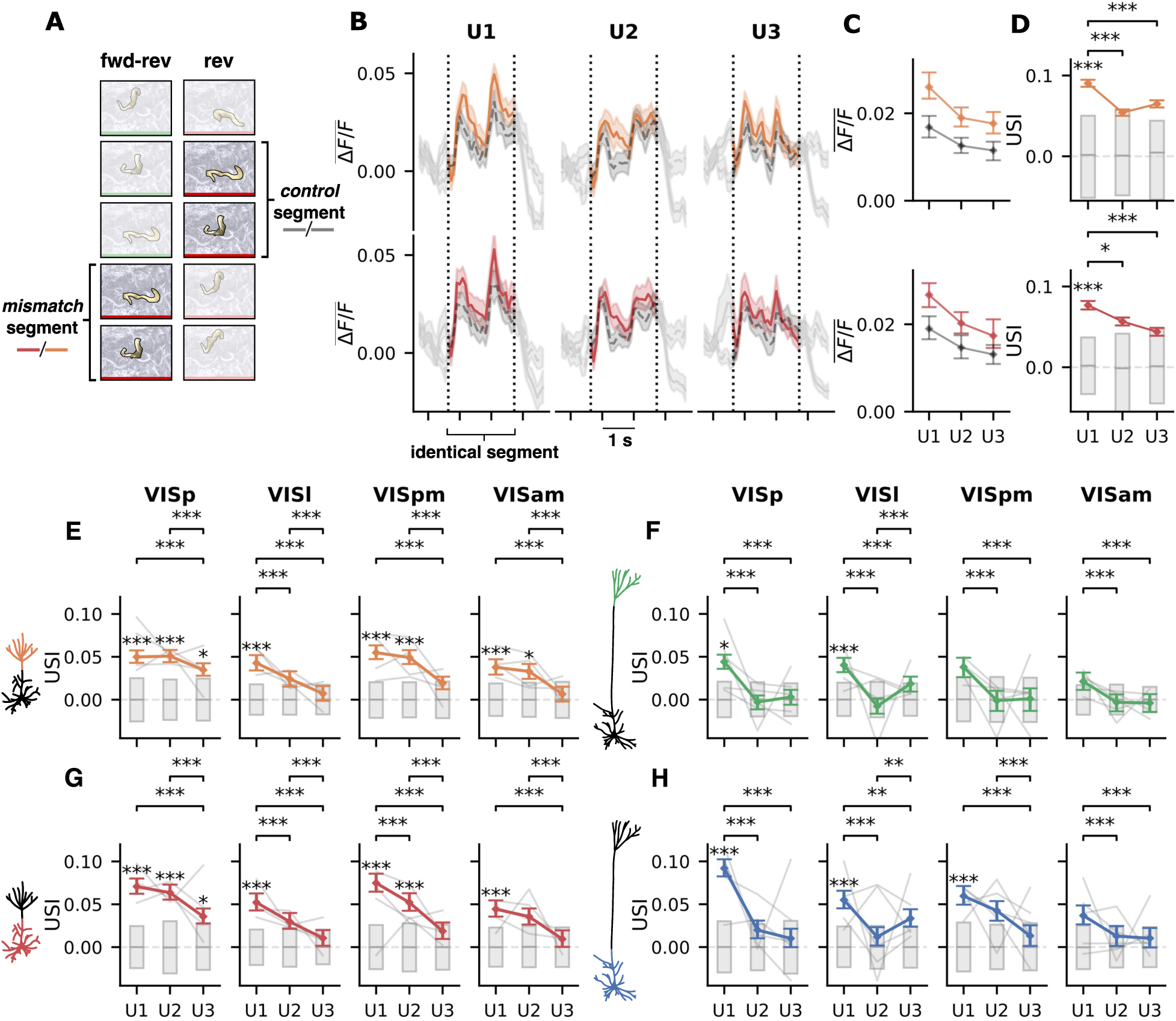
Neuronal responses to identical segments of the “worms” movie in different conditions and unexpected selectivity index (USI) analysis. **(A)** Depiction of the control (reverse) and mismatch (forward-reverse) movie sequences, with labels marking the identical segment of clip used in subsequent comparisons. Green frames represent forward playback, while red frames represent reverse playback. **(B)** Mean Δ*F/F* across ROIs and trials for L2/3 apical dendrites (upper, orange) and somata (lower, red) during presentation of the same movie segment in mismatch and control conditions. Colored series represent the responses in the mismatch condition, when the segment was presented as a violation of the habituated forward version of the movie. Grey series represent the responses to the same segment when presented as part of a full reverse presentation of the movie, not preceded by a habituated sequence. Dashed vertical lines indicate the window in which the stimuli match across conditions, with grey tails for each series outside of the window added for context. Error bands represent a 95 % CI estimated using a hierarchical bootstrap, samples constructed with replacement over ROIs and trials. **(C)** Mean of the windowed portion of the Δ*F/F* in (B) over ROIs, trials, as well as time, yielding one value per session, for sessions U1, U2, and U3. Error bars represent a 95 % CI estimated with a hierarchical bootstrap as in (B). **(D)** Unexpected selectivity index (USI) computed for the same Δ*F/F* as in (B) and (C). Colored series represent the mean USI across ROIs for each session, grey bars represent the 95% CI of a distribution constructed from permutations (see Methods 4.3.1). Error bars represent a 95 % CI estimated with a hierarchical bootstrap as in (B) and (C). **(E)-(H)** Selectivity of responses to the mismatch condition (USIs), across areas and cell lines, averaged over L2/3 distal apical dendritic ROIs **(E)** and L2/3 somata **(G)**, and L5 apical dendrites **(F)** and L5 somata **(H)**. Grey bars represent 95% CI for a permutation based null distribution. Error bars represent a 95 % CI estimated using a hierarchical bootstrap. Fine grey lines show average individual subject USIs. * *p <* 0.05, ** *p <* 0.01, *** *p <* 0.005 (see Methods 4.3.1).

We quantified the observed differences in responses in the two stimulus conditions, using an “unexpected selectivity index” (USI), which measured the magnitude of responses to stimuli in the mismatch condition versus the control condition—the higher the USI, the larger the relative response in the mismatch condition (see Methods 4.3.1). We found that USI values were significantly positive in the first session of exposure to the mismatch condition, and, on average, decreased across sessions with additional exposure to the mismatch movie (Fig. 2D). This is consistent with predictive processing theories, and suggests that neurons in VISp are indeed more active when a stimulus is presented in a sequence that deviates from previous experience.

### 2.3 Increased responses to reversals in natural movies in all areas and compartments

We extended our USI analysis across laminae, compartments and visual areas to identify which combinations showed evidence of mismatch responses. For natural movie stimuli, we found evidence for a mismatch response that was similar across compartments and areas, and broadly consistent across laminae (though the mismatch response remained stronger over days in L2/3; Fig. 2E-H). Mismatch responses were strongest upon introduction of violations, and decreased over days of exposure to the mismatch stimuli.

L2/3 pyramidal neurons are the principal source of bottom-up feedforward signals to higher cortical areas ^52,53,54,55,56^. In hierarchical predictive coding (HPC), the output of a cortical area is the error signal. Consequently, L2/3 is often targeted to isolate mismatch or prediction error responses ^24,34,43,57^. In L2/3 somata we observed significantly positive USIs during the session where violations were first introduced (U1). This corresponds to an elevated response to identical stimuli, when presented as a violation of a habituated sequence, compared to when not a violation (Fig. 2C,G).

L5 pyramidal neurons are a source of contextual feedback signals, and may be uniquely suited to integrating top-down predictive information ^2,24,34,43,57^. L5 somata also showed significantly positive USIs on U1, in all areas except for VISam, as well as in VISp and VISl apical dendrites. In both laminae, USIs decreased significantly between sessions, particularly from U1 to U3, indicating a reduced mismatch response (Fig. 2F, H).

Compartmental theories of predictive processing have proposed that mismatch may be computed within single cells, through the integration of different signals received in somatic and apical dendritic compartments ^10,11,12,14^. Here we found that responses in both apical and somatic compartments were broadly similar. In L2/3, both somatic and apical ROIs exhibited the same general trend: a strong positive USI when mismatch stimuli were first introduced, with a subsequent decrease over days (Fig. 2E vs. Fig. 2G). This global whole-cell mismatch response did not reflect the difference in morphology, inputs or potentially distinct computational roles of these compartments. A weaker version of the same trend appeared to hold in both somatic and apical dendritic ROIs in L5, but mean USIs were overall less significant. Mean USIs were significantly positive only on U1 in L5: for somata in VISp, VISl and VISpm, and for apical dendrites in VISp and VISl (Fig. 2F vs. Fig. 2H). This suggests that by U2, the responses in L5 were similar in both conditions and that there was less top-down feedback targeting L5 apical dendrites in the mismatch condition, particularly in VISpm and VISam.

We found that trends were also very similar across visual areas (Fig. 2E-H). In particular, in L2/3, all areas exhibited the strong initial response to mismatch stimuli on U1. Further, these initially positive USIs decreased systematically in all areas over days of exposure to the violation stimuli. This consistency in responses might suggest similarity in the information represented in these visual areas or a similar scalar modulation due to a higher-order error signal. In L5 we see weaker evidence for these trends, i.e., USIs were significantly positive on U1 for some areas/compartments. In area/compartment combinations with significantly positive USIs on U1, USIs also decrease significantly from U1 to U3.

For the natural movie stimuli, we found unexpectedly homogeneous mismatch responses in somata and apical dendrites, across visual areas, and to a weaker extent across cell lines. This is not in line with work predicting distinct mismatch responses in neural compartments targeted by top-down and bottom-up inputs or different levels of the visual hierarchy ^17,25^. Thus overall we found mismatch responses suggestive of predictive processing, but perhaps more distributed than some theories would predict ^2,3,57^.

### 2.4 Mismatch responses cannot be explained by other effects/factors

Attributing changes in responses to the mismatch condition requires carefully ruling out effects of potentially confounding stimulus properties or response history. In our case, novelty or stimulus differences cannot explain the difference in responses, since the stimuli in mismatch or control conditions were identical. Long-term adaptation (repetition suppression) could account for the reduction in mean responses over days, but would not account for the difference in responses to identical stimuli across conditions within a session ^58,59^. Further, it is unlikely that release from adaptation accounts for the elevated responses in the mismatch condition for the natural movies, since the reverse sequences in the mismatch condition repeated frames which were just presented. If anything, this would increase adaptation and decrease the responses in the mismatch condition. Finally, in both conditions, neural responses were very close in magnitude at the onset of the mismatch sequence (Fig. S1). Thus the difference is unlikely to be attributable to an initial difference in activity. The mismatch responses therefore do not appear to be driven by differences in the stimulus history, other than the initial sequence in the mismatch condition setting up specific expectations based on previous habituation.

Running speed has been shown to modulate neural activity in the visual cortex, as have changes in pupil diameter ^60,61,62,63,64,65,66^**^?^**. To determine how these behavioral effects contribute to our results, we compared running speed and pupil diameter across conditions. We found no significant differences between conditions (Fig. S4).

To verify that this elevation in responses was unique to the mismatch condition, we performed an additional control analysis consisting of a “same-same” comparison. The beginning of the fully reversed control and the reverse-forward “secondary control” are identical up to and including the mismatch sequence (Fig. S2A). They thus both contain stimulus segments that are identical to the reversed mismatch sequence, but in two different contexts which do not violate habituated expectations. Responses to that stimulus sequence in these two contexts were similar, without the increases seen in the comparisons above (Fig. S2B). Unlike Fig. 2C, average responses to these segments were consistently within error (Fig. S2C). This was also borne out in the calculation of “pseudo-USIs” for the “same-same” comparison (Fig. S2D), which were not significantly different from chance, with the exception of L2/3 apical dendrites on U1 (Fig. S2E-H). However, this single significant result, out of many comparisons after Bonferroni correction, may reflect residual session-level variance. We also observed significant changes across sessions, which appear broadly to align with adaptation effects.

A final potential confound in the mismatch natural movies is the reversal of visual flow and internal motion within the clips ^67^. Perhaps there could be a release from adaptation of motion or direction selective subpopulations due to the change in motion at the point of reversal ^68^. Against this interpretation, in a secondary control, we found responses to a forward reversal within a reversed movie sequence were reduced as compared to responses to the identical clip within the habituated movie (Fig. S3). This comparison lacks the habituated expectation cue required to create a mismatch condition, but includes the reversal of visual flow. The *decreased* activity in this case indicates that neural responses were more selective to the habituated forward sequence in its original context, and not elevated by reversal of visual flow, reversal motion, or change at violation onset. This further supports the interpretation that it is specifically the violation of habituated expectation (possibly including natural statistics of movies more broadly) which drives the elevated responses in the mismatch conditions for the natural movies.

Altogether, our results suggest that neurons in mouse visual cortex respond with increased activity to the same inputs when they violate expectations established by habituation and cued by the preceding sequence of stimuli.

### 2.5 Increased responses to reversals are driven by many cells, and show consistency across trials but not sessions

Given the evidence for elevated responses to the mismatch condition outlined above, we next investigated the distribution of these differences across the population, presentations of the stimulus, and sessions. Some studies suggest there may be specific error cells, or subpopulations with reliable positive or negative selectivity to mismatch stimuli ^6,7,8,25^, while others support a more distributed representation of mismatch responses ^16,17,18,23,38,44,45,69^. By examining the distribution of responses across ROIs we found that the positive population-level selectivity is driven by many cells, rather than a small subpopulation. Fig. 3A visualizes the proportion of ROIs with elevated responses to mismatch stimuli. The percentile of each ROI’s actual USI was computed relative to a null distribution of USIs (generated from shuffled control/mismatch labels), and resulting relative counts were plotted. Under the null hypothesis, where there is no systematic difference between responses to control vs. mismatch stimuli, this analysis would yield perfectly flat histograms, corresponding to uniform distributions of percentile values.

**Figure 3.**
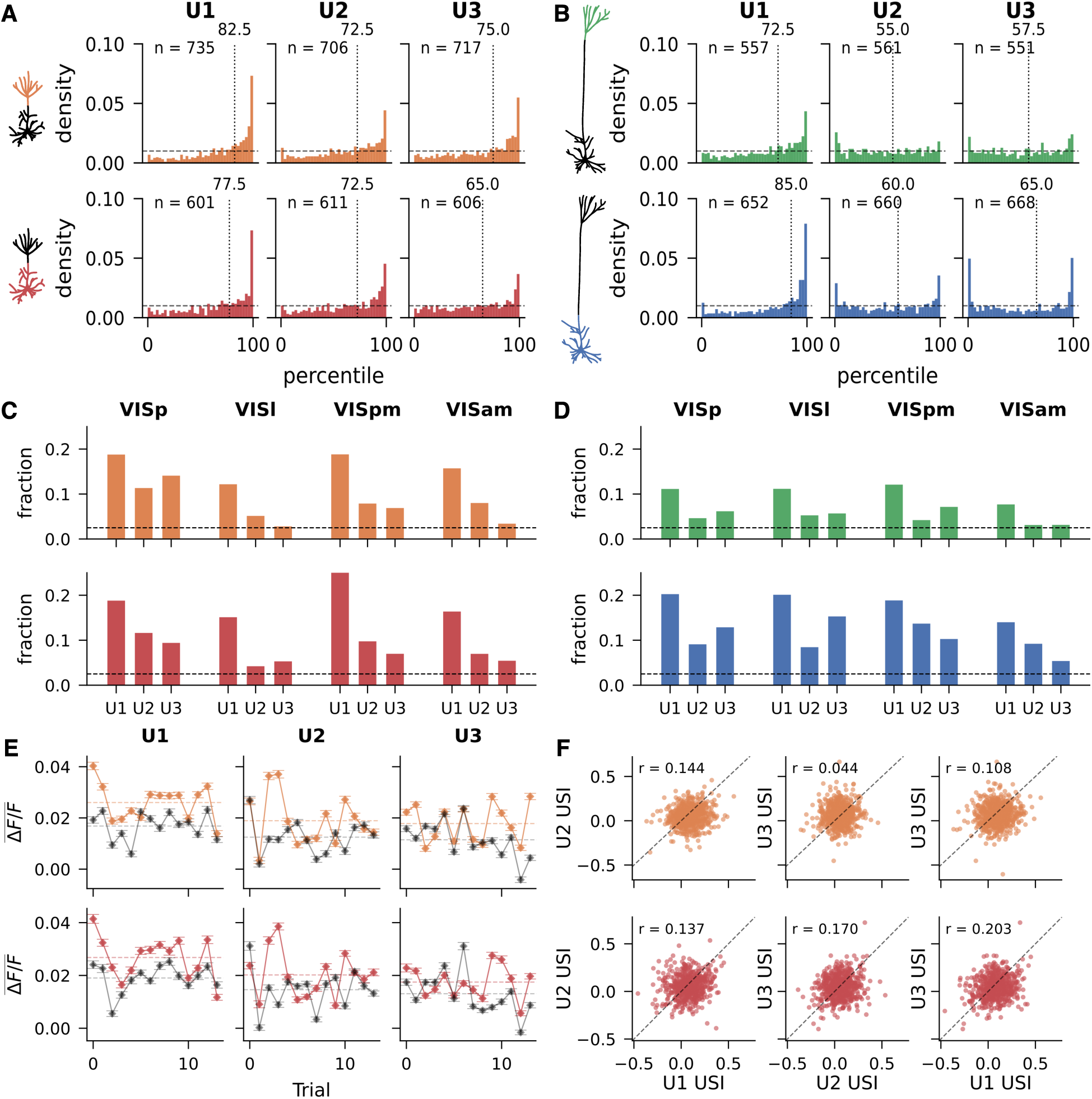
Many ROIs in VISp contribute to mismatch selectivity. **(A)**, **(B)** Distribution of ROI percentiles for ROIs in VISp for L2/3 and L5 cell line, respectively. Bars represent normalized proportion of ROIs whose USI fell within a given percentile bin of its shuffled distribution. Dashed horizontal line indicates the density expected in each bin for a uniform distribution, the expected density if ROI responses to the mismatch condition were no different from the control. Dashed vertical line marks the median percentile. Number of ROIs reported in top left. **(C)**, **(D)** Fraction of ROIs falling in the top-most bin (> 97.5 %) by percentile for the “worms” movie, across areas. This fraction is consistently above that predicted by chance (2.5 %; black dashed line) on U1, and decays to near the null line over sessions. **(E)** Mean response to the identical segment in mismatch and control conditions, broken down by trial index for ROIs in L2/3 of VISp. Colored series represent the mismatch condition, grey series represent the control condition. Points indicate the mean Δ*F/F* of each trial. Dashed horizontal lines mark the mean Δ*F/F* across all trials for each condition. **(F)** Scatterplots comparing USIs between sessions for ROIs tracked across days in L2/3 of VISp. Subplots are pairwise comparisons of the USIs across sessions, with points corresponding to individual ROIs. Dashed line marks y = x. Pearson correlation reported in top left. * *p <* 0.05, ** *p <* 0.01, *** *p <* 0.005.

Instead, in L2/3 VISp the density of USIs for natural movies was concentrated in higher percentiles (Fig. 3A), arguing against the null hypothesis, with the median initially lying above 75 % (and 18.8 % of both somatic and dendritic ROIs > 0.975). Over days, the median and density decreased, although even by U3 the number of ROIs with positive selectivity above the 97.5th percentile remains larger than expected by chance, and the median remains above 50 percent. Distributions for other areas can be found in Fig. S5. L5 ROIs exhibit a similar but much less pronounced effect (Fig. 3B), with far fewer statistically significantly high USI values, particularly in the apical dendrites. L5 neurons in VISp and VISl also showed mixed selectivity on U2 and U3, where the numbers of ROIs with positive and negative selectivity were both larger than expected by chance, particularly somatic ROIs (Fig. S5).

We next compared USIs for matched ROIs between each pair of sessions (Fig. 3F). USIs between sessions were somewhat correlated, but the relationship was very weak. That is, despite the systematic population-level trend (Fig. 2E, G) and the large number of ROIs with strong positive selectivity in all sessions (Fig. 3A), the actual subset of ROIs selective to mismatch stimuli varied substantially across sessions.

To identify the course of population mismatch responses within each session, we compared the mean responses to each control and mismatch presentation for one natural movie (worms). Although responses were most elevated in trials 1 and 2, we also found clear differences between stimulus conditions throughout U1 (Fig. 3E). Mean responses in each condition remain similar across trials, each fluctuating about distinct overall session means. Over days of exposure this difference in responses between conditions reduces, as reflected in the USI analysis above. However, we still see elevated responses to mismatch stimuli in the majority of trials, even in U3, rather than in just a few outlier trials with elevated responses.

Overall, we found that the observed mismatch responses were distributed across a broad population which changed between sessions, suggesting that they are not driven by a specific subpopulation or ‘error neurons’ with dedicated mismatch selectivity ^6,25^. Further, the timescale for learning driven by our mismatch stimuli is likely on the order of days rather than within minutes of exposure.

### 2.6 Gabor sequence stimuli do not elicit mismatch responses

We next investigated whether Gabor sequence stimuli elicited similar mismatch responses. Our Gabor stimuli were specifically designed to provoke such responses (see Methods 4.2.2), based on previous studies that found mismatch responses to similar sequences ^17,33^. We also incorporated certain controls for adaptation and effects of gross differences in stimulus, such as changing Gabor patch locations.

In contrast to the natural movies, we did not observe clear evidence for distinct responses to mismatch and control conditions in L2/3 for Gabor sequences. This is despite seeing clear onset and offset responses to these sequences, and changes aligned with the changing Gabor frames (Fig. 4B, C), both evidence that responses are driven by the stimulus. Repeating the USI analysis, we did not find significantly positive selectivity for the mismatch condition on U1, or the global trend in selectivity, decreasing from U1 to U3. We did, however, observe parallel between-session differences in both compartments and broadly similar between-session differences across visual areas.

**Figure 4.**
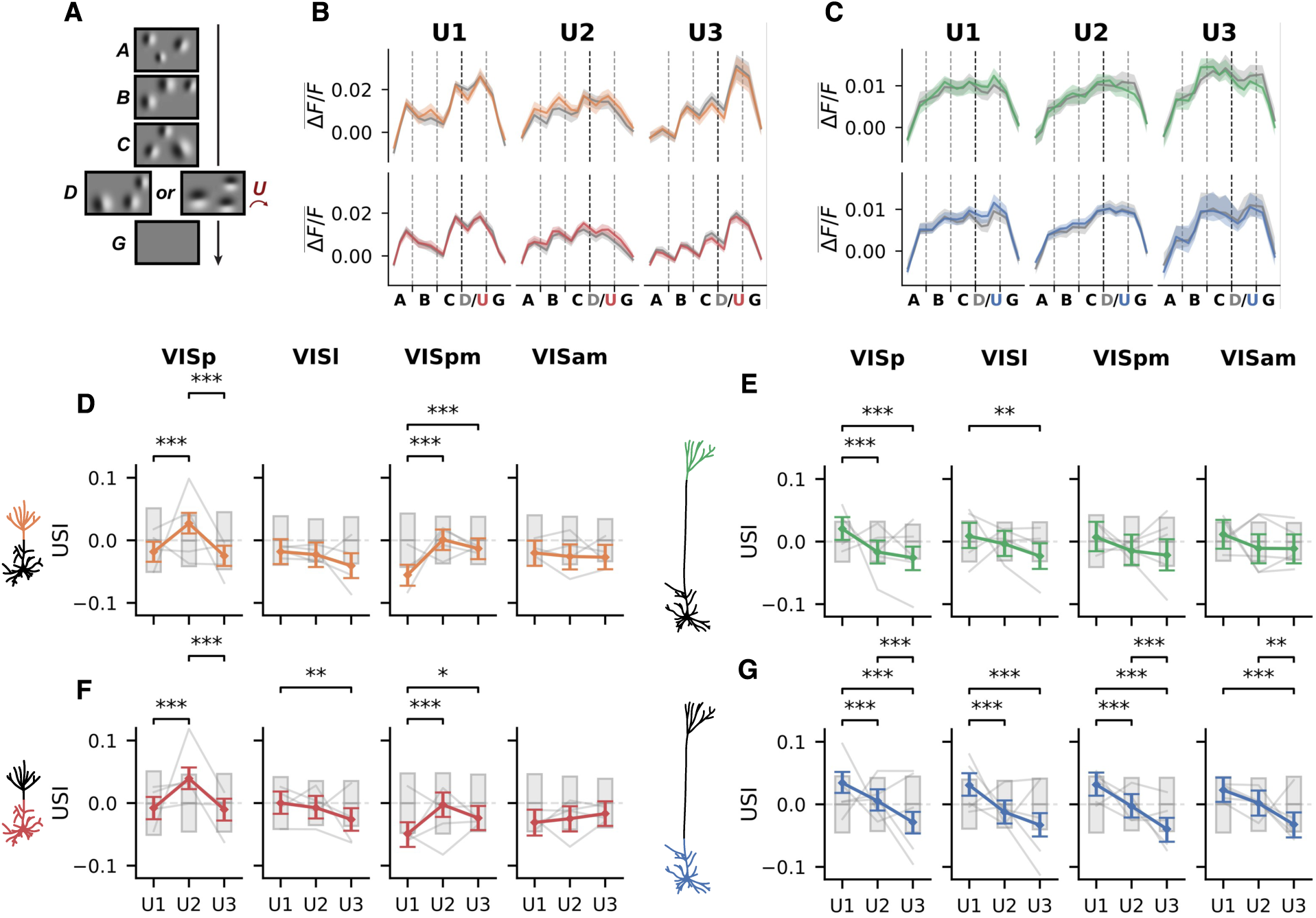
Analysis of responses in each condition for Gabor patch stimuli. **(A)** Structure of fixed Gabor stimuli. Subjects were habituated to sequences of frames A-B-C-D-G, with a common mean orientation. Unexpected sequences A-B-C-U-G were then introduced, in which the U frame had patches in the same location as D, but rotated by 90 degrees relative to the other frames in the sequence. **(B)**, **(C)** Mean Δ*F/F* across trials and ROIs for L2/3 and L5 respectively. Colored lines represent the mean response to the mismatch sequences (A-B-C-U-G), while grey represents the mean response to the control sequences (A-B-C-D-G). Vertical grey dashed lines demarcate the timepoints belonging to each frame in a sequence. Vertical black dashed line indicates the point of divergence in the two kinds of sequence. Colored bands represent a 95 % CI estimated using a hierarchical bootstrap. **(D)**-**(G)** USIs for sequences of Gabor patches with a fixed orientation across areas, sessions, and cell lines. Grey bars represent 95% CI for a permutation based null distribution. Error bars represent a 95 % CI estimated using a hierarchical bootstrap. Fine grey lines show average individual subject USIs. * *p <* 0.05, ** *p <* 0.01, *** *p <* 0.005.

In L5, we found evidence for selectivity decreasing significantly over days of exposure to pattern-violating stimuli (Fig. 4E, G). The systematic decrease in USIs is particularly striking, given the consistency across compartments and areas. However, selectivity to mismatch stimuli was never itself significant. Furthermore, the same analysis applied to the rotating sequences of Gabors did not replicate this trend (Fig. S7), and instead showed a trend in the opposite direction (but also without significant selectivity).

Separately, we examined the responses of orientation selective cells (Fig. 5). A concern with the population level analysis is that a small effect, or effects of opposite sign, might be lost amid random variation in non-condition-selective cells. Given the structure of our stimuli, orientation selective cells are especially likely to be causally implicated in the processing of particular frames, and might therefore be the cells in which our Gabor stimuli induce mismatch responses.

**Figure 5.**
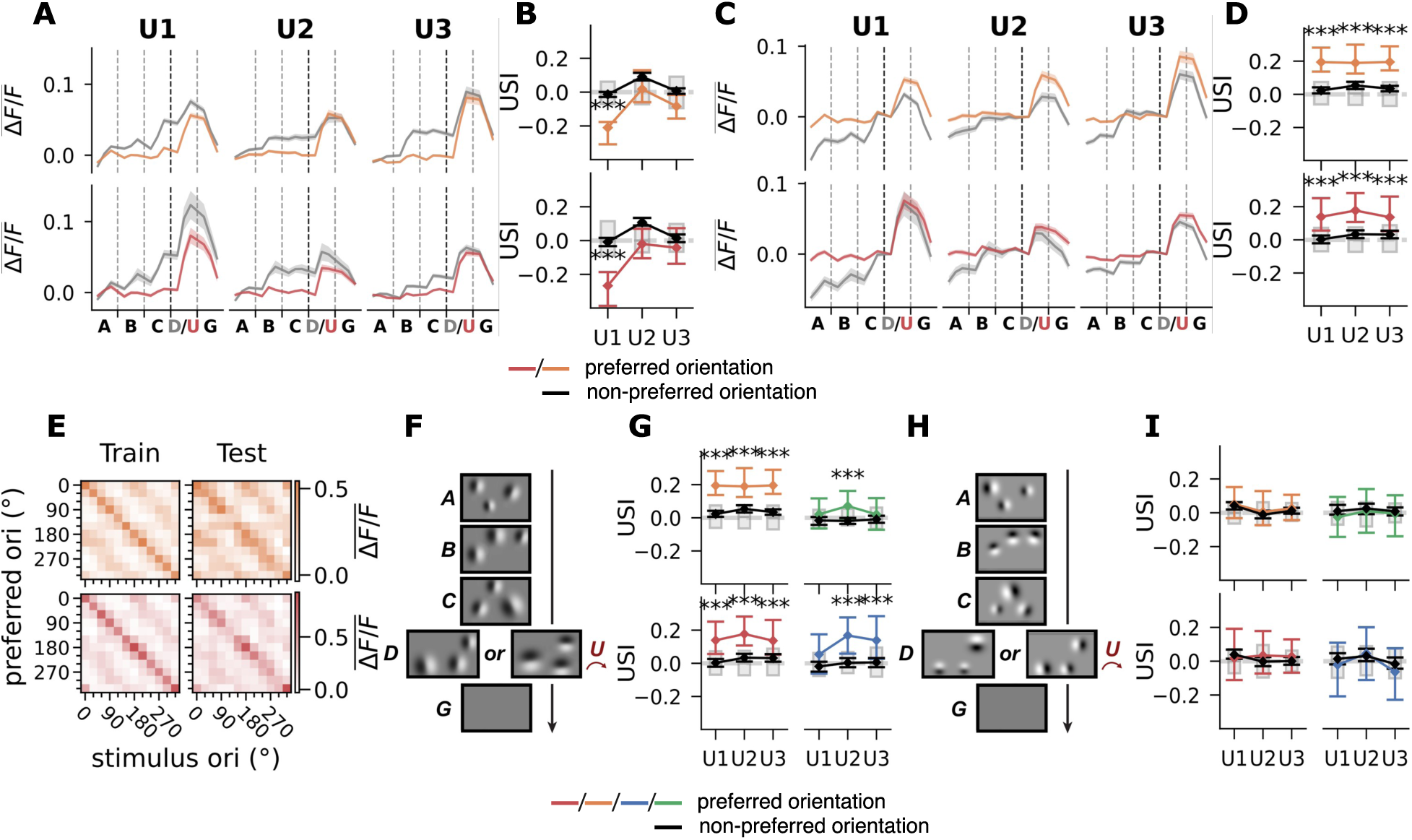
Responses to Gabor stimuli restricted to orientation-selective cells. **(A)** Mean responses for orientation selective cells to A-B-C-D-G and A-B-C-U-G sequences in which D and U matched their preferred orientation. **(B)** USIs broken down with orientation selectivity. Colored lines are USIs calculated for orientation selective cells to trials in which D/U matched their preferred orientation. Black lines are USIs for the same ROIs to trials not matching their preferred orientation. **(C)** Same as (A), but after subtracting the Δ*F/F* value at the first time point of D/U on each trial, from that trial’s Δ*F/F* time series. **(D)** USIs computed after the same subtraction. **(E)** Response heatmaps for orientation selective cells. Control sequences were split into disjoint training and test sets of equal size. These were further chosen so that the test set contained a subset of sequences matched to the mismatch sequences in D/U-frame orientation, which was used in USI analyses. Train heatmaps show the mean Δ*F/F* across trials and timepoints for each stimulus orientation, for cells identified as selective to each preferred orientation. Test heatmaps show the same, but for held-out test trials. Diagonal bands can be seen in both, consistent with selectivity to the preferred orientation at each angle. See Methods 4.3.3 for further analysis details. **(F)** Example of fixed Gabor sequence stimulus. **(G)** USIs broken down with orientation selectivity for L2/3 and L5 ROIs from VISp, for fixed Gabor stimuli, after subtraction of initial time point magnitude. Color-coding as in (B). Orientation selective cells show large positive USIs for their preferred orientation (colored), but not for other orientations (black). **(H)** Example of rotating Gabor sequence stimulus. **(I)** USIs broken down by orientation selectivity for L2/3 and L5 ROIs from VISp, for rotating Gabor stimuli, after subtraction of initial time point magnitude. Orientation selective cells no longer show strongly positive USIs for their preferred orientation, which are now near zero, similar to their non-preferred orientation and non-selective cells. * *p <* 0.05, ** *p <* 0.01, *** *p <* 0.005.

Specifically, since these cells are active in response to particular orientations, they will respond selectively to particular D frames in a way that can be anticipated based on the orientations of the earlier frames in the sequence. Further, if higher order predictions are cued by feedforward signals, based on the selectivity of these neurons to the orientations of the ABC sequences, the top-down predictive signals related to D/U orientations would likely target these same cells ^70^. Thus, they are more likely to have their responses modulated by top-down feedback.

We found two competing effects: the mean Δ*F/F* at each frame ramps over the course of control sequences, in which patches with the preferred orientation are presented repeatedly in different locations, but the change in mean Δ*F/F* from frame C to U is greater in mismatch sequences than the change from frame C to D in control sequences (Fig. 5A, B). We subsequently corrected for the fact that the calcium Δ*F/F* signal does not have time to return to zero at the first time point of the D/U frames. Specifically, the baseline Δ*F/F* at the first time point of the D/U frame on each trial was subtracted from that trial’s Δ*F/F* time series (Fig. 5C, D).

With this baseline correction, we found evidence for an adaptation effect, but not for mismatch responses in orientation selective cells. USIs for the corrected fixed Gabor sequences were strongly positive, reflecting the diminished change in response at frame D to the repeated presentation of the same orientation relative to the U frame, which is the first presentation of the preferred orientation within the mismatch trial (Fig. 5F, G). For rotating Gabor sequences, in which the preferred orientation was not repeated in the control condition, and repeated only once in the mismatch condition, the USIs were instead near zero (Fig. 5H, I). This strongly suggests that the difference for the fixed Gabor sequences was due to adaptation, not mismatch. It is also worth noting that we did not see these adaptation effects reflected in the population level means (Fig. 4D-G), supporting the earlier suspicion that small effects might be washed out.

Overall, our results suggest that these Gabor sequences do not elicit significant mismatch responses in L2/3 or L5. This is inconsistent with previous work using similar stimuli, although lacking some of our controls ^15,17,33^. On the other hand, this may be consistent with recent work which proposes that apparent mismatch responses to gratings-type stimuli may actually be an adaptation effect ^19^, especially given our findings for orientation selective cells. It is also consistent with recent work suggesting simple artificial stimuli do not effectively elicit mismatch responses ^21,22,71^. Our negative results for Gabor sequences also contrast with the clear mismatch responses to reversals in natural movies we observed. Taken together, the natural movies and Gabor results seem to suggest that naturalistic stimuli may have different effects/salience from artificial stimuli, which may contribute to mismatch responses.

### 2.7 Somato-dendritic coupling is unchanged by mismatch stimuli

Another principal aim of our study was to characterize the degree of coupling between somata and connected distal apical dendrites, and how coupling varied across stimulus conditions. Many theories of cortical processing suggest either coactivation or comparatively independent activity in apical dendrites and somata during periods of mismatch between top-down expectations and bottom-up inputs ^17,25,43^. Apical inputs can robustly drive neuronal firing when they coincide with perisomatic inputs, and apical plateau potentials can generate high-frequency bursts that are thought to promote synaptic plasticity ^36,37^. It has also been suggested that mismatch between predicted and actual somatic activity may be computed within the apical dendrites and exploit local disinhibition in the apical arbor ^11,12,14,43^.

To study somato-dendritic coupling, we first matched apical dendritic and somatic ROIs to single cells using tracings of z-stacks assembled from in vivo neurophysiological recordings (Fig. 6A (i); see also Methods 4.4). Reconstructed neurons in our z-stacks were matched to individual distal apical dendrite ROIs from one recording plane to their source somata ROIs in the other plane (Fig. 6A (ii-iii)). Matched compartments exhibited both independent and coupled activity in somatic and dendritic compartments (Fig. 6B). Many matched pairs were correlated overall, although with considerable variance in coactivation for smaller ΔF/F (Fig. 6C). As a control, we shuffled our matched pairs to create a set of deliberately mismatched pairs. The matched pairs showed systematically higher Pearson correlation, compared to shuffled pairs (Fig. 6D). For further validation of the matching process see Methods 4.4.2.

**Figure 6.**
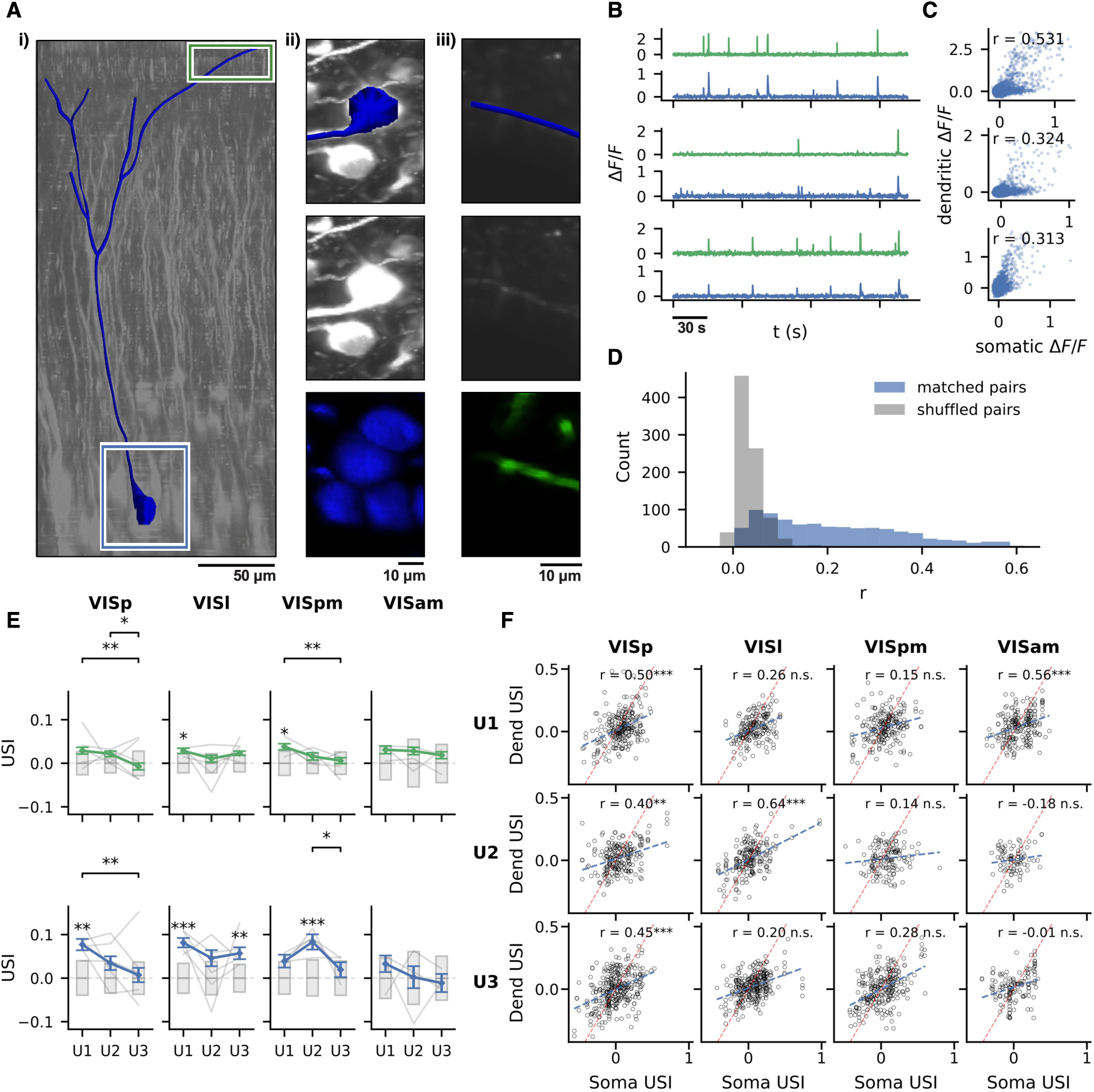
Somato-dendritic coupling. **(A)** Examples of ROIs matched to a traced cell. **(i)** Example tracing of an L5 pyramidal neuron. **(ii)** From top to bottom: the traced soma, the soma as it appears in the z-stack, the corresponding ROI shaded in blue (accompanied by nearby ROIs). **(iii)** The same as (ii) for a dendritic ROI matched to the same cell, with position indicated in (i). **(B)** Example time traces for somatic and dendritic ROIs matched using tracings. Green corresponds to dendritic Δ*F/F*, blue corresponds to somatic. **(C)** Scatterplots of dendritic vs somatic Δ*F/F* across a full imaging session for the same pairs as in (B), with Pearson correlation. **(D)** Histograms of Pearson correlations for time traces of pairs of somatic and dendritic ROIs. Blue corresponds to actual matches identified from tracings. Grey corresponds to an equal sized null sample constructed by randomly picking either the somatic or dendritic ROI from each actual match, and pairing it with a randomly selected ROI from the other plane. **(E)** USIs for natural movies for connected soma-dendrite pairs. Grey bars represent 95% CI for a permutation based null distribution. Error bars are the SEM over ROIs. Fine grey lines show average individual subject USIs. **(F)** Comparison of USI between connected soma-dendrite pairs, with correlation between compartmental USIs shown. * *p <* 0.05, ** *p <* 0.01, *** *p <* 0.005.

We repeated the USI mismatch selectivity analysis for natural movies, restricted to somatic and dendritic ROIs with a confirmed anatomical connection. The trends were broadly similar to the L5 population trends in Fig. 2F, H, with larger USIs on U1 which decreased toward zero over subsequent sessions, although in the restricted comparison this decrease across sessions was only significant in VISp and VISpm (Fig. 6E). On U1, somatic USIs were significantly positive in VISp and VISl, while apical dendritic USIs were significant only in VISl and VISpm. VISam once again never exhibited significant mismatch selectivity in either compartment. Two slight deviations were found, with VISl showing reduced but still significant selectivity on U3 and VISpm showing significant selectivity only on U2, that became negligible on U3. Examining the correlation between the USIs (i.e., the similarity in mismatch responses) of connected distal apical dendrites and somata, we found strong correlation in VISp for all sessions, as well as in VISl (U2) and VISam (U1) (Fig. 6F). This matches the previously observed population trend of broadly parallel responses across compartments (Fig. 2F, H).

As an initial measure of coupling between somatic and dendritic activity in a given neuron, we computed the Pearson correlation of their respective Δ*F/F* time series (Methods 4.3.4). We found similar somato-dendritic correlation during predictable and unpredictable stimulus sequences, suggesting the coupling of somatic and dendritic activity was not modulated by mismatch stimuli (Fig. S11). Responses to all natural movies reflected this independence of coupling to violations of habituated sequences.

Large transient elevations in the Δ*F/F* signal are the clearest indication of depolarization in our recordings (as can be seen in Fig. 6B). Such large transients have been shown to correlate with high-frequency spikes or bursting in somata and back-propagating action potentials, spikes or plateau potentials in dendrites, and hence they represent the most reliable/obvious indicator of elevated neural activity ^72,73^. We isolated these transients from the Δ*F/F* time series for each ROI (Methods 4.3.4), and defined coincident or coupled transients as those occurring within 300 ms of each other (Fig. 7A). In all neurons, we observed instances of simultaneous transients in both compartments and independent activity in both dendrites and soma, i.e., without significant coactivation of connected compartments (Fig. 7C).

**Figure 7.**
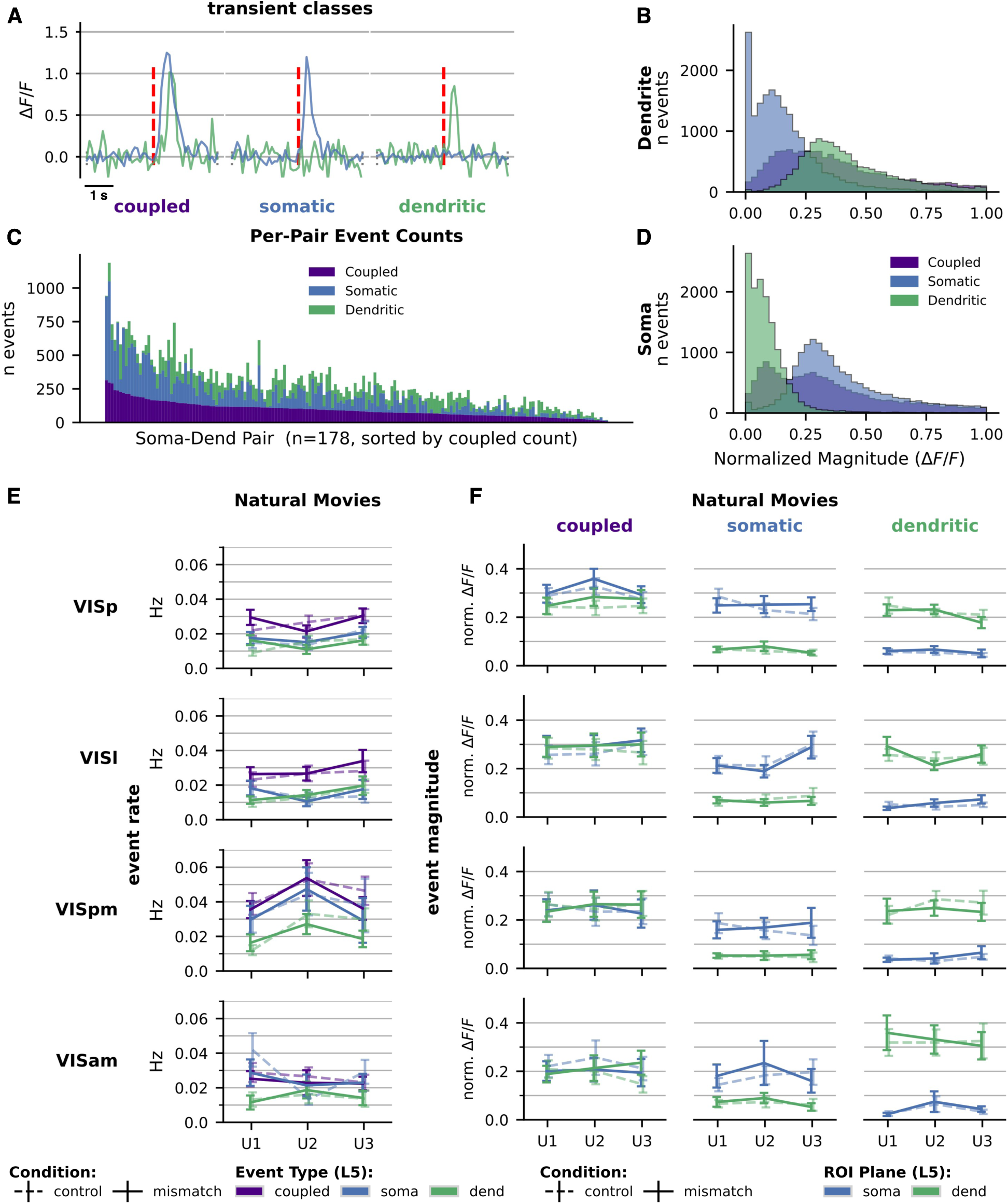
Analysis of somato-dendritic coupling using Δ*F/F* transients for L5 connected soma-dendrite pairs identified by tracing in volumetric imaging. **(A)** Comparison of transient classes for a connected soma and dendrite pair. **(B)** Histogram of normalized magnitude of transients per class, for dendritic ROIs in VISp. **(C)** Stacked histogram showing the per pair transient event counts in VISp. **(D)** Same as (B) for somatic ROIs in VISp. **(E)** Rates of transients for each class, averaged across connected pairs, for natural movies. **(F)** Normalized Δ*F/F* magnitudes of transients for each class, averaged across connected pairs, for natural movies. * *p <* 0.05, ** *p <* 0.01, *** *p <* 0.005. See Table S2 for the number of pairs in each comparison and Fig. S12 for similar Gabor sequence comparisons. See Table S1 for number of connected pairs per area.

We investigated whether these transients varied across stimulus conditions in several ways. First, we compared their rates, whether coupled across a given neuron or decoupled within one of its compartments. For natural movies, we found they occurred with similar frequency across stimulus conditions (Fig. 7E). We also compared the magnitudes of the three classes of transients, but again for natural movies found that transient magnitudes were not modulated by stimulus condition (Fig. 7F). Results were similarly negative for Gabor sequences, again with a single exception for rotate Gabor sequences where dendritic transients were elevated during control sequences on U2 in VISam, although this difference was driven by a few transients from two dendritic ROIs (see Fig. S12).

To extend this comparison to more of our ROIs and to L2/3 subjects for which we did not have matches from z-stack tracings, we conducted additional analyses using pairs matched based on coincident activity. Matched pairs were identified using a magnitude-weighted overlap of the largest transients between somatic and dendritic ROIs (Methods 4.5). These matches warrant much lower confidence than the anatomically connected pairs we identified by tracing our volumetric z-stacks, and this matching process biases any subsequent analysis towards ROI pairs with higher coupling.

For natural movies the rates of transients, whether coupled across a given neuron or decoupled within one of its compartments, occurred with similar frequency for mismatch and control stimuli in both L5 and L2/3, with the exception of the coupled class on U1 in L5 VISp (Fig. S13A, C). This single outlier among the 48 comparisons for L5 does show a higher rate of coupled transients during the mismatch condition for natural movies, but it is contradicted by the more reliable equivalent comparison based on anatomical matching (Fig. 7E). Similarly, for natural movies the magnitudes of the three classes of transients were not significantly modulated by stimulus condition. This was true across all visual areas and both cell lines, with the exception of minute responses in L2/3 somata during dendritic transients in the control condition on U3 in VISp (Fig. S13B, D). Results were similarly negative for Gabor sequences, with no significant differences in the rates or magnitudes of transients in mismatch versus control conditions (Fig. S14, Fig. S15). Overall, the mismatch responses we observed during natural movies (Fig. 2E-H) did not correspond to, and thus cannot be explained by changes in coupling (in L2/3, Fig. S13A,B or in L5, Fig. 7E,F).

Thus, despite finding significant selectivity to mismatch conditions for the natural movies (Figs. 2-3), we found that this selectivity was not reflected by changes in somato-dendritic coupling. Research related to coincident inputs driving bursting ^37,39^ predicts weakened coupling during mismatch stimuli, whereas theories centered on the segregation of dendrites by chronic inhibition predict strengthened coupling ^11,36^. Our results do not support either of these predictions, as we did not find evidence for changes in the coactivation of somatic and dendritic compartments during mismatch responses.

## 3 Discussion

Reversals of habituated natural movie segments elicited robust mismatch responses in both somatic and distal apical dendritic compartments across all recorded visual areas. Purely sensory sequence predictions, without closed-loop motor feedback, appear sufficient to drive cortical expectation-violation signals. The lack of similar responses to Gabor sequences, suggests that the continuous spatiotemporal structure of natural scenes is critical to such expectation-based modulation. Despite clear compartmental mismatch responses, somato-dendritic coupling was unchanged. This constrains the potential contribution of single-neuron coincidence-detection mechanisms to purely visual prediction-error signaling.

We observed clearly elevated mismatch responses to unexpected reversals of natural movie stimuli (Results 2.2 & 2.3). These responses could not be explained by adaptation to repeated stimuli, since the reverse sequences in the mismatch condition repeated frames which were just presented. We used responses to the fully reversed movie as a control to be sure that the increase in responses was not simply due to the novelty of the stimulus or the basic selectivity of the cells being recorded. In line with predictive processing theories ^2,4,74^, the enhanced responses to the reversals were strongest on the first day of exposure, when the reversals were most unpredictable, and decreased over multiple days. Long-term adaptation across sessions could account for a reduction in mean responses over days ^58,59^, but not for the elevated mismatch response relative to control responses within the same session, given the equivalent prior exposure history.

These mismatch responses were strongest in L2/3 neurons of VISp, but also present in other regions and in L5. The stronger and more sustained responses in L2/3 are consistent with the central role of L2/3 posited by some predictive processing theories, specifically L2/3 as the principal source of feedforward error signals to higher cortical areas ^34,75^. Previous studies have similarly found stronger visual sensorimotor mismatch responses in L2/3^24,25,38^. Although responses in L2/3 were stronger, we also found mismatch responses in L5 somata, particularly on U1 and decreasing with subsequent exposure. The weaker and less sustained L5 response may reflect the dissociation between visual perception and action. Further, in a passive, unrewarded paradigm, the drive from the lateral posterior (LP) thalamus to L5 apical compartments is intermittent, reducing L5 mismatch sensitivity relative to L2/3^38,76,77,78^.

The trend in mismatch responses was strikingly parallel across visual areas. Based on some theories of hierarchical visual processing we might have expected greater differences between these areas ^2,3,20,44,45^. The systematic downward trend in mismatch responses we found is most consistent with a common modulatory broadcast reaching all areas simultaneously, a scalar gain-like modulation ^23^, rather than the vectorized modulation predicted by a locally computed hierarchy of prediction errors.

Finally, when we examined the responses in the apical dendrites versus the somata we did not observe significant differences in the responses of the two compartments. We also did not observe changes in apical events or somato-dendritic coupling in response to mismatch stimuli, even when we restricted our analysis to compartments that had identifiable connections within our volumetric z-stack images. These results do not support simple theories of sensory prediction error calculations in the apical dendrites ^10,11,12,14,43^.

In contrast to the mismatch responses we observed with natural movies, we did not find strong evidence for mismatch activity in response to sequences of random Gabor patches. At a population level, we did not see clear evidence for a general increase in activity when the orientation of the Gabor patches was unexpected. When we targeted our analysis to cells that were selective for the unexpected orientations we did see some evidence for an increased response to a new orientation following a sequence of repeats, but this response could easily be explained by adaptation effects, not prediction violations. This interpretation is further supported by the lack of evidence for increased responses to an unexpected repeat of orientations following a sequence of rotations.

The results for natural movies are broadly consistent with the idea that the neocortex generates predictions about upcoming stimuli based on prior experience, and can update these predictions with enough new data. The presence of mismatch responses to natural movies and absence of such responses to artificial Gabor sequences supports the view that the visual system is tuned to natural statistics, and that naturalistic stimuli are likely critical to generating sensory mismatch responses. Supporting this assertion, a recent study also found greater mismatch responses in L2/3 in the mouse visual cortex, specifically for natural but not artificial stimuli ^22^.

Our intention, in designing our Gabor stimuli, was to minimally modify the stimuli used in a previous study ^17^, in order to reproduce those results, but with the benefit of our simultaneous imaging of compartments and imaging in multiple areas. However, we did introduce a critical modification to the Gabor sequences in the present study. For the mismatch frame, we changed only the orientation (removing the change in location), so that the patch positions were the same as in the control condition. This ensured that any mismatch responses were driven only by changes in orientation, and mitigated potential release from adaptation due to changing positions. However, although the mice were head-fixed, eye movements could also change which neurons were activated by a given Gabor frame. Ultimately, this study did not reproduce the prior results ^17^ for VISp, in particular the different trends in USIs in somatic and dendritic compartments. It is possible that this lack of replication was a result of the changes we made to the stimuli. Alternatively, it could be that the lack of simultaneous imaging in the previous study was key to those results.

There are several limitations to our study that should be recognized. First, because the mice were not being rewarded in any way, they may not have been engaged by the stimuli and may not have been actively attending to them ^32^. This could have reduced the effects of prediction violations. On the other hand, this experimental design avoided confounds due to reward prediction errors, thereby helping us to isolate the effects of sensory prediction errors ^79,80^. Second, we observed changes in the responses over days that suggested that the neurons were learning about the mismatch stimuli and incorporating them into an internal predictive model. However, we cannot say with certainty what types of changes were occurring in the circuit at a mechanistic level. As well, the temporal resolution of our imaging made it impossible to identify the temporal order of responses across planes (i.e., across compartments or between areas). This made it challenging to draw conclusions about the relative timing of top-down or bottom-up signals.

Overall, our findings suggest that the neocortex implements predictive processing that can be driven by purely sensory mismatch. They are also consistent with previous findings suggesting that visual systems are tuned or more sensitive to natural statistics and natural sequences ^21,22,29,30,32,71,81,82^. Future work should unpack the specific microcircuit mechanisms responsible for generating predictions and detecting mismatches.

## 4 Detailed Methods

### 4.1 Experimental animals and calcium imaging

The dataset analyzed in this study was obtained through the Allen Institute for Brain Science OpenScope initiative. All animal procedures were reviewed and approved by the Institutional Animal Care and Use Committee (IACUC) at the Allen Institute for Brain Science.

Two transgenic mouse lines were used to selectively express GCaMP6f in pyramidal neurons within chosen cortical layers: Cux2-CreERT2;Camk2a-tTA;Ai93 (layer 2/3 pyramidal neurons) and Rbp4-Cre KL_100;Camk2a-tTA;Ai93 (layer 5 pyramidal neurons). Mice underwent cranial window implantation surgery and were housed individually on a reverse light–dark cycle. All experiments were conducted during the dark phase. Additional details regarding Cre lines, surgical procedures, habituation, and quality control are provided in ^42,83^.

After 1–2 weeks of post-surgical recovery, animals underwent intrinsic signal imaging for retinotopic mapping and segmentation of target visual areas. This was followed by a two-week habituation period. During week 1, mice were habituated to handling, head fixation, and running on a disc. In week 2, they were exposed to visual habituation stimuli for progressively increasing durations (10–50 minutes).

Imaging sessions proceeded as follows. First, two sessions (*∼*70 minutes each) were devoted to z-stack acquisition while habituation stimuli were presented. Next, three 70-minute dual-plane imaging sessions were conducted with habituation stimuli, followed by three additional 70-minute dual-plane sessions which included mismatch stimuli. Imaging was performed in four visual areas during these sessions.

Multi-plane calcium imaging was carried out using a Dual-Beam Mesoscope (Multiscope) ^84^. Laser excitation at 910 nm was provided by a Ti:Sapphire laser (Chameleon Ultra II, Coherent). The excitation beam was split into two paths, with one beam delayed by half the laser’s excitation period (6.25 ns), enabling multiplexed simultaneous imaging from two focal planes (e.g., L1 20 µm and L5 375 µm depth, or L1 50–75 µm and L2/3 175 µm depth). Visual areas were sampled sequentially (VISp, VISl, VISpm, VISam), with resonant scanners over a 400 µm field of view at 512 × 512 pixel resolution, yielding a sampling rate of 10 Hz per plane.

Sessions that failed quality control were excluded, resulting in a final sample of nine mice, all with six high-quality experimental sessions (E1-E3 and U1-U3): four L2/3 mice (all male) and five L5 mice (two male, three female).

Neuronal recordings were motion-corrected using Suite2p and denoised using DeepInterpolation ^85,86^. ROI masks were segmented from denoised movies using the robust estimation algorithm described in ^41,87^. Fluorescence time series from somatic and dendritic ROIs were extracted, neuropil-subtracted, demixed, and converted to Δ*F/F* signals from the raw calcium movies using the Allen Software Development Kit (SDK), as described in ^83,88^. Neuropil correction, combined with a 180-second sliding window to compute a rolling baseline fluorescence (F), ensured that Δ*F/F* traces were robust to variations in background fluorescence across mice and imaging planes.

ROIs identified as duplicates or unions, those overlapping motion borders, or those deemed excessively noisy were excluded from further analysis ^83,88^. After filtering: VISp 232–948 dendritic and 146–664 somatic ROIs, VISl 315–952 dendritic and 118–580 somatic ROIs, VISpm 242–953 dendritic and 32–598 somatic ROIs, VISam 333–958 dendritic and 69–543 somatic ROIs per mouse per session were retained. The complete dataset is publicly accessible in Neurodata Without Borders (NWB) format through the DANDI Archive ^89,90^. ROIs were matched across days using a customized version of the Allen Institute’s n-way matching package.

Typical Z-stack acquisition consisted of a *∼* 70 minute recording session, during which habituation stimuli were presented. The field of view was extended to 500 µm with a resolution of 768 x 768 pixels. This allowed for extra coverage outside each edge of the imaging planes from each experimental session. 100-frame sequences were acquired at each depth, starting at the brain surface and stepping downwards by *∼*1 µm between planes. For L2/3, four stacks were acquired per session, over depths of 0-225 µm. For L5, two dendritic stacks were acquired (0-200 µm) and one somatic stack (0-450 µm). At least one set of stacks was acquired per visual area, with the Multiscope facilitating the acquisition of dual stacks for adjacent areas in some cases (e.g., VISam and VISpm). If signal-to-noise was low, z-stack acquisition was repeated after all experimental sessions were complete.

### 4.2 Visual mismatch stimuli

During each habituation and imaging session, mice viewed both the natural movies and Gabor sequences. The two stimuli were presented in consecutive blocks, in random order, for equal durations. All stimuli were projected onto a flat 24-inch monitor positioned 10 cm from the right eye. The monitor was rotated and tilted to be perpendicular to the eye’s optic axis, and the stimuli were spatially warped to approximate presentation on a spherical projection screen.

Pre-recording habituation sessions increased in duration over successive days, ranging from 10 to 50 minutes. In contrast, the z-stack acquisition and optical imaging sessions were *∼* 70 minutes and included *∼* 30 minutes of the natural movies and *∼* 30 minutes of Gabor sequences. Stimulus blocks were flanked by gray screen periods lasting 30 seconds. Each session ended with two additional control stimuli, static and moving gratings with the same spatial frequency as the Gabor patches, presented for 90 seconds each.

Critically, each natural movie was presented 450 times over a period of 2 weeks prior to the introduction of mismatch movies (60 times each for a full-length E-day habituation session). Thus all subjects had substantial long-term exposure to each stimulus. During the final 3 imaging sessions, each natural movie presentation consisted of 15 habituated sequences (fwd), 15 mismatch (fwd-rev), and 15 of each of two controls (rev and rev-fwd), in random order. For Gabor sequences, each sequence type was presented 4500 times over 2 weeks prior to the introduction of mismatch sequences (600 times each for a full length E-day habituation session). During the final 3 imaging sessions, 20% of sequences include mismatch U frames (i.e. 480 habituated sequences and 120 mismatch sequences) for each sequence type.

#### 4.2.1 Natural movies

We created three types of natural movie stimuli, based on three different sources: ‘Worms’, a video of dozens of worms wriggling in soil (i.e., a prey stimulus), ‘Flowers’, a video of two wild mice running and climbing outdoors (i.e., a conspecific stimulus), and ‘Touch of Evil’, the motion picture. 9 seconds containing dynamic action were edited from each of these sources to form a base natural movie. Each frame was then normalized and matched for mean brightness to our gray screen stimuli. To help mitigate adaptation effects to fixed objects within each movie, we added random motion. This was achieved by cropping each frame of the original videos, while adding a smooth random trajectory to this field of view across the frames, so that each was cropped differently. These modified, mean-corrected 9-second clips formed our habituation stimuli. We dubbed them forward (fwd), to distinguish them (i.e., their ordinary temporal order) from our mismatch and control stimuli.

Mismatch natural movies were created from these forward clips by adding a short reversal in the order of frames. Approximately 6 seconds into the habituated natural movie, the sequence reversed for 3 seconds (i.e., a sequence a-b-c-d-e-f-g-h-i became a-b-c-d-e-f-f-e-d). Thus the mismatch was created by a subsection of the habituation movie being played back in reverse. These mismatch natural movies, with 6 seconds identical to habituated movies and 3 seconds reversed, were dubbed forward-reverse (fwd-rev).

Two additional controls were created. The reverse (rev) control consisted of the identical frames as in our habituation movies, in reverse order. The reverse-forward control (rev-fwd) was equivalent to making a pseudo-mismatch stimulus from our reverse control movies, with 6 seconds identical to reverse movies and 3 seconds reversed (i.e., presented in correct temporal order, matching 3 seconds from the habituated movies.)

Critically this set of controls provided two points of comparison for the reversed mismatch sequences in fwd-rev movies (i.e., from both rev and rev-fwd sequences). In both cases, the mismatch sequences occur in the context of an unhabituated reversed sequence. They both lack the context of the preceding habituated sequence intended to create expectation violation or mismatch effects.

#### 4.2.2 Gabor sequences

The Gabor sequence stimuli were adapted from that used in two previous studies ^17,33^. They both consisted of repeating 1.5-second sequences composed of five consecutive 300-ms frames (A-B-C-D-G). Frames A, B, C, and D each contained 30 Gabor patches defined by specific locations and sizes, whereas frame G was uniformly gray.

For ‘Fixed Gabor’ sequences, within each session the locations and sizes of the Gabor patches were uniquely fixed for a given frame type (e.g., all A frames were identical in layout, but differed from B, C and D frames). These spatial configurations were resampled between mice and between days (i.e., both Gabor patch size and location), such that no two sessions, even for the same mouse, contained identical Gabor sequences. Patch locations were sampled uniformly across the visual field, and patch sizes were sampled uniformly between 10° and 20° of visual angle. Thus, within a given session, the Fixed Gabor sequences were defined by the characteristic locations of the Gabors for each A-B-C-D frame.

Within each sequence (A-B-C-D-G), Gabor orientations were drawn randomly from a von Mises distribution with concentration parameter *κ* = 16 and a shared mean. The mean orientation was randomly selected for each sequence and counterbalanced across 12 values (0°, 30°, 60°, … 330°). Consequently, although a wide range of orientations was presented across a session, orientations within an individual sequence were tightly clustered.

Mismatch Fixed Gabor sequences were generated by replacing the D frame with a U frame (A-B-C-U-G). U frames differed from D frames only by orientation statistics: their orientations were sampled from a von Mises distribution with a mean shifted by 90° relative to the preceding regular frames (A-B-C), depending on the original mean. Unlike some previous studies ^15,17,33^, we did not change the spatial layout (distinct patch locations and sizes). This control was added to avoid effects due to release from adaptation or driven by large changes in spatial distribution of stimuli. Thus the mismatch was due to the unexpected orthogonal shift in orientation alone.

For ‘Rotate Gabor’ sequences, the A-B-C-D sequences were composed identically to the fixed Gabors in terms of the locations and sizes of the Gabor patches. Patch locations were similarly sampled uniformly across the visual field, and patch sizes were sampled uniformly between 10° and 20° of visual angle.

Within each Rotate Gabor sequence orientations were again drawn randomly from a von Mises distribution with concentration parameter *κ* = 16, but unlike the Fixed Gabor sequences, the mean was not shared within a given sequence. The mean orientation of the A frame was similarly randomly selected for each sequence and counterbalanced across 12 values (0°, 30°, 60°, … 330°). However, the mean orientation of subsequent frames shifted by 90° (e.g., a mean of 0° for A, 90° for B, 180° for C, and 270° for D). Thus the Rotate Gabor sequences were defined by this sequential orthogonal shift in orientation.

Mismatch Rotate Gabor sequences were generated by replacing the D frame with a U frame (A-B-C-U-G). Again U frames differed from D frames only by orientation statistics: in this case there was no orthogonal shift in orientation, effectively making (A-B-C-C-G) sequences, where the mismatch was due to a lack of expected change in orientation.

### 4.3 Analysis

#### 4.3.1 Unexpected selectivity index

We quantified the difference in the response to expected and unexpected conditions for a given ROI using an “unexpected selectivity index”, following the approach of Gillon et al. ^17^. This was defined as:

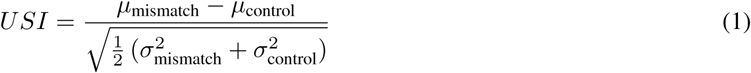

where the means (*µ*_mismatch_ and *µ*_control_) and the variances (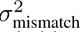 and 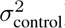) were calculated across the Δ*F/F* across timepoints for the version of the stimulus that violated the habituated sequence, and the control sequence respectively.

For movies, the “mismatch” timepoints corresponded to the reversed segment of the “forward-reverse” condition, while the “control” timepoints corresponded to the identical segment from the “reverse” condition. For Gabor stimuli, the “mismatch” timepoints corresponded to frames U-G of the unexpected A-B-C-U-G sequence, while the “control” timepoints corresponded to frames D-G of the habituated A-B-C-D-G sequence. Gray G frames were included for analysis of the Gabor stimuli as the response elicited by the D frame carried over into the G frame (see Fig. 4B & C).

Summary figures were created by computing the mean and confidence intervals (CI) across ROIs after pooling over subjects. For movie stimuli, a USI for each ROI was first computed for individual movies, then the average taken across the three movies to produce a single average USI value. Confidence intervals were estimated using a hierarchical bootstrap, in which 10^4^ means were recomputed while sampling an equal number of ROIs with replacement, and for each ROI independently resampling trials with replacement.

To assess whether the mean USI was significantly different from chance, and whether mean USIs were significantly different across sessions, we used nonparametric hypothesis tests. To identify sessions with mean USI significantly different from chance, the actual mean USI was compared to 10^4^ values obtained by computing the same metric after shuffling condition labels. To identify significant differences in the mean USI between sessions, the actual difference between means was compared to a distribution constructed by computing the same difference after randomly flipping the session labels for individual ROIs with 50% probability.

To compensate for the increased likelihood of false positives, we applied a Bonferroni correction to all comparisons of a given type. This correction factor was typically 48, to account for comparisons for each combination of two cell lines, two planes, four areas, and three sessions/pairwise comparisons.

#### 4.3.2 Breakdown of USIs

The percentile of each ROI’s USI was computed relative to a shuffled distribution for that ROI. The shuffled distribution for each ROI was obtained by recomputing the USI after randomly permuting the mismatch and control labels 10^4^ times. Histograms show the normalized density of the resulting percentiles, pooling ROIs across subjects.

The mean Δ*F/F* was computed for each ROI during the relevant segment for each trial, and the mean computed across ROIs pooled over subjects to get a single value for each trial. Error bars represent SEM across ROIs. Each unexpected session contained 15 repeats of each movie condition, for each movie type.

The mean USIs were computed for each ROI, for each movie type, then averages were taken across movie types for each ROI. The correlation was computed between the USIs in one session, and the USIs of the same ROIs in another session, to assess if mismatch selectivity was reflected in the same ROIs across different sessions. Correlations computed between the USIs for pairwise combinations of sessions. Significance tested via permutation of day labels.

#### 4.3.3 USIs for orientation selective cells

To identify orientation selective cells, presentations of each type of Gabor stimulus were split into a set for the unexpected sequences, a training set of expected sequences to identify orientation selective cells, and a held-out test set of expected sequences of equal size for validation. The held-out test set was constructed by first collecting the closest expected sequence to each unexpected sequence whose D orientation matched the U orientation to build a subset that could be used in comparisons with the mismatch condition. Additional remaining expected sequences were randomly selected so as to match the size of the train split, for use in validation of tuning properties. A gOSI for each ROI was computed based on its mean activity across presentations of the D frame for each orientation. A null distribution of gOSIs for each ROI was constructed by shuffling orientation labels, and selective ROIs identified as those with a gOSI magnitude in the top 5 % of shuffled values.

For each orientation, control and mismatch sequences were identified as those whose D frame or U frame matched that orientation. A population-level mean Δ*F/F* was then computed at each timepoint for these conditions using only selective ROIs with that preferred orientation. A USI for each selective ROI was similarly computed over presentations of the D/U/G frames for its preferred orientation, and for comparison, a USI was also computed across all presentations of non-preferred orientations. Finally, USIs for non-selective cells were computed across all stimulus presentations as another point of reference.

#### 4.3.4 Comparison of somato-dendritic coupling

For each pair of somatic and dendritic ROIs matched to the same cell, the Pearson correlation between somatic and dendritic time series was computed over concatenated time points belonging to each condition, where the relevant time points for unexpected and expected sequences were defined as in Methods 4.3.1. A significant difference between the two distributions for each session was tested for using a nonparametric hypothesis test. To correct for multiple comparisons, p-values were multiplied by a Bonferroni correction of 3.

The multi-plane mesoscope introduced a potential confound, as signals from each plane are interleaved, and can cause ‘ghosting’ between planes if the sensor has not had time to reset between scans. To mitigate the effect of ghosting on our coupling analyses, we eliminated from all coupling comparisons pairs of ROIs which at any point were imaged using the same sensor. Fortunately, due to the anatomy of pyramidal neuron apical arbors and the orientation of the cells across planes, dendritic ROIs from a given neuron tended to be spread out from 50-300 µm away from the connected soma by the time they reached L1.

Calcium transients were identified using a multi-stage pipeline, informed by the ground-truth comparisons of somatic calcium signals for the Cux2 and Rbp4 lines ^72,91^. We used a model, trained on ground truth simultaneous recordings, to produce deconvolved estimated spike rate series for each ROI (CASCADE, GCaMP6f excitatory neuron model, noise-matched to our 10 Hz imaging conditions ^92^). The output spike probability time series was smoothed using a 1D Gaussian filter (*σ* = 1 frame) to reduce sample jitter while maintaining temporal alignment. Peaks in the smoothed probability series exceeding a threshold of 0.30 were identified as candidate event times.

At each candidate time, the Δ*F/F* signal was assessed using two criteria, either of which was sufficient to confirm a transient: (a) the 1D Gaussian-smoothed (*σ* = 1 frame) Δ*F/F* crossed 2*×* the median absolute deviation (MAD) of the ROI’s baseline Δ*F/F* for at least 200 ms (2 frames at 10 Hz), within 500 ms of the CASCADE probability peak or (b) the raw Δ*F/F* exceeded 3*×* MAD above the baseline median, within 500 ms of the CASCADE probability peak. The 2 MAD / 200 ms primary criterion, although somewhat liberal, is more likely to correspond to multiple spikes within our 100 ms sampling window ^72^. The 3 MAD raw secondary criterion was added to capture smaller, briefer depolarizations, particularly in dendritic ROIs, which are still well above the noise floor. Both conditions are intentionally inclusive to ensure that smaller but unmistakable coactivation events in a connected pair were still captured (see technical transients below).

Onset timing was refined from the spike rate peaks to sub-frame precision using PCHIP (Piecewise Cubic Hermite Interpolating Polynomial) interpolation of the Δ*F/F* time series, upsampled to 10 ms resolution (10*×* the 100 ms frame interval). The onset was identified as the local minimum in the upsampled Δ*F/F* within the 300 ms window preceding the peak, the local minimum at which the signal begins its sustained rise. These onsets were vetted for temporal precision against both the spike rate traces and estimated spike timing. The PCHIP-based onset time was used for all stimulus alignment and inter-compartment timing comparisons.

A coincidence window of *±*300 ms was used to define simultaneity between soma and dendrite events. This window is calibrated to GCaMP6f kinetics at 10 Hz, 50-100 ms rise time, and *∼*150–300 ms decay time. A 300 ms window is narrower than that used by some prior studies ^91^ but avoids spurious coincidences from temporally adjacent but mechanistically unrelated events. Given the compartmental sets of identified transients, for a given identified matched pair, these were further classified as coupled (within *±*300 ms of a transient in the other compartment) or independent (labelled either dendritic or somatic). The independent transients were subject to a secondary comparison, to check for smaller depolarization in the matched compartment.

For the independent somatic or dendritic events, the calcium series of matched ROI was assessed using the 2/3 MAD screening described above, to capture coactivation events not identified in our initial screening, which likely still demonstrate depolarization. Those which failed this secondary screening made up our final set of isolated compartmental transients. Whenever identified, these technical transients were labeled as ‘weak’ and only treated on equal footing with other coupled transients after this reassignment was vetted by separate analyses (i.e., confirming that excluding/reassigning these did not impact any results or their statistical significance).

Transient rates were calculated per matched cell (Hz - transients per second). The magnitude of each transient was defined as the difference between the peak Δ*F/F* and the onset Δ*F/F*. These magnitudes were normalized by the maximum transient magnitude for each compartment, to facilitate comparison across cells, areas, subjects and lines. We exploited the same stimulus specific windows as defined above for the USI analyses Methods 4.3.1. We adopted a similar nonparametric hypothesis test, shuffling conditional (mis)match labels 10^4^ times, to identify significant differences in transient rate or size.

### 4.4 Matching ROIs between somatic and dendritic recording planes

#### 4.4.1 Reconstructing cells using volumetric imaging

For a given visual area, individual z-stack planes were motion corrected using a modified version of NoRMCorre and denoised by a modified pipeline using DeepInterpolation ^86,93,94^. Because we combined multiple z-stacks for a given area, planes were first matched in the z-depth. One stack was used as a template scaffold, and the other stacks were aligned to structures in it using a constrained sequence of global rigid, patch-wise rigid and patch-wise optical flow registration (OpenCV, scikit-learn) ^95,96^. The combined stack was then aligned using a constrained global rigid registration to ensure that cross-plane structures were aligned across all z-depths. We also developed a local contrast adjustment algorithm to reveal structures with low relative brightness.

Cells within these z-stacks were traced using Neurolucida 360 (Fig. S10) ^97^. First, somata with a visible match in the somatic ROI masks were identified. The apical dendrites of traced somata were then traced upwards to the depth of the dendritic imaging plane. Tracings were obtained from five L5 subjects for VISp and VISl, and three subjects for VISpm and VISam. Our Cux2 L2/3 subjects proved to have too much latent neuropil GCaMP expression, which obscured fine dendritic structural detail, making tracing impossible.

#### 4.4.2 Matching ROIs and evaluating results

To align the experimental movies to the z-stacks, we manually defined a target depth range, and set of likely best depths. We then optimized local image comparisons with custom structural metrics to identify the best depths with matching fine structures. To better align specific ROIs to structures in the z-stacks, we also developed a constrained sequence of global rigid, patch-wise rigid and patch-wise affine registration. Dendritic masks were also re-processed to extract the most accurate structural detail provided by the EXTRACT probability maps ^41,98,99,100^. These aligned masks were then automatically compared to reconstructed neurons from our tracing, producing sets of potential matches between ROIs and traced cells.

Potential somatic and dendritic matches were then manually curated by two independent researchers and their assessments combined to yield a rating of matching likelihood. Potential matches were scored 0 for impossible, 1 for possible, 2 for likely, 3 for highly probable. The scores from independent assessments were summed to produce a rating from 0 to 6. Somatic and dendritic ROIs receiving a score of 3 or below were filtered, so that remaining ROIs received an overall rating of 4 or higher. Pairs of remaining ROIs matched to the same tracing were considered matched to the same cell (Table S1).

### 4.5 Pseudo-matching ROIs between somatic and dendritic recording planes

An extended pseudo-matching procedure was developed to assign dendrite ROIs to their parent soma without direct anatomical reconstruction (k-to-1, dendrite to soma matching). Population-wide synchronous activity inflated raw somato-dendritic correlation uniformly across true and false candidate pairs, especially in L2/3. Exploiting the large calcium transients, which more selectively reflect burst-like whole-cell firing, proved to be more effective for matching. Onset frames were binarized and dilated by a ±3-frame coincidence window, and each candidate pair was scored by a bidirectional overlap statistic (Eq. 2) ^98,99,100^. The minimum of the fraction of each compartment’s activity explained by the other was weighted by each event’s magnitude, rather than counted uniformly. This weighting was motivated by physiological studies demonstrating that the largest depolarization events in dendrites or somata are most likely to correspond to global neuronal firing ^72,73^. It was also confirmed against the high confidence connected pairs: an event’s magnitude correctly ranked it as shared (versus independent) 77% of the time on the soma side and 80% on the dendrite side, assessed by the area under the ROC curve. Screening against the bidirectional fractions penalized somata/dendrites with substantial uncorrelated activity, similar to intersection-over-union; weighting both the numerator and denominator by magnitude penalizes a large unmatched event without imposing a hard veto.

For dendrite *d* and soma *s*, let *t* index *d*’s onset frames for transient with normalized magnitude *m_d_*(*t*), and *t^′^* index *s*’s onset frames with magnitude *m_s_*(*t^′^*). Each onset train is dilated by *±*3 frames to define a coincidence window.

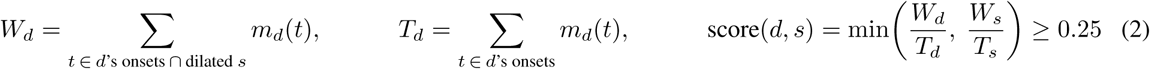

For each dendrite, the best-scoring soma and runner-up were compared. Population synchrony can inflate a false runner-up, and further non-linear dynamics and imaging differences made a fixed ratio threshold unreliable. A pair was accepted only if that margin survived a block bootstrap (1000 resamples of non-overlapping 300-frame blocks), i.e., the lower 5th-percentile bound of this resampled ratio exceeded 1.0, meaning the best-scoring soma remained the better match, even compared to the runner-up, in at least 95% of the resampled reconstructions. ROIs were also screened by noise, skewness, event count, and peak amplitude, to ensure valid activity dominated the matching, not outliers or noise. However, only 60.0 % (503/839) of high-confidence traced ROI pairs (c *≥* 4) passed this conservative functional recording quality control (QC) screen.

The above method was validated against an independent reference set of 237 L5 soma-dendrite pairs confirmed by reconstruction/tracing and a functional criterion, uncontaminated by the matching parameters (QC pass, bump-denoised correlation of CASCADE traces > 0.3, and runner-up ratio *≤* 2.0). 88.2 % of these pairs were recovered (209/237), using a bidirectional coactivation score floor of 0.25. This validation was limited to L5, but the same method/parameters were applied to identifying potential L2/3 matched pairs, without independent confirmation. Out of dozens of tested methods, the above yielded the cleanest conservative set of matches across areas, subjects and cell lines (see Table S6, Table S8). Pairs were post screened for ghosting (Methods 4.3.4), but no ROIs with spatial overlap were matched.

## 5 Acknowledgements

This work was supported by NSERC (Discovery Grant: RGPIN-2020-05105 (BAR) and RGPIN-2019-06379 (JZ); Discovery Accelerator Supplement: RGPAS-2020-00031 to BAR); the Joan and Jerome Snyder Chair in Vision Science (JZ); and CIFAR (Canada AI Chair to BAR; Learning in Machines and Brains Fellowships to BAR and JZ); Mitacs Research Training Award (TMH); and Ontario Graduate Scholarship (TMH). This research was enabled in part by support provided by Calcul Québec (https://www.calculquebec.ca/en/) and the Digital Research Alliance of Canada (https://alliancecan.ca/en). The authors acknowledge the material support of NVIDIA in the form of computational resources.

The experimental dataset in this project was obtained as part of the OpenScope program, which is operated by the Allen Institute, Neural Dynamics program and funded by US NIH grant U24NS113646 (JAL, CK). We thank the OpenScope steering committee and Karel Svoboda for their support. This research was also supported by the Allen Institute, founded by Jody Allen – chair and co-founder of Allen Family Philanthropies, and the late Paul G. Allen – investor, philanthropist, and co-founder of Microsoft. We gratefully acknowledge their vision and generosity, which made this work possible.

The authors would also like to offer special thanks to Flavia Arias, for their assistance with neuron tracing in volumetric z-stacks, and to Colin Bredenberg for feedback on the orientation-selective analyses.

## 6 Supplement

**Figure S1.**
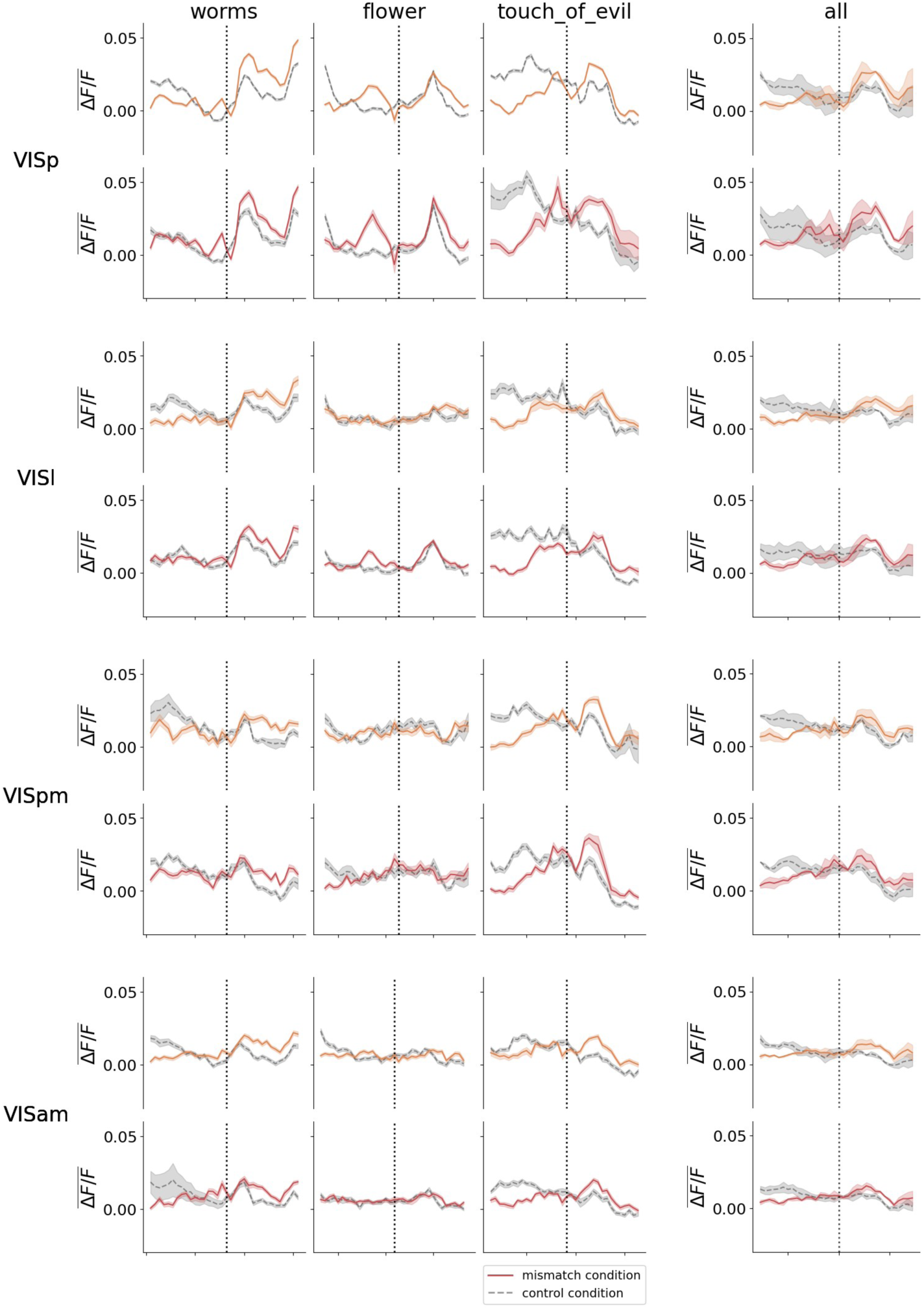
Time traces at onset of mismatch segment for U1 session.

**Figure S2.**
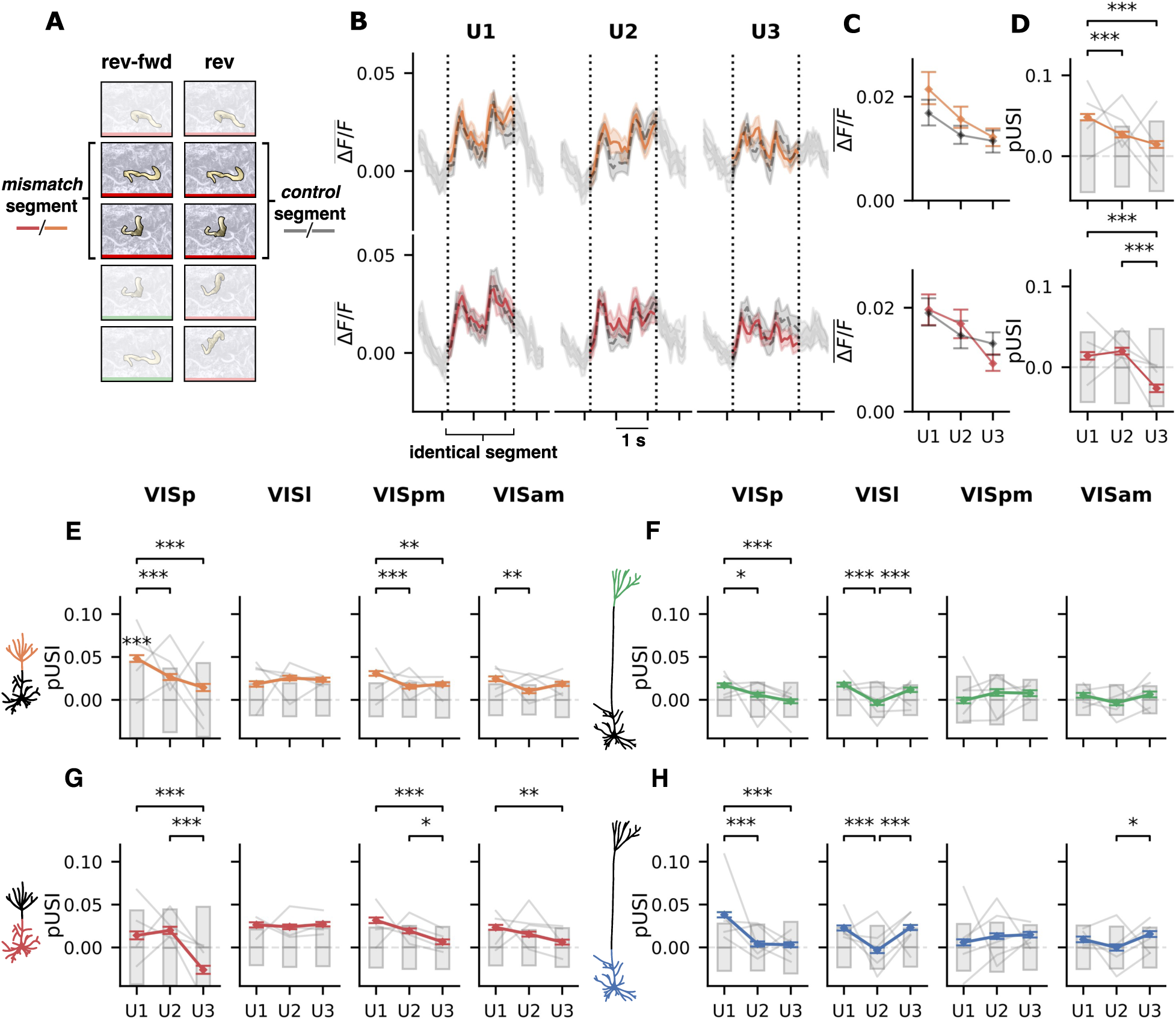
“Same-same” comparison of responses to an identical segment in the “rev” and “rev-fwd” conditions. **(A)** Depiction of the control (reverse) and substitute for the mismatch (reverse-forward) movie sequences, with labels marking the identical segment of clip used in subsequent comparisons. Green frames represent forward playback, while red frames represent reverse playback. Because reverse and reverse-forward conditions begin identically, the response to the segment in these conditions should be identical. **(B)** Mean Δ*F/F* across ROIs and trials during presentation of the same movie segment in reverse and reverse-forward conditions. Colored series represent the responses in the reverse-forward condition, and grey the responses to the same segment when presented as part of the reverse condition. Orange series correspond to apical dendritic responses and red series to somatic responses. Dashed vertical lines indicate the window in which the stimuli match across conditions, with grey tails for each series outside of the window added for context. Colored bands represent a 95 % CI estimated using a hierarchical bootstrap. **(C)** Mean of the windowed portion of the Δ*F/F* in (B) over ROIs, trials, as well as time, yielding one value per session, for sessions U1, U2, and U3. Error bars represent a 95 % CI estimated using a hierarchical bootstrap. **(D)** “Pseudo” unexpected selectivity index (pUSI) computed for the same Δ*F/F* as in (B) and (C). Colored series represent the mean pUSI across ROIs for each session, grey bars represent the 95% CI of a distribution constructed from permutations. Error bars represent a 95 % CI estimated using a hierarchical bootstrap. **(E)-(H)** Selectivity of responses to the mismatch condition (pUSIs), across areas and cell lines, averaged over L2/3 distal apical dendritic ROIs **(E)** and L2/3 somata **(G)**, and L5 apical dendrites **(F)** and L5 somata **(H)**. Grey bars represent 95% CI for a permutation based null distribution. Error bars represent a 95 % CI estimated using a hierarchical bootstrap. Fine grey lines show average individual subject pUSIs. * *p <* 0.05, ** *p <* 0.01, *** *p <* 0.005.

**Figure S3.**
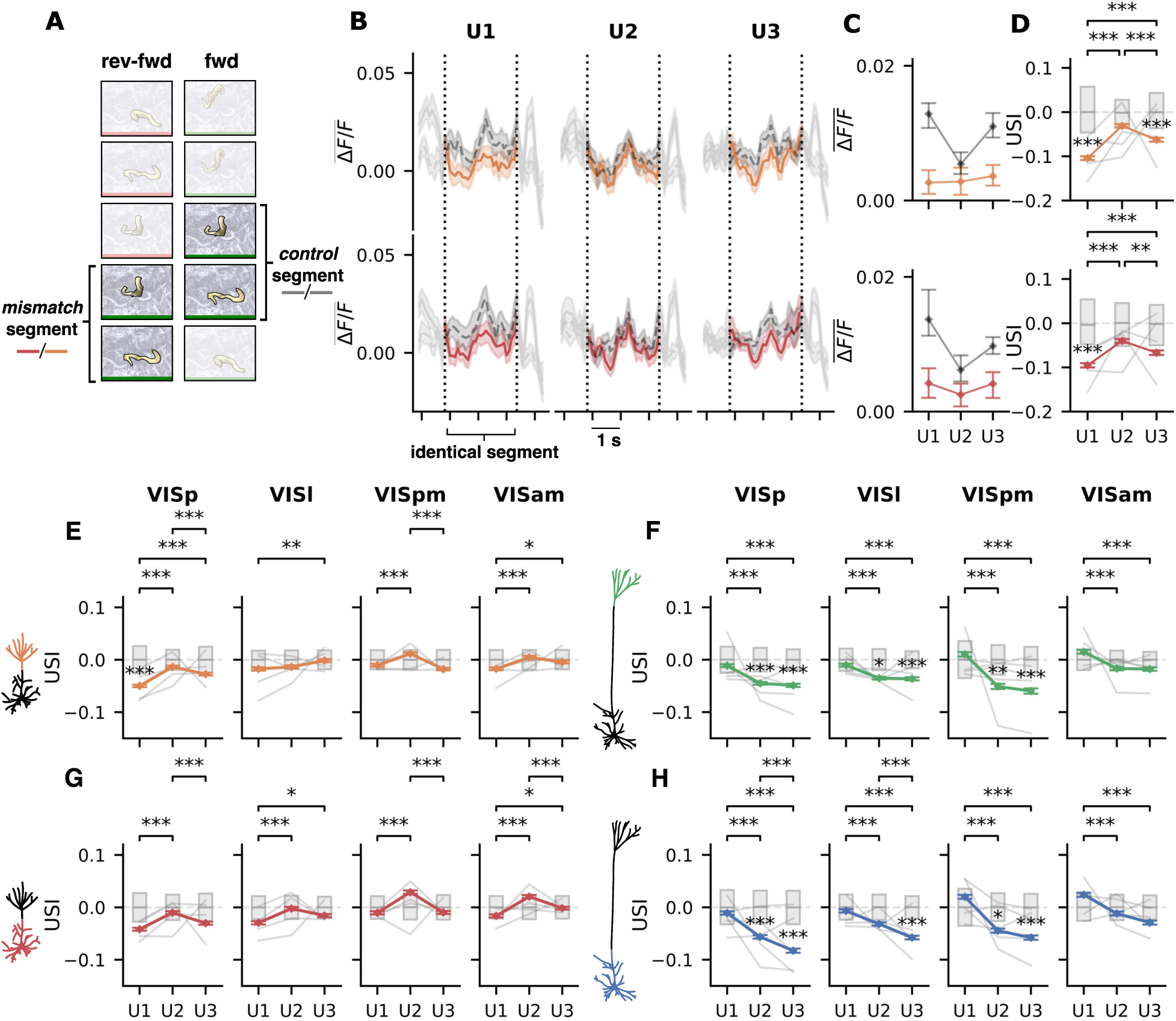
Comparison of responses to an identical segment in the “fwd” and “rev-fwd” conditions. Same as Fig. S2, but with the “fwd” sequences in the role of the control condition, and “rev-fwd” sequences in the role of the mismatch condition.

**Figure S4.**
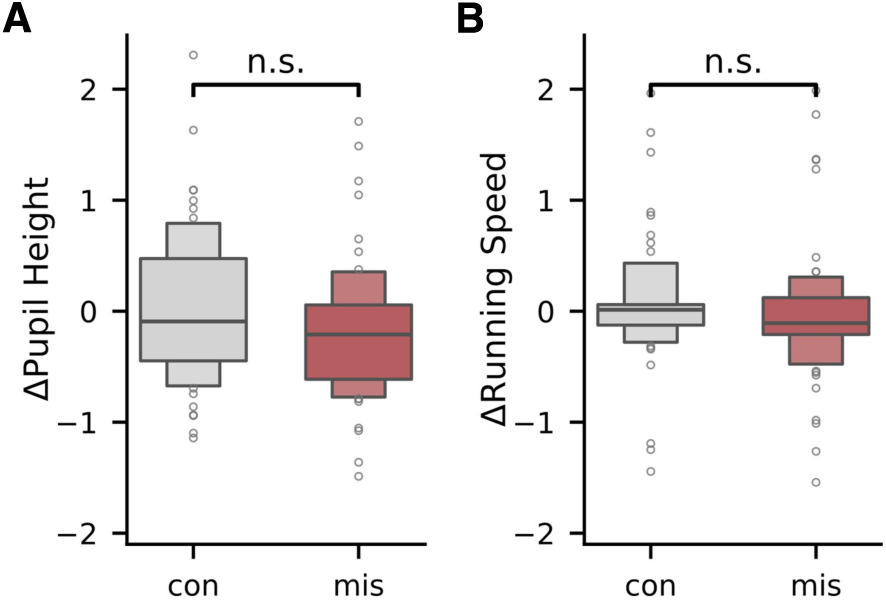
Absence of behavioral differences across mismatch conditions. **(A)** Distribution of z-scored changes in pupil height across trials in each mismatch condition. **(B)** Distribution of z-scored changes in running speed across trials in each mismatch condition.

**Figure S5.**
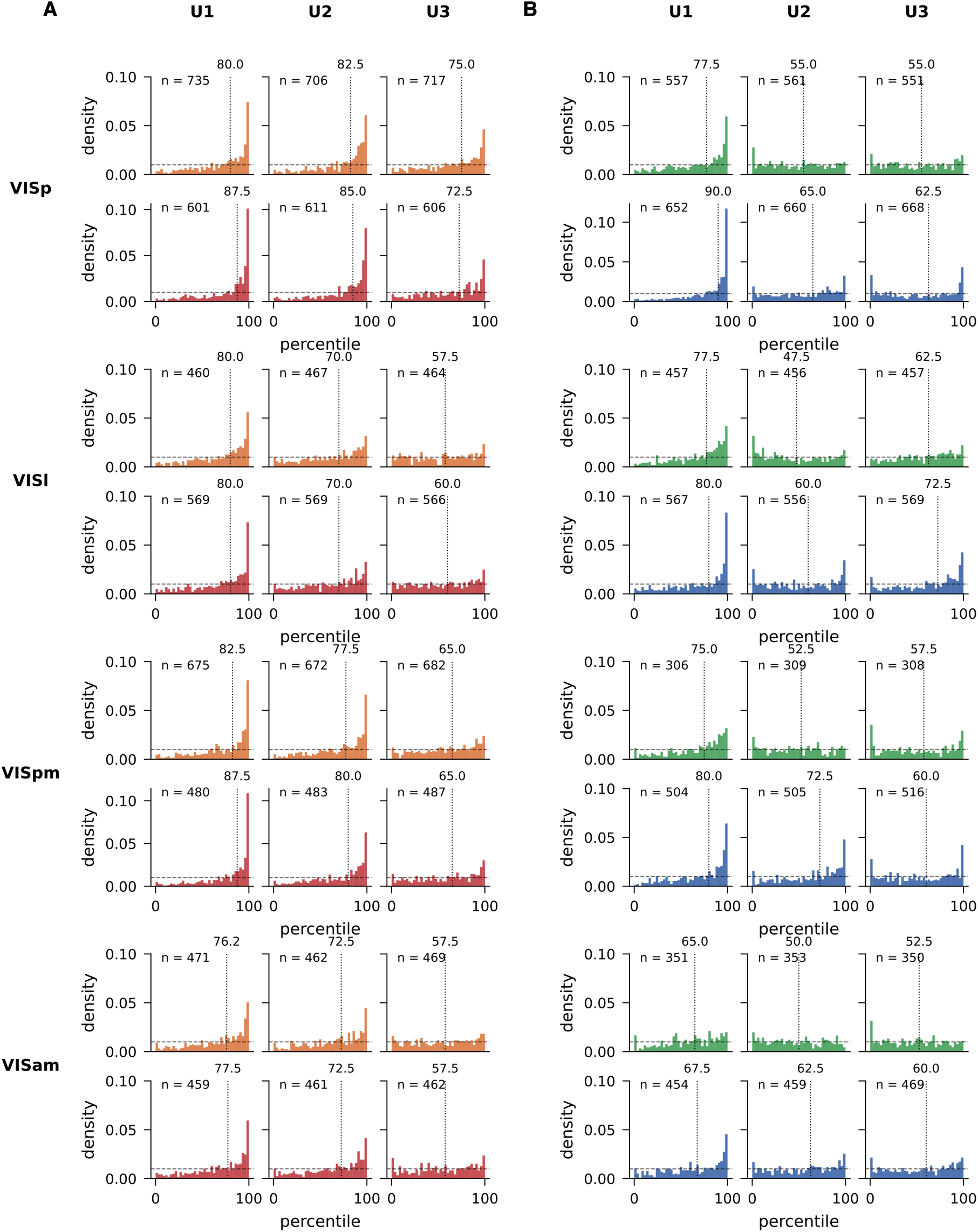
USI percentile distributions across all areas and sessions for natural movie stimuli. **(A)** Percentile distributions for L2/3 ROIs. **(B)** Percentile distributions for L5 ROIs.

**Figure S6.**
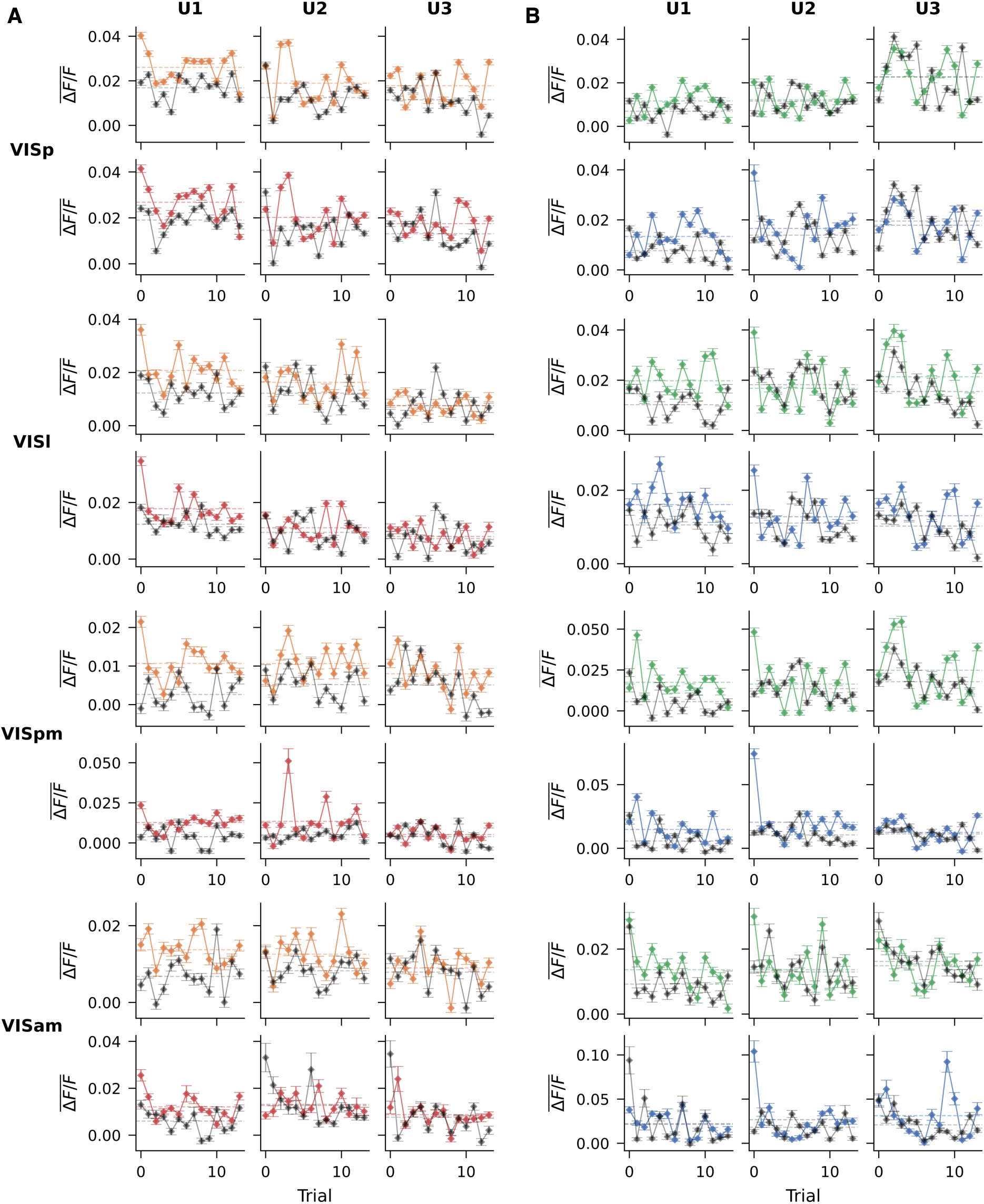
Mean activity by trial index across all areas and sessions for natural movie stimuli. **(A)** Mean activity plots for L2/3 ROIs. **(B)** Mean activity plots for L5 ROIs.

**Figure S7.**
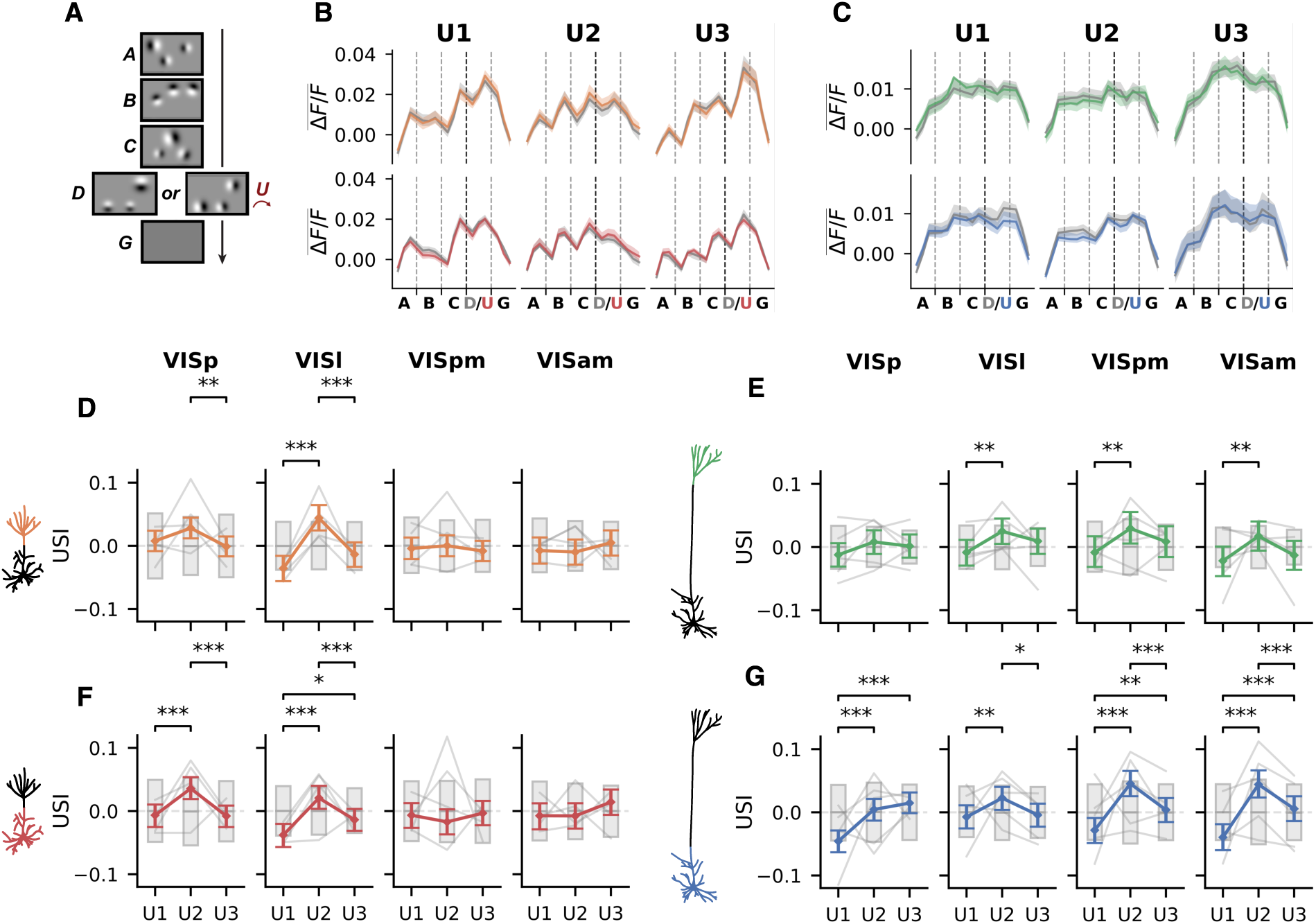
Analysis of mean responses by condition for sequences of rotating Gabor patch stimuli. **(A)** Structure of rotating Gabor patch stimuli. Subjects were habituated to sequences of frames A-B-C-D-G, in which patches in each frame were drawn from a distribution with a mean orientation that rotated by 90 degrees relative to the preceding frame. Unexpected sequences A-B-C-U-G were then introduced, in which the U frame had patches in the same location as D, but with the *same* mean orientation as frame C. **(B)**, **(C)** Mean Δ*F/F* across trials and ROIs for L2/3 and L5 respectively. Colored lines represent the mean response to the mismatch sequences (A-B-C-U-G), while grey represents the mean response to the control sequences (A-B-C-D-G). Vertical grey dashed lines demarcate the timepoints belonging to each frame in a sequence. Vertical black dashed line indicates the point of divergence in the two kinds of sequence. **(D)**-**(G)** USIs for sequences of Gabor patches with rotating orientations across areas, sessions, and cell lines. Grey bars represent 95% CI for a permutation based null distribution. Error bars represent a 95 % CI estimated using a hierarchical bootstrap. Fine grey lines show average individual subject USIs. * *p <* 0.05, ** *p <* 0.01, *** *p <* 0.005.

**Figure S8.**
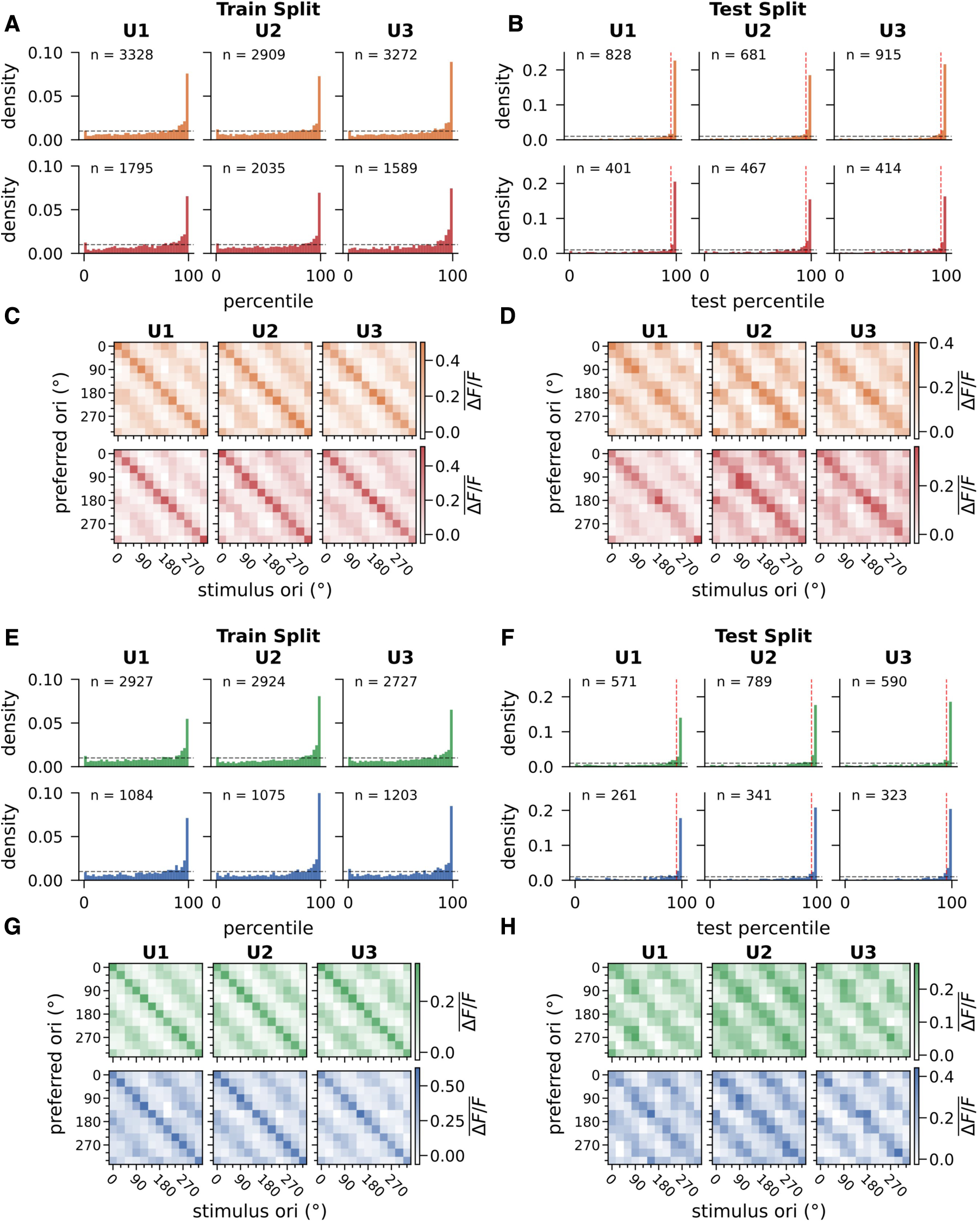
Extended analysis of orientation selectivity supporting Fig. 5. **(A)** Distribution of gOSI percentiles on the training split for L2/3 ROIs. **(B)** Distribution of gOSI values on the test split for L2/3 ROIs identified as orientation-selective based on their percentile in the training split. **(C)** Mean activity across stimulus orientations on the training split by preferred orientation for cells identified as orientation-selective. **(D)** Mean activity across stimulus orientations on the test split by preferred orientation for cells identified as orientation-selective. **(E)**-**(H)** Same as (A)-(D) but for L5 ROIs.

**Figure S9.**
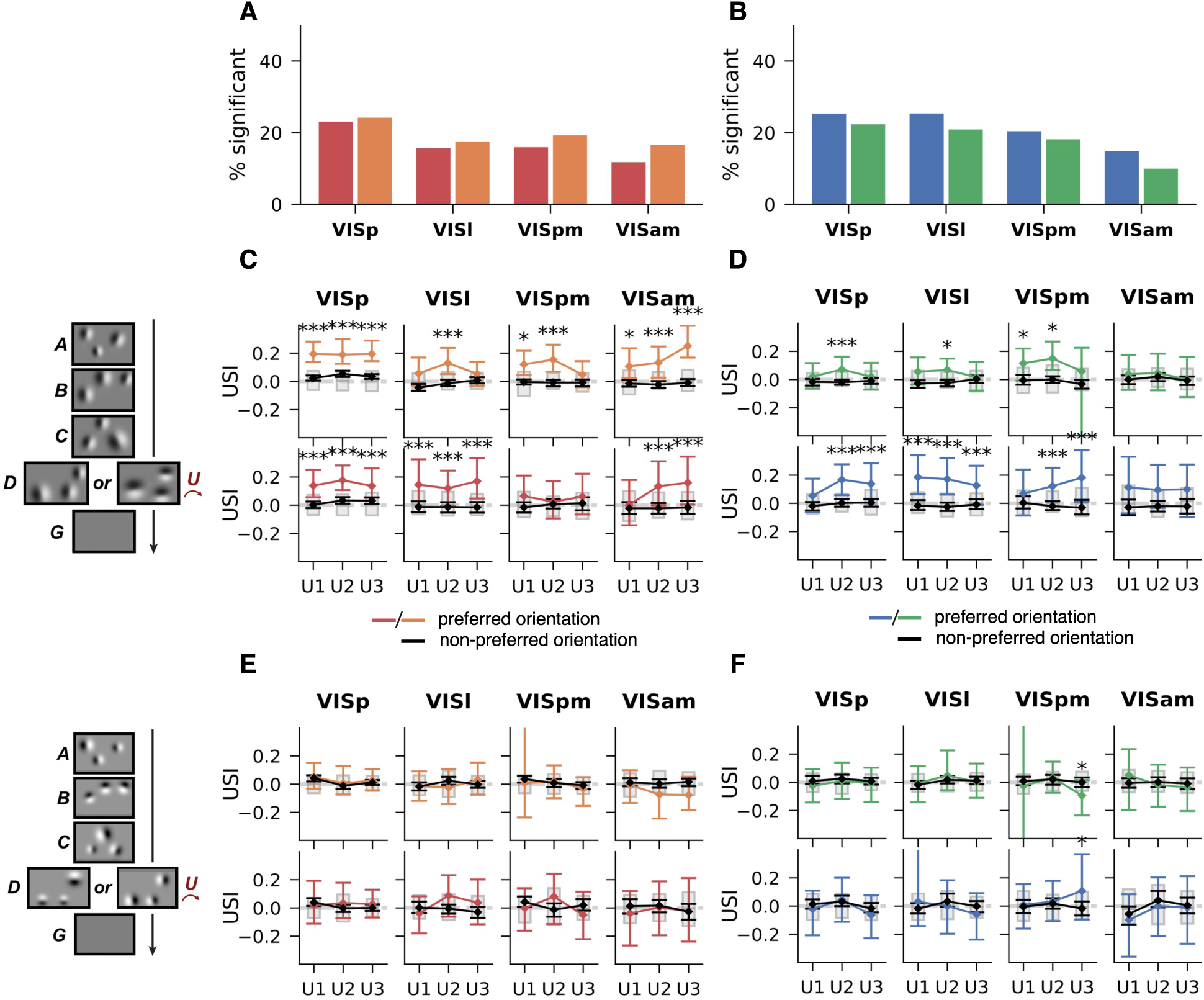
Full USI results for orientation selective USIs as in Fig. 5. **(A)**, **(B)** Percentage of L2/3 and L5 ROIs identified as significantly orientation selective. **(C)**, **(D)** Mean USIs for fixed Gabor stimuli across areas, sessions, and cell lines for significantly orientation-selective ROIs. **(E)**, **(F)** Same as (C), (D) but for rotating Gabor stimuli. * *p <* 0.05, ** *p <* 0.01, *** *p <* 0.005.

**Figure S10.**
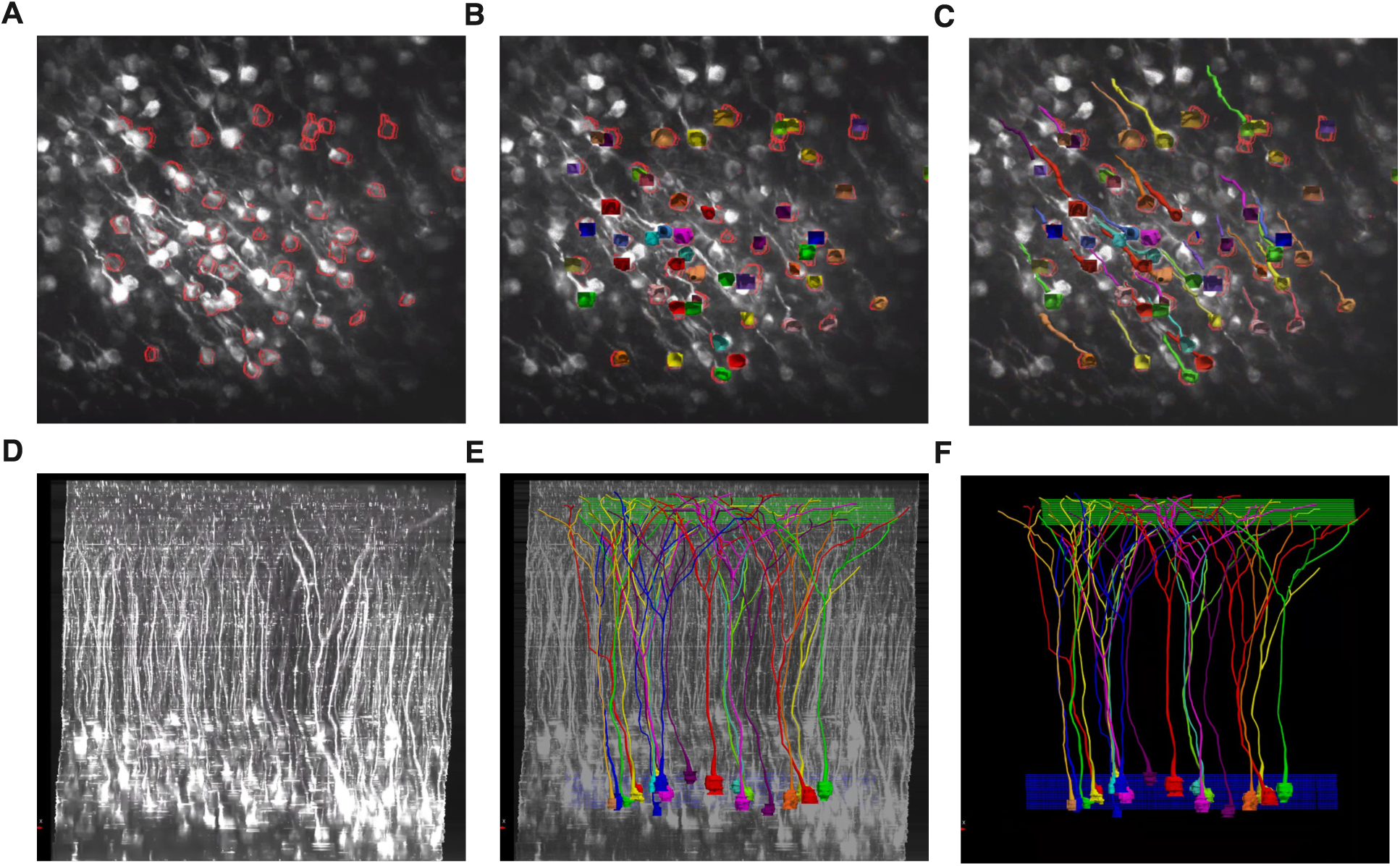
Z-Stack tracing process. **(A)** Somatic ROIs segmented from functional imaging recordings outlined in red, superimposed on z-stack. **(B)** Traced somata in z-stack found to match a superimposed mask. **(C)** Apical arbors were then traced upwards from each traced soma. **(D)** Z-stack without tracing. **(E)** The same Z-stack as (D) with tracing superimposed, and functional imaging planes indicated in green (dendritic) and blue (somatic). **(F)** The same tracing and functional imaging planes as (E) without the z-stack.

**Figure S11.**
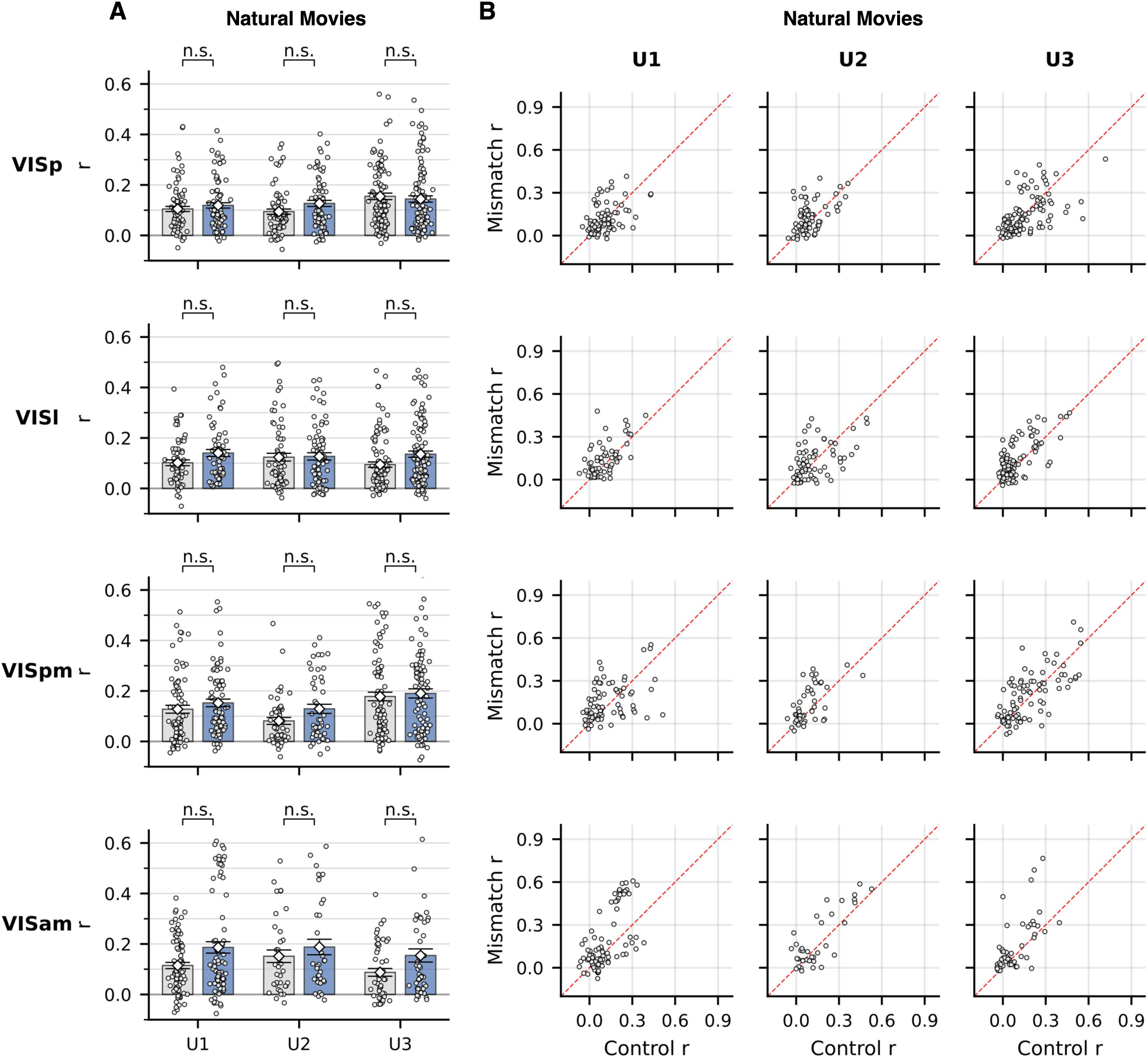
Pearson correlations for natural movie stimuli. **(A)** Bar plots showing mean Pearson correlation in control and mismatch conditions. Colored bars represent mean correlation in the mismatch condition and gray that in the control condition, with errorbars for the SEM. Scatterpoints represent the correlation of individual soma-dendrite ROI pairs. **(B)** Scatterplots showing relationship between Pearson correlations in each condition. Scatterpoints represent individual soma-dendrite ROI pairs. Dashed red line represents y = x. * *p <* 0.05, ** *p <* 0.01, *** *p <* 0.005.

**Figure S12.**
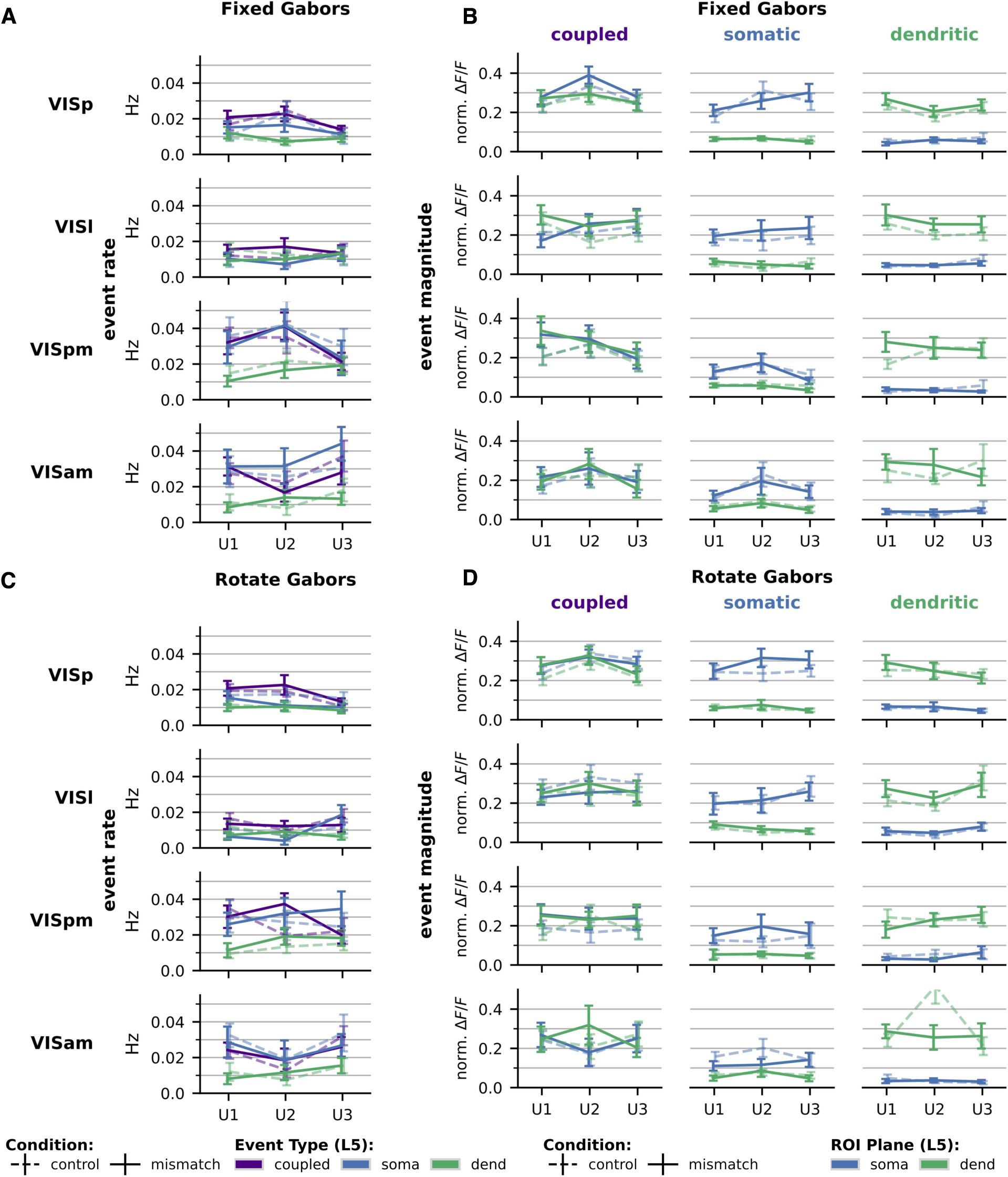
Extended analysis of somato-dendritic coupling using Δ*F/F* transients for L5 connected soma-dendrite pairs identified by tracing in volumetric imaging. **(A)** Rates of transients for each class, averaged across connected pairs, for fixed Gabor sequences. **(B)** Normalized Δ*F/F* magnitudes of transients for each class, averaged across connected pairs, for fixed Gabor sequences. **(C)** Same as (A), for rotate Gabor sequences. **(D)** Same as (B), for rotate Gabor sequences. * *p <* 0.05, ** *p <* 0.01, *** *p <* 0.005. See Table S4 for number of pairs used for each comparison.

**Figure S13.**
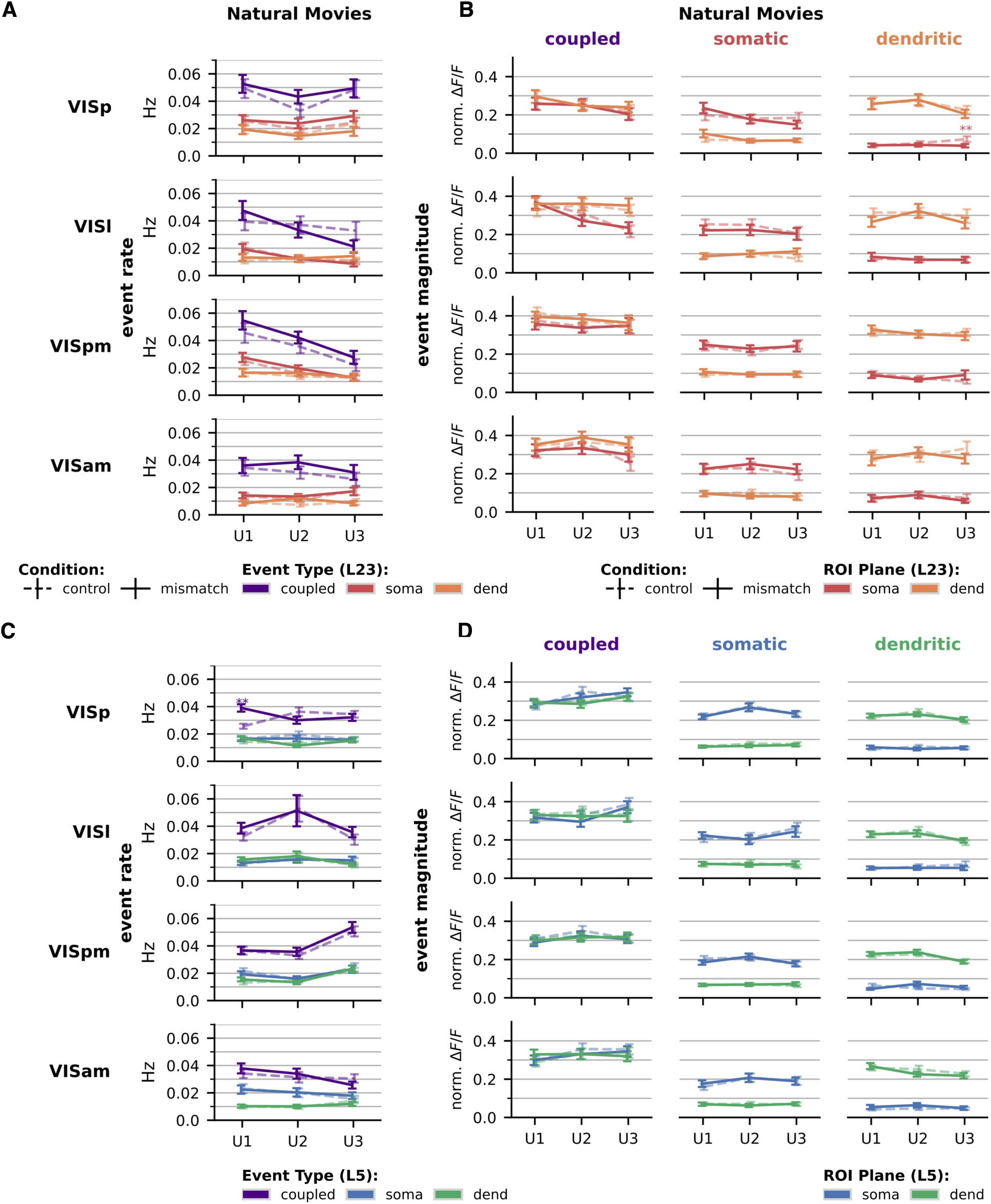
Analysis of transients for soma-dendrite pairs matched based on coupling, for natural movie stimuli. Same event rate and event magnitude comparisons as fig. 7. For L2/3 **(A)**, **(B)** and L5 **(C)**, **(D)**. * *p <* 0.05, ** *p <* 0.01, *** *p <* 0.005. Although matching based on coupling may yield a biased set of pairs, the modest matching threshold (Methods 4.5) allows for the independent activity seen above.

**Figure S14.**
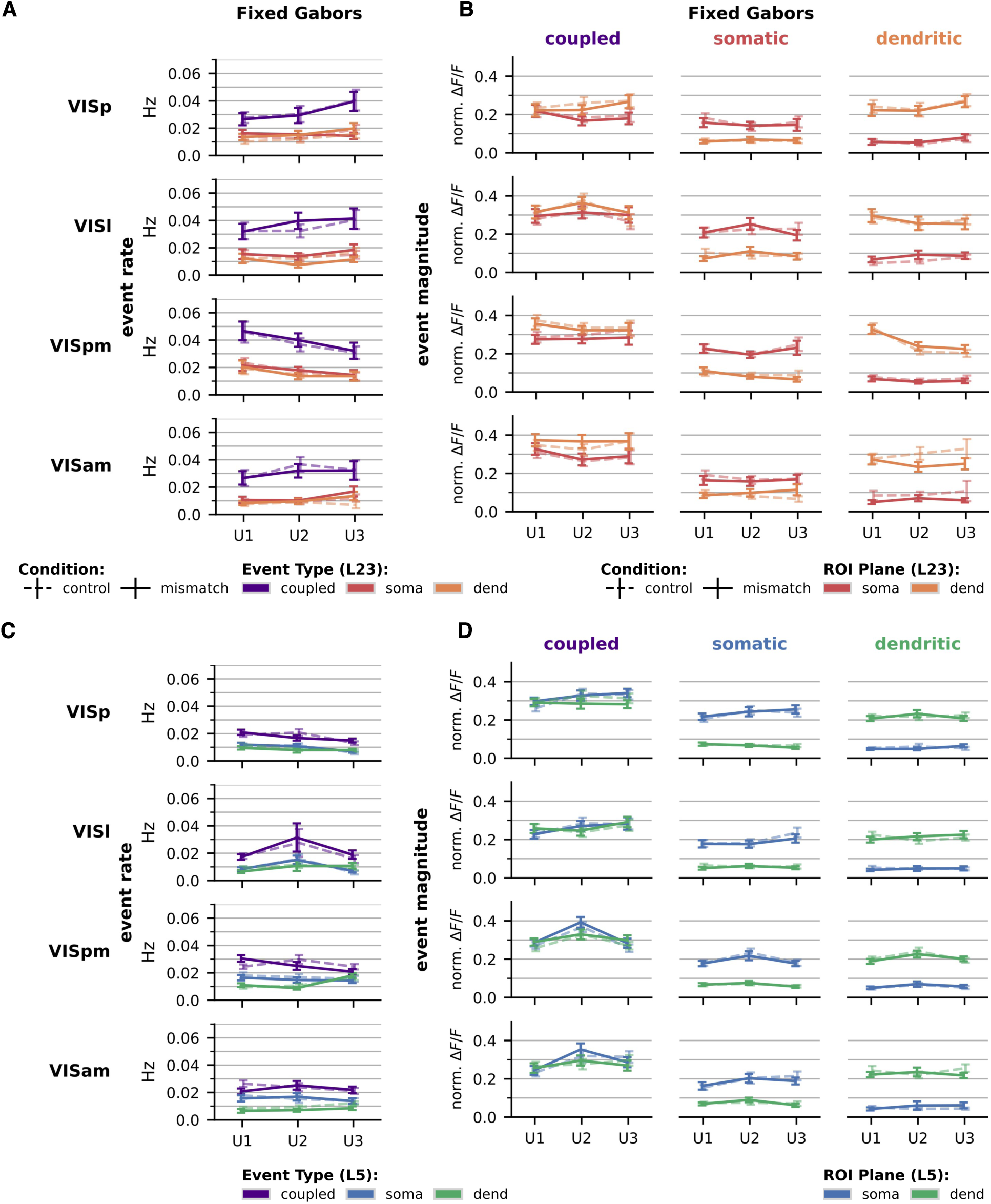
Same as Fig. S13, transients for soma-dendrite pairs matched based on coupling for fixed Gabor stimuli.

**Figure S15.**
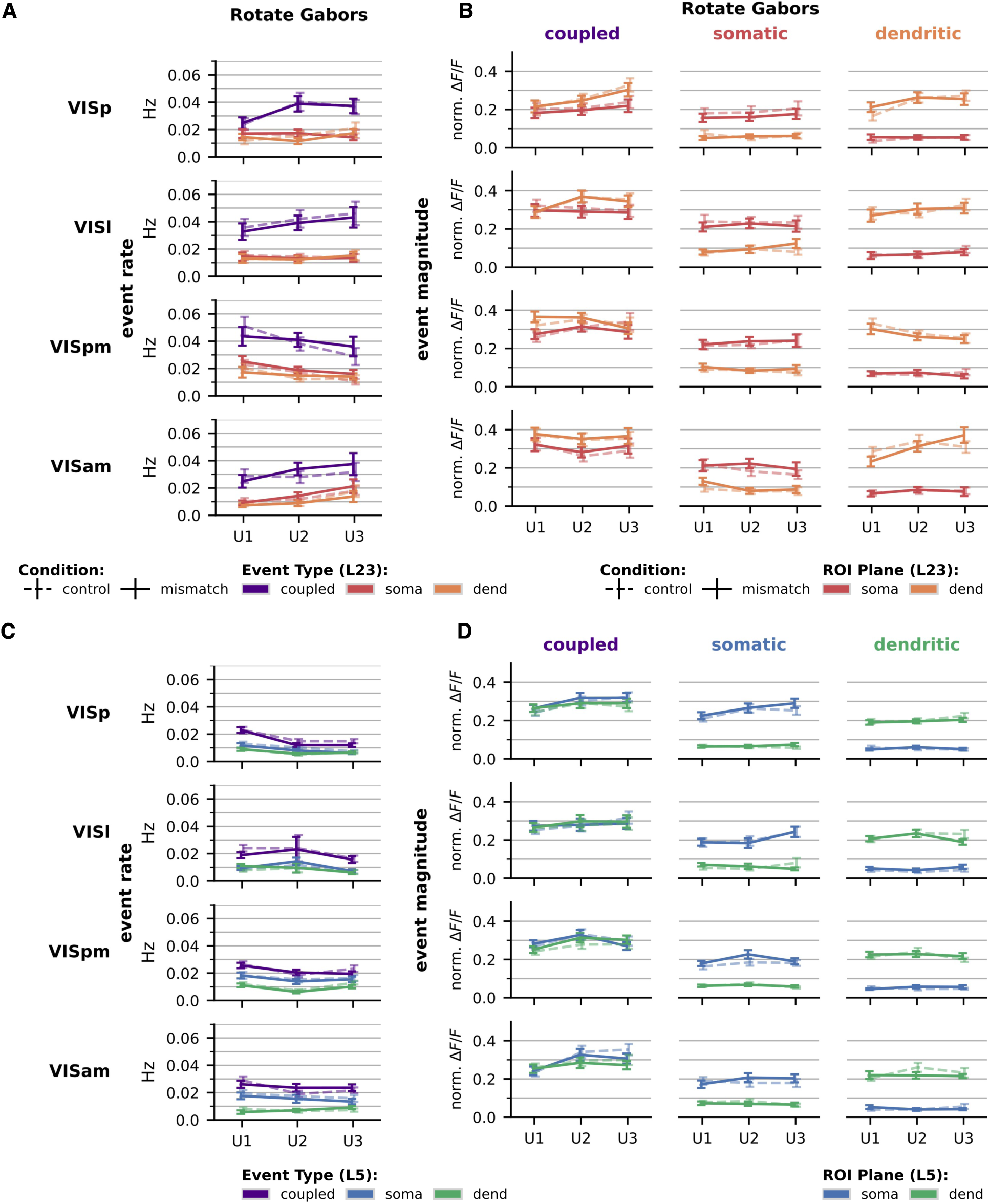
Same as Fig. S13, transients for soma-dendrite pairs matched based on coupling for rotating Gabor stimuli.

**Table S1.** Anatomically matched L5 soma-dendrite pairs identified via tracing, organized by visual area, compartment, and confidence threshold. Counts summed across all across U-sessions.

| Confidence | Area | Soma (ROIs) | Dend (ROIs) | Pairs |
|---|---|---|---|---|
| *c* ≥ 1 | VISp | 278 | 606 | 663 |
|  | VISl | 175 | 388 | 411 |
|  | VISpm | 132 | 254 | 272 |
|  | VISam | 115 | 257 | 281 |
| *c* ≥ 2 | VISp | 258 | 496 | 531 |
|  | VISl | 163 | 343 | 362 |
|  | VISpm | 132 | 254 | 272 |
|  | VISam | 115 | 257 | 281 |
| *c* ≥ 4 | VISp | 190 | 282 | 297 |
|  | VISl | 126 | 213 | 226 |
|  | VISpm | 86 | 140 | 148 |
|  | VISam | 80 | 153 | 168 |
| *c* ≥ 5 | VISp | 113 | 144 | 151 |
|  | VISl | 73 | 102 | 108 |
|  | VISpm | 43 | 61 | 63 |
|  | VISam | 34 | 64 | 74 |

**Table S2:** L5 tracing-matched (confidence>=4), number of pairs per area with transients during natural movies (flower / worms / touch of evil) during each session, separated by condition (mismatch (mm) / control (ct) / intersection (int)) and transient class (Fig. 7). Intersection (int) includes all pairs with transients in both mismatch (mm) and control (ct) conditions for a given stimulus.

|  |  | natural movie | flower |  |  | worms |  |  | touch of evil |  |  |
|---|---|---|---|---|---|---|---|---|---|---|---|
|  |  |  | ct | mm | int | ct | mm | int | ct | mm | int |
| area | day | class |  |  |  |  |  |  |  |  |  |
| **VISp** | U1 | coupled | 30 | 30 | 19 | 25 | 26 | 18 | 15 | 20 | 7 |
|  |  | dendritic | 16 | 20 | 8 | 13 | 17 | 5 | 10 | 17 | 5 |
|  |  | somatic | 19 | 19 | 11 | 15 | 18 | 9 | 9 | 13 | 5 |
|  | U2 | coupled | 24 | 22 | 14 | 25 | 18 | 12 | 22 | 17 | 7 |
|  |  | dendritic | 19 | 15 | 7 | 14 | 12 | 2 | 12 | 8 | 2 |
|  |  | somatic | 13 | 16 | 10 | 12 | 7 | 4 | 10 | 11 | 7 |
|  | U3 | coupled | 41 | 36 | 29 | 38 | 34 | 23 | 26 | 30 | 16 |
|  |  | dendritic | 21 | 22 | 10 | 24 | 22 | 15 | 20 | 20 | 10 |
|  |  | somatic | 24 | 24 | 16 | 18 | 24 | 13 | 22 | 22 | 15 |
| **VISl** | U1 | coupled | 17 | 19 | 12 | 18 | 13 | 7 | 12 | 16 | 9 |
|  |  | dendritic | 9 | 13 | 8 | 10 | 9 | 3 | 8 | 11 | 3 |
|  |  | somatic | 10 | 9 | 5 | 9 | 15 | 6 | 10 | 13 | 10 |
|  | U2 | coupled | 17 | 12 | 8 | 19 | 17 | 11 | 13 | 12 | 8 |
|  |  | dendritic | 13 | 14 | 8 | 14 | 15 | 6 | 7 | 10 | 3 |
|  |  | somatic | 5 | 7 | 4 | 9 | 5 | 3 | 8 | 4 | 3 |
|  | U3 | coupled | 16 | 15 | 9 | 14 | 19 | 10 | 15 | 14 | 10 |
|  |  | dendritic | 15 | 9 | 4 | 14 | 12 | 8 | 11 | 10 | 3 |
|  |  | somatic | 4 | 4 | 3 | 9 | 12 | 7 | 8 | 8 | 7 |
| **VISpm** | U1 | coupled | 20 | 16 | 15 | 16 | 19 | 13 | 11 | 12 | 6 |
|  |  | dendritic | 6 | 9 | 5 | 10 | 9 | 5 | 7 | 5 | 1 |
|  |  | somatic | 9 | 8 | 5 | 10 | 16 | 8 | 10 | 6 | 5 |
|  | U2 | coupled | 14 | 10 | 9 | 13 | 17 | 12 | 15 | 16 | 14 |
|  |  | dendritic | 13 | 10 | 9 | 12 | 9 | 6 | 12 | 11 | 7 |
|  |  | somatic | 11 | 13 | 10 | 10 | 11 | 8 | 12 | 10 | 9 |
|  | U3 | coupled | 18 | 13 | 12 | 13 | 16 | 12 | 13 | 6 | 6 |
|  |  | dendritic | 10 | 10 | 6 | 9 | 8 | 6 | 11 | 8 | 7 |
|  |  | somatic | 12 | 10 | 8 | 9 | 8 | 4 | 7 | 9 | 6 |
| **VISam** | U1 | coupled | 16 | 9 | 9 | 13 | 17 | 13 | 10 | 13 | 5 |
|  |  | dendritic | 8 | 6 | 3 | 7 | 6 | 2 | 6 | 4 | 2 |
|  |  | somatic | 10 | 8 | 6 | 13 | 10 | 7 | 12 | 8 | 8 |
|  | U2 | coupled | 7 | 5 | 3 | 8 | 9 | 6 | 7 | 5 | 4 |
|  |  | dendritic | 4 | 4 | 3 | 6 | 7 | 5 | 5 | 7 | 4 |
|  |  | somatic | 3 | 7 | 3 | 5 | 7 | 2 | 5 | 5 | 1 |
|  | U3 | coupled | 6 | 5 | 2 | 12 | 10 | 7 | 9 | 11 | 7 |
|  |  | dendritic | 9 | 8 | 7 | 4 | 4 | 2 | 3 | 3 | 1 |
|  |  | somatic | 8 | 5 | 4 | 9 | 10 | 5 | 9 | 9 | 7 |

**Table S3:** L5 tracing-matched (confidence>=4), number of large calcium transients per area during natural movies (flower / worms / touch of evil) during each session, separated by condition (mismatch (mm) / control (ct) / intersection (int)) and transient class (Fig. 7). Intersection (int) includes all transients from pairs active in both mismatch (mm) and control (ct) conditions for a given stimulus.

|  |  | natural movie | flower |  |  | worms |  |  | touch of evil |  |  |
|---|---|---|---|---|---|---|---|---|---|---|---|
|  |  |  | ct | mm | int | ct | mm | int | ct | mm | int |
| area | day | category |  |  |  |  |  |  |  |  |  |
| **VISp** | U1 | coupled | 66 | 97 | 127 | 49 | 54 | 76 | 17 | 23 | 18 |
|  |  | dendritic | 21 | 50 | 44 | 19 | 28 | 21 | 14 | 19 | 13 |
|  |  | somatic | 44 | 45 | 70 | 28 | 37 | 39 | 16 | 23 | 18 |
|  | U2 | coupled | 62 | 43 | 78 | 39 | 34 | 44 | 37 | 35 | 37 |
|  |  | dendritic | 32 | 30 | 36 | 27 | 18 | 6 | 20 | 11 | 8 |
|  |  | somatic | 23 | 41 | 49 | 27 | 12 | 17 | 19 | 23 | 32 |
|  | U3 | coupled | 99 | 84 | 158 | 80 | 77 | 110 | 44 | 58 | 69 |
|  |  | dendritic | 40 | 40 | 45 | 48 | 42 | 71 | 33 | 34 | 33 |
|  |  | somatic | 71 | 52 | 106 | 47 | 50 | 69 | 45 | 47 | 67 |
| **VISl** | U1 | coupled | 38 | 45 | 66 | 29 | 27 | 35 | 31 | 41 | 58 |
|  |  | dendritic | 14 | 17 | 25 | 11 | 14 | 9 | 16 | 17 | 15 |
|  |  | somatic | 30 | 18 | 34 | 20 | 31 | 28 | 28 | 32 | 57 |
|  | U2 | coupled | 34 | 19 | 34 | 41 | 48 | 63 | 21 | 31 | 42 |
|  |  | dendritic | 15 | 19 | 22 | 21 | 21 | 21 | 11 | 12 | 9 |
|  |  | somatic | 12 | 15 | 23 | 13 | 11 | 13 | 20 | 13 | 21 |
|  | U3 | coupled | 25 | 31 | 39 | 32 | 44 | 55 | 45 | 47 | 80 |
|  |  | dendritic | 23 | 10 | 9 | 27 | 42 | 56 | 20 | 19 | 10 |
|  |  | somatic | 6 | 10 | 12 | 16 | 19 | 27 | 27 | 34 | 59 |
| **VISpm** | U1 | coupled | 61 | 34 | 85 | 27 | 44 | 61 | 22 | 23 | 26 |
|  |  | dendritic | 9 | 20 | 22 | 14 | 13 | 14 | 10 | 11 | 5 |
|  |  | somatic | 39 | 24 | 49 | 31 | 35 | 51 | 21 | 26 | 39 |
|  | U2 | coupled | 42 | 27 | 50 | 22 | 42 | 55 | 59 | 56 | 109 |
|  |  | dendritic | 19 | 21 | 34 | 24 | 18 | 25 | 32 | 24 | 47 |
|  |  | somatic | 31 | 36 | 61 | 38 | 36 | 67 | 36 | 42 | 72 |
|  | U3 | coupled | 58 | 38 | 84 | 25 | 30 | 49 | 37 | 26 | 50 |
|  |  | dendritic | 23 | 15 | 29 | 25 | 13 | 31 | 29 | 20 | 42 |
|  |  | somatic | 58 | 37 | 83 | 26 | 21 | 37 | 14 | 18 | 25 |
| **VISam** | U1 | coupled | 28 | 12 | 31 | 32 | 32 | 59 | 14 | 20 | 18 |
|  |  | dendritic | 16 | 16 | 23 | 10 | 6 | 6 | 7 | 7 | 4 |
|  |  | somatic | 19 | 17 | 25 | 52 | 36 | 73 | 35 | 20 | 45 |
|  | U2 | coupled | 9 | 7 | 7 | 16 | 12 | 20 | 13 | 13 | 20 |
|  |  | dendritic | 4 | 8 | 8 | 8 | 9 | 14 | 11 | 10 | 16 |
|  |  | somatic | 5 | 10 | 11 | 8 | 13 | 7 | 6 | 7 | 2 |
|  | U3 | coupled | 10 | 8 | 6 | 16 | 15 | 22 | 18 | 20 | 28 |
|  |  | dendritic | 13 | 14 | 24 | 9 | 10 | 12 | 4 | 3 | 3 |
|  |  | somatic | 17 | 9 | 18 | 20 | 20 | 26 | 19 | 15 | 27 |

**Table S4:** L5 tracing-matched (confidence>=4), number of pairs per area with transients during Gabor sequences (fixed / rotate) during each session, separated by condition (mismatch (mm) / control (ct) / intersection (both)) and transient class (Fig. S12). Intersection (int) includes all pairs with transients in both mismatch (mm) and control (ct) conditions for a given stimulus.

|  |  | Gabor type | fixed |  |  | rotate |  |  |
|---|---|---|---|---|---|---|---|---|
|  |  |  | ct | mm | both | ct | mm | both |
| area | day | class |  |  |  |  |  |  |
| **VISp** | U1 | coupled | 32 | 35 | 24 | 32 | 31 | 23 |
|  |  | dendritic | 24 | 23 | 14 | 27 | 23 | 16 |
|  |  | somatic | 22 | 25 | 17 | 28 | 23 | 14 |
|  | U2 | coupled | 29 | 31 | 20 | 30 | 27 | 22 |
|  |  | dendritic | 16 | 16 | 8 | 18 | 20 | 12 |
|  |  | somatic | 26 | 22 | 19 | 22 | 18 | 12 |
|  | U3 | coupled | 30 | 37 | 20 | 31 | 35 | 21 |
|  |  | dendritic | 33 | 29 | 20 | 25 | 22 | 13 |
|  |  | somatic | 18 | 23 | 15 | 27 | 22 | 19 |
| **VISl** | U1 | coupled | 22 | 26 | 17 | 23 | 22 | 12 |
|  |  | dendritic | 18 | 14 | 9 | 14 | 13 | 7 |
|  |  | somatic | 10 | 11 | 7 | 17 | 11 | 9 |
|  | U2 | coupled | 13 | 17 | 8 | 13 | 17 | 8 |
|  |  | dendritic | 18 | 15 | 8 | 9 | 14 | 7 |
|  |  | somatic | 8 | 9 | 5 | 8 | 4 | 3 |
|  | U3 | coupled | 17 | 15 | 9 | 23 | 14 | 12 |
|  |  | dendritic | 14 | 18 | 7 | 12 | 12 | 6 |
|  |  | somatic | 10 | 13 | 10 | 12 | 14 | 12 |
| **VISpm** | U1 | coupled | 22 | 20 | 18 | 23 | 20 | 18 |
|  |  | dendritic | 13 | 11 | 8 | 11 | 8 | 4 |
|  |  | somatic | 14 | 14 | 13 | 13 | 14 | 10 |
|  | U2 | coupled | 13 | 18 | 11 | 14 | 19 | 14 |
|  |  | dendritic | 14 | 12 | 8 | 10 | 11 | 6 |
|  |  | somatic | 13 | 16 | 12 | 14 | 12 | 12 |
|  | U3 | coupled | 17 | 15 | 14 | 15 | 15 | 9 |
|  |  | dendritic | 12 | 15 | 9 | 13 | 15 | 11 |
|  |  | somatic | 10 | 10 | 7 | 12 | 13 | 8 |
| **VISam** | U1 | coupled | 15 | 18 | 14 | 17 | 16 | 13 |
|  |  | dendritic | 8 | 8 | 5 | 7 | 7 | 5 |
|  |  | somatic | 14 | 12 | 10 | 18 | 13 | 12 |
|  | U2 | coupled | 8 | 8 | 6 | 7 | 8 | 5 |
|  |  | dendritic | 5 | 7 | 2 | 5 | 6 | 2 |
|  |  | somatic | 9 | 9 | 7 | 9 | 5 | 4 |
|  | U3 | coupled | 14 | 13 | 11 | 15 | 13 | 12 |
|  |  | dendritic | 8 | 10 | 6 | 9 | 10 | 7 |
|  |  | somatic | 12 | 14 | 11 | 15 | 15 | 13 |

**Table S5:**
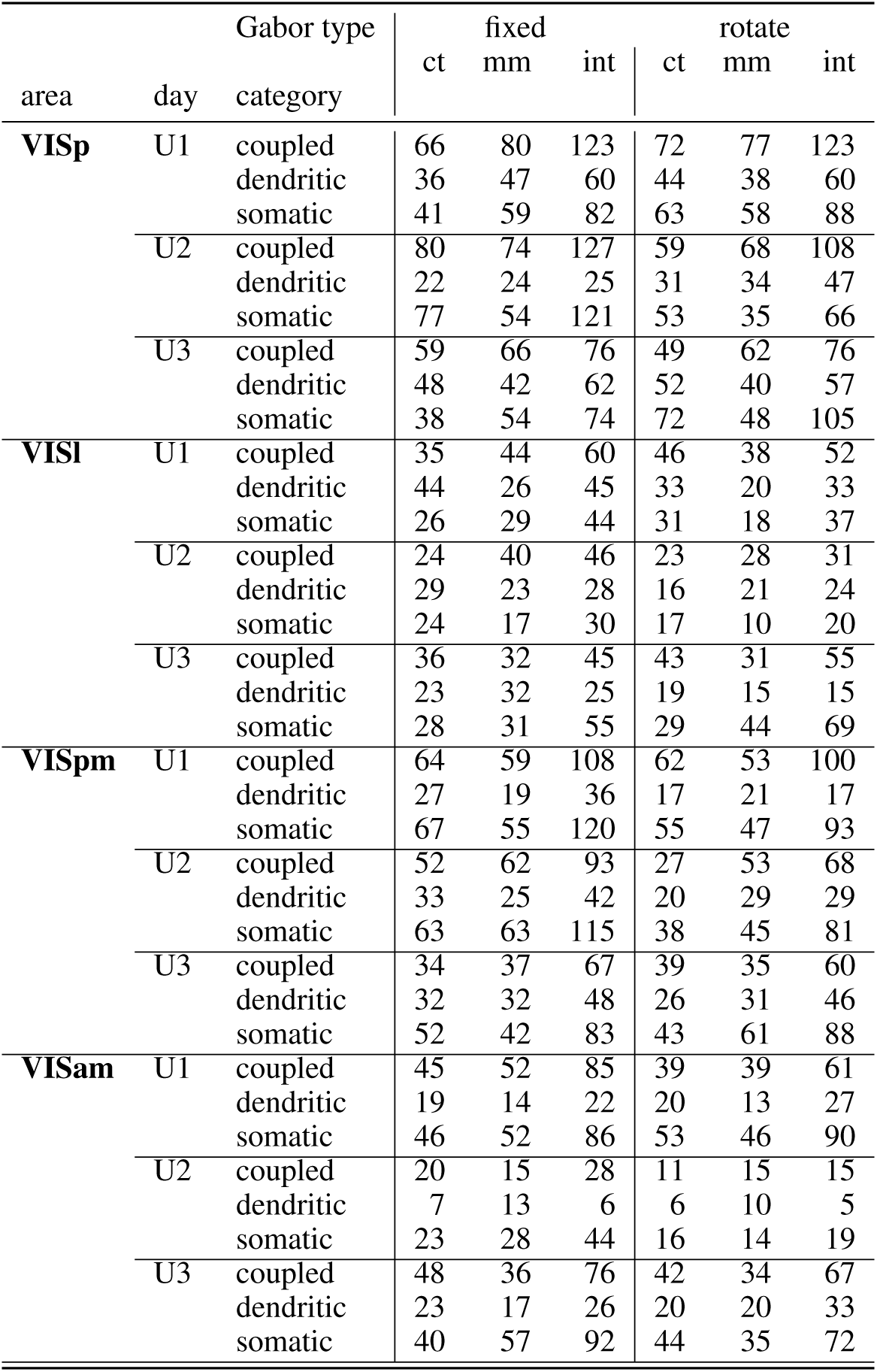
L5 tracing-matched (confidence>=4), number of large calcium transients per area during Gabor sequences (fixed / rotate) during each session, separated by condition (mismatch (mm) / control (ct) / intersection (int)) and transient class (Fig. S12). Intersection (int) includes all transients from pairs active in both mismatch (mm) and control (ct) conditions for a given stimulus.

|  |  | Gabor type | fixed |  |  | rotate |  |  |
|---|---|---|---|---|---|---|---|---|
|  |  |  | ct | mm | int | ct | mm | int |
| area | day | category |  |  |  |  |  |  |
| **VISp** | U1 | coupled | 66 | 80 | 123 | 72 | 77 | 123 |
|  |  | dendritic | 36 | 47 | 60 | 44 | 38 | 60 |
|  |  | somatic | 41 | 59 | 82 | 63 | 58 | 88 |
|  | U2 | coupled | 80 | 74 | 127 | 59 | 68 | 108 |
|  |  | dendritic | 22 | 24 | 25 | 31 | 34 | 47 |
|  |  | somatic | 77 | 54 | 121 | 53 | 35 | 66 |
|  | U3 | coupled | 59 | 66 | 76 | 49 | 62 | 76 |
|  |  | dendritic | 48 | 42 | 62 | 52 | 40 | 57 |
|  |  | somatic | 38 | 54 | 74 | 72 | 48 | 105 |
| **VISl** | U1 | coupled | 35 | 44 | 60 | 46 | 38 | 52 |
|  |  | dendritic | 44 | 26 | 45 | 33 | 20 | 33 |
|  |  | somatic | 26 | 29 | 44 | 31 | 18 | 37 |
|  | U2 | coupled | 24 | 40 | 46 | 23 | 28 | 31 |
|  |  | dendritic | 29 | 23 | 28 | 16 | 21 | 24 |
|  |  | somatic | 24 | 17 | 30 | 17 | 10 | 20 |
|  | U3 | coupled | 36 | 32 | 45 | 43 | 31 | 55 |
|  |  | dendritic | 23 | 32 | 25 | 19 | 15 | 15 |
|  |  | somatic | 28 | 31 | 55 | 29 | 44 | 69 |
| **VISpm** | U1 | coupled | 64 | 59 | 108 | 62 | 53 | 100 |
|  |  | dendritic | 27 | 19 | 36 | 17 | 21 | 17 |
|  |  | somatic | 67 | 55 | 120 | 55 | 47 | 93 |
|  | U2 | coupled | 52 | 62 | 93 | 27 | 53 | 68 |
|  |  | dendritic | 33 | 25 | 42 | 20 | 29 | 29 |
|  |  | somatic | 63 | 63 | 115 | 38 | 45 | 81 |
|  | U3 | coupled | 34 | 37 | 67 | 39 | 35 | 60 |
|  |  | dendritic | 32 | 32 | 48 | 26 | 31 | 46 |
|  |  | somatic | 52 | 42 | 83 | 43 | 61 | 88 |
| **VISam** | U1 | coupled | 45 | 52 | 85 | 39 | 39 | 61 |
|  |  | dendritic | 19 | 14 | 22 | 20 | 13 | 27 |
|  |  | somatic | 46 | 52 | 86 | 53 | 46 | 90 |
|  | U2 | coupled | 20 | 15 | 28 | 11 | 15 | 15 |
|  |  | dendritic | 7 | 13 | 6 | 6 | 10 | 5 |
|  |  | somatic | 23 | 28 | 44 | 16 | 14 | 19 |
|  | U3 | coupled | 48 | 36 | 76 | 42 | 34 | 67 |
|  |  | dendritic | 23 | 17 | 26 | 20 | 20 | 33 |
|  |  | somatic | 40 | 57 | 92 | 44 | 35 | 72 |

**Table S6:**
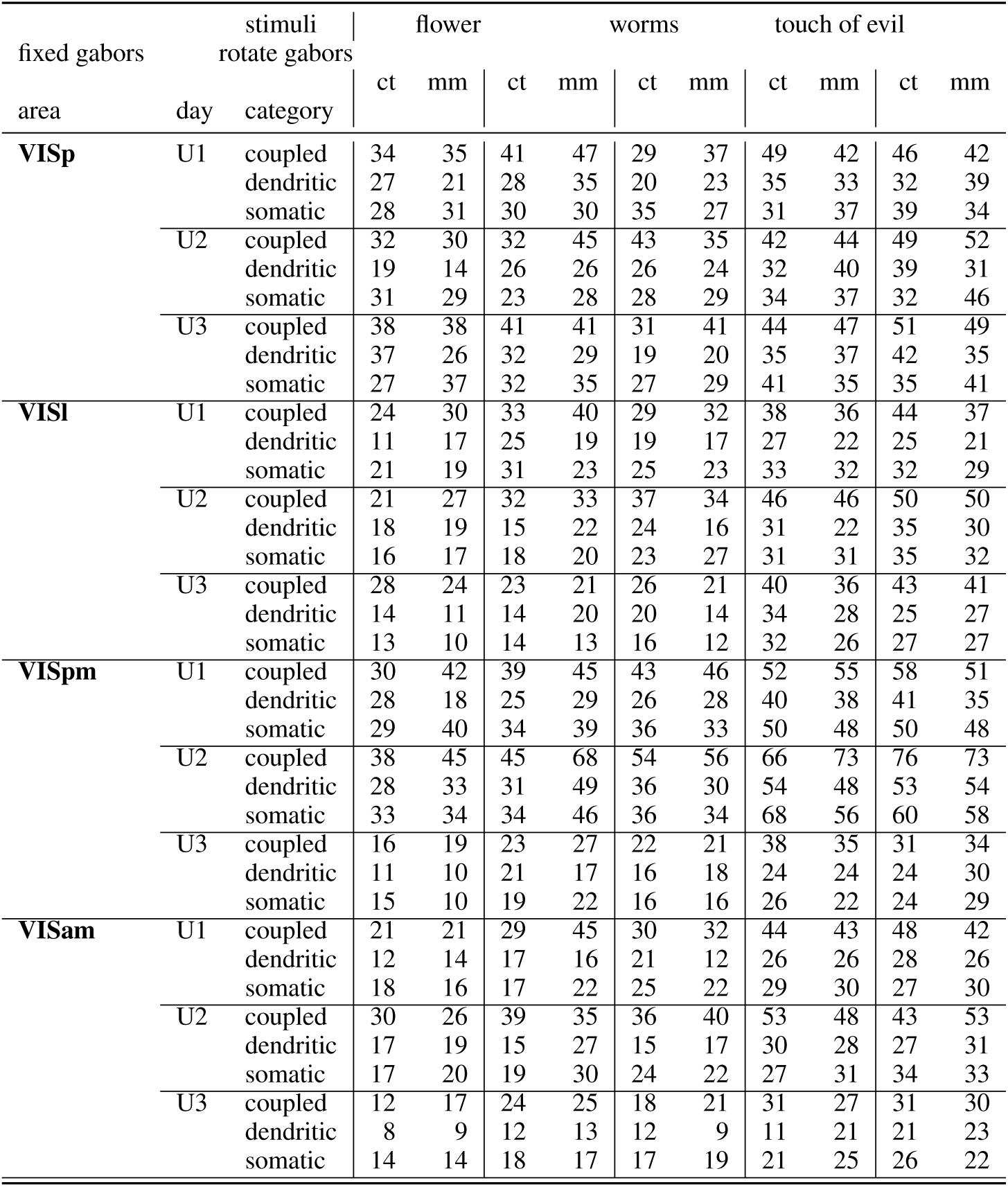
L2/3 pseudo-matching, number of pairs (n) per area included in each comparison (Fig. S13 A, B, Fig. S14 A, B, Fig. S15 A, B) (i.e., with transients of the coupled (coup.), somatic (soma.) or dendritic (dend.) class in each session, for stimulus and condition (mismatch (mm) / control (ct)).

|  |  | stimuli | flower |  |  |  | worms |  | touch of evil |  |  |  |
|---|---|---|---|---|---|---|---|---|---|---|---|---|
| fixed gabors |  | rotate gabors |  |  |  |  |  |  |  |  |  |  |
|  |  |  | ct | mm | ct | mm | ct | mm | ct | mm | ct | mm |
| area | day | category |  |  |  |  |  |  |  |  |  |  |
| **VISp** | U1 | coupled | 34 | 35 | 41 | 47 | 29 | 37 | 49 | 42 | 46 | 42 |
|  |  | dendritic | 27 | 21 | 28 | 35 | 20 | 23 | 35 | 33 | 32 | 39 |
|  |  | somatic | 28 | 31 | 30 | 30 | 35 | 27 | 31 | 37 | 39 | 34 |
|  | U2 | coupled | 32 | 30 | 32 | 45 | 43 | 35 | 42 | 44 | 49 | 52 |
|  |  | dendritic | 19 | 14 | 26 | 26 | 26 | 24 | 32 | 40 | 39 | 31 |
|  |  | somatic | 31 | 29 | 23 | 28 | 28 | 29 | 34 | 37 | 32 | 46 |
|  | U3 | coupled | 38 | 38 | 41 | 41 | 31 | 41 | 44 | 47 | 51 | 49 |
|  |  | dendritic | 37 | 26 | 32 | 29 | 19 | 20 | 35 | 37 | 42 | 35 |
|  |  | somatic | 27 | 37 | 32 | 35 | 27 | 29 | 41 | 35 | 35 | 41 |
| **VISl** | U1 | coupled | 24 | 30 | 33 | 40 | 29 | 32 | 38 | 36 | 44 | 37 |
|  |  | dendritic | 11 | 17 | 25 | 19 | 19 | 17 | 27 | 22 | 25 | 21 |
|  |  | somatic | 21 | 19 | 31 | 23 | 25 | 23 | 33 | 32 | 32 | 29 |
|  | U2 | coupled | 21 | 27 | 32 | 33 | 37 | 34 | 46 | 46 | 50 | 50 |
|  |  | dendritic | 18 | 19 | 15 | 22 | 24 | 16 | 31 | 22 | 35 | 30 |
|  |  | somatic | 16 | 17 | 18 | 20 | 23 | 27 | 31 | 31 | 35 | 32 |
|  | U3 | coupled | 28 | 24 | 23 | 21 | 26 | 21 | 40 | 36 | 43 | 41 |
|  |  | dendritic | 14 | 11 | 14 | 20 | 20 | 14 | 34 | 28 | 25 | 27 |
|  |  | somatic | 13 | 10 | 14 | 13 | 16 | 12 | 32 | 26 | 27 | 27 |
| **VISpm** | U1 | coupled | 30 | 42 | 39 | 45 | 43 | 46 | 52 | 55 | 58 | 51 |
|  |  | dendritic | 28 | 18 | 25 | 29 | 26 | 28 | 40 | 38 | 41 | 35 |
|  |  | somatic | 29 | 40 | 34 | 39 | 36 | 33 | 50 | 48 | 50 | 48 |
|  | U2 | coupled | 38 | 45 | 45 | 68 | 54 | 56 | 66 | 73 | 76 | 73 |
|  |  | dendritic | 28 | 33 | 31 | 49 | 36 | 30 | 54 | 48 | 53 | 54 |
|  |  | somatic | 33 | 34 | 34 | 46 | 36 | 34 | 68 | 56 | 60 | 58 |
|  | U3 | coupled | 16 | 19 | 23 | 27 | 22 | 21 | 38 | 35 | 31 | 34 |
|  |  | dendritic | 11 | 10 | 21 | 17 | 16 | 18 | 24 | 24 | 24 | 30 |
|  |  | somatic | 15 | 10 | 19 | 22 | 16 | 16 | 26 | 22 | 24 | 29 |
| **VISam** | U1 | coupled | 21 | 21 | 29 | 45 | 30 | 32 | 44 | 43 | 48 | 42 |
|  |  | dendritic | 12 | 14 | 17 | 16 | 21 | 12 | 26 | 26 | 28 | 26 |
|  |  | somatic | 18 | 16 | 17 | 22 | 25 | 22 | 29 | 30 | 27 | 30 |
|  | U2 | coupled | 30 | 26 | 39 | 35 | 36 | 40 | 53 | 48 | 43 | 53 |
|  |  | dendritic | 17 | 19 | 15 | 27 | 15 | 17 | 30 | 28 | 27 | 31 |
|  |  | somatic | 17 | 20 | 19 | 30 | 24 | 22 | 27 | 31 | 34 | 33 |
|  | U3 | coupled | 12 | 17 | 24 | 25 | 18 | 21 | 31 | 27 | 31 | 30 |
|  |  | dendritic | 8 | 9 | 12 | 13 | 12 | 9 | 11 | 21 | 21 | 23 |
|  |  | somatic | 14 | 14 | 18 | 17 | 17 | 19 | 21 | 25 | 26 | 22 |

**Table S7:**
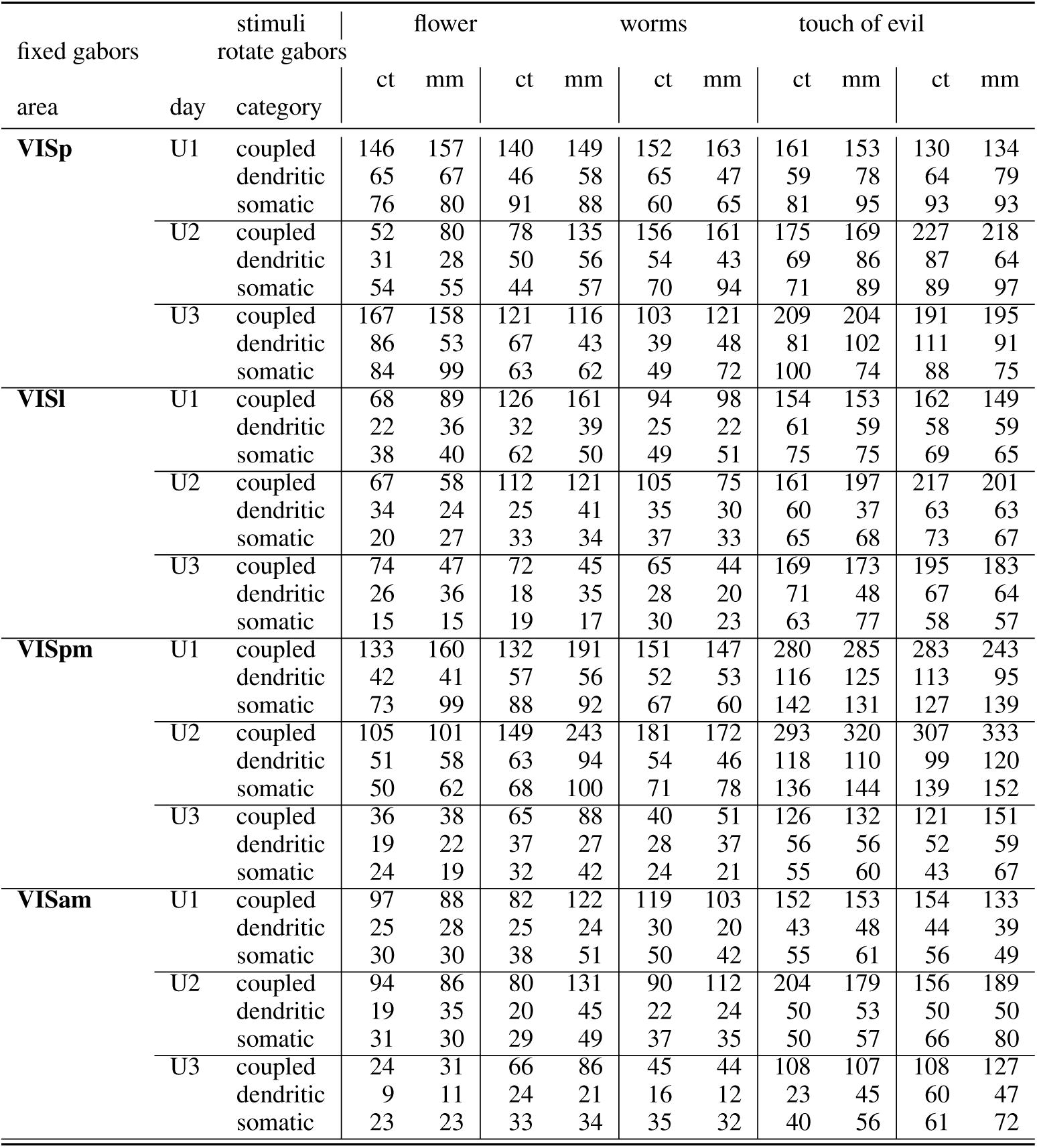
L2/3 pseudo-matching, number of large calcium transients per area included in each comparison (Fig. S13 A, B, Fig. S14 A, B, Fig. S15 A, B) (i.e., with transients of the coupled (coup.), somatic (soma.) or dendritic (dend.) class in each session, for stimulus and condition (mismatch (mm) / control (ct)).

|  |  | stimuli | flower |  |  |  | worms |  | touch of evil |  |  |  |
|---|---|---|---|---|---|---|---|---|---|---|---|---|
| fixed gabors |  | rotate gabors |  |  |  |  |  |  |  |  |  |  |
|  |  |  | ct | mm | ct | mm | ct | mm | ct | mm | ct | mm |
| area | day | category |  |  |  |  |  |  |  |  |  |  |
| **VISp** | U1 | coupled | 146 | 157 | 140 | 149 | 152 | 163 | 161 | 153 | 130 | 134 |
|  |  | dendritic | 65 | 67 | 46 | 58 | 65 | 47 | 59 | 78 | 64 | 79 |
|  |  | somatic | 76 | 80 | 91 | 88 | 60 | 65 | 81 | 95 | 93 | 93 |
|  | U2 | coupled | 52 | 80 | 78 | 135 | 156 | 161 | 175 | 169 | 227 | 218 |
|  |  | dendritic | 31 | 28 | 50 | 56 | 54 | 43 | 69 | 86 | 87 | 64 |
|  |  | somatic | 54 | 55 | 44 | 57 | 70 | 94 | 71 | 89 | 89 | 97 |
|  | U3 | coupled | 167 | 158 | 121 | 116 | 103 | 121 | 209 | 204 | 191 | 195 |
|  |  | dendritic | 86 | 53 | 67 | 43 | 39 | 48 | 81 | 102 | 111 | 91 |
|  |  | somatic | 84 | 99 | 63 | 62 | 49 | 72 | 100 | 74 | 88 | 75 |
| **VISl** | U1 | coupled | 68 | 89 | 126 | 161 | 94 | 98 | 154 | 153 | 162 | 149 |
|  |  | dendritic | 22 | 36 | 32 | 39 | 25 | 22 | 61 | 59 | 58 | 59 |
|  |  | somatic | 38 | 40 | 62 | 50 | 49 | 51 | 75 | 75 | 69 | 65 |
|  | U2 | coupled | 67 | 58 | 112 | 121 | 105 | 75 | 161 | 197 | 217 | 201 |
|  |  | dendritic | 34 | 24 | 25 | 41 | 35 | 30 | 60 | 37 | 63 | 63 |
|  |  | somatic | 20 | 27 | 33 | 34 | 37 | 33 | 65 | 68 | 73 | 67 |
|  | U3 | coupled | 74 | 47 | 72 | 45 | 65 | 44 | 169 | 173 | 195 | 183 |
|  |  | dendritic | 26 | 36 | 18 | 35 | 28 | 20 | 71 | 48 | 67 | 64 |
|  |  | somatic | 15 | 15 | 19 | 17 | 30 | 23 | 63 | 77 | 58 | 57 |
| **VISpm** | U1 | coupled | 133 | 160 | 132 | 191 | 151 | 147 | 280 | 285 | 283 | 243 |
|  |  | dendritic | 42 | 41 | 57 | 56 | 52 | 53 | 116 | 125 | 113 | 95 |
|  |  | somatic | 73 | 99 | 88 | 92 | 67 | 60 | 142 | 131 | 127 | 139 |
|  | U2 | coupled | 105 | 101 | 149 | 243 | 181 | 172 | 293 | 320 | 307 | 333 |
|  |  | dendritic | 51 | 58 | 63 | 94 | 54 | 46 | 118 | 110 | 99 | 120 |
|  |  | somatic | 50 | 62 | 68 | 100 | 71 | 78 | 136 | 144 | 139 | 152 |
|  | U3 | coupled | 36 | 38 | 65 | 88 | 40 | 51 | 126 | 132 | 121 | 151 |
|  |  | dendritic | 19 | 22 | 37 | 27 | 28 | 37 | 56 | 56 | 52 | 59 |
|  |  | somatic | 24 | 19 | 32 | 42 | 24 | 21 | 55 | 60 | 43 | 67 |
| **VISam** | U1 | coupled | 97 | 88 | 82 | 122 | 119 | 103 | 152 | 153 | 154 | 133 |
|  |  | dendritic | 25 | 28 | 25 | 24 | 30 | 20 | 43 | 48 | 44 | 39 |
|  |  | somatic | 30 | 30 | 38 | 51 | 50 | 42 | 55 | 61 | 56 | 49 |
|  | U2 | coupled | 94 | 86 | 80 | 131 | 90 | 112 | 204 | 179 | 156 | 189 |
|  |  | dendritic | 19 | 35 | 20 | 45 | 22 | 24 | 50 | 53 | 50 | 50 |
|  |  | somatic | 31 | 30 | 29 | 49 | 37 | 35 | 50 | 57 | 66 | 80 |
|  | U3 | coupled | 24 | 31 | 66 | 86 | 45 | 44 | 108 | 107 | 108 | 127 |
|  |  | dendritic | 9 | 11 | 24 | 21 | 16 | 12 | 23 | 45 | 60 | 47 |
|  |  | somatic | 23 | 23 | 33 | 34 | 35 | 32 | 40 | 56 | 61 | 72 |

**Table S8:**
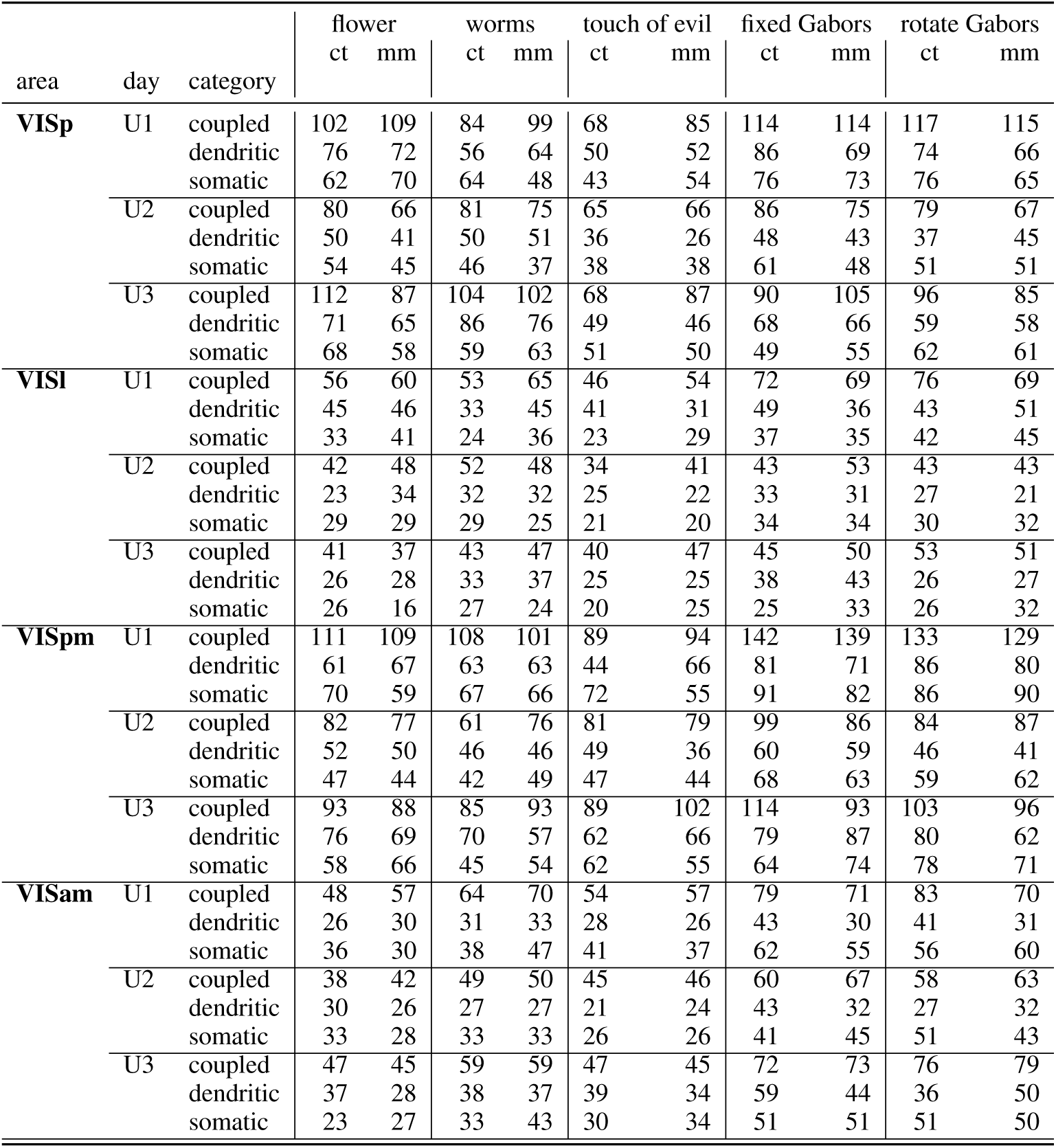
L5 pseudo-matching, number of pairs (n) per area included in each comparison (Fig. S13 C, D, Fig. S14 C, D, Fig. S15 C, D) (i.e., with transients of the coupled (coup.), somatic (soma.) or dendritic (dend.) class in each session, for stimulus and condition (mismatch (mm) / control (ct)).

|  |  |  | flower |  | worms |  | touch of evil |  | fixed Gabors |  | rotate Gabors |  |
|---|---|---|---|---|---|---|---|---|---|---|---|---|
|  |  |  | ct | mm | ct | mm | ct | mm | ct | mm | ct | mm |
| area | day | category |  |  |  |  |  |  |  |  |  |  |
| **VISp** | U1 | coupled | 102 | 109 | 84 | 99 | 68 | 85 | 114 | 114 | 117 | 115 |
|  |  | dendritic | 76 | 72 | 56 | 64 | 50 | 52 | 86 | 69 | 74 | 66 |
|  |  | somatic | 62 | 70 | 64 | 48 | 43 | 54 | 76 | 73 | 76 | 65 |
|  | U2 | coupled | 80 | 66 | 81 | 75 | 65 | 66 | 86 | 75 | 79 | 67 |
|  |  | dendritic | 50 | 41 | 50 | 51 | 36 | 26 | 48 | 43 | 37 | 45 |
|  |  | somatic | 54 | 45 | 46 | 37 | 38 | 38 | 61 | 48 | 51 | 51 |
|  | U3 | coupled | 112 | 87 | 104 | 102 | 68 | 87 | 90 | 105 | 96 | 85 |
|  |  | dendritic | 71 | 65 | 86 | 76 | 49 | 46 | 68 | 66 | 59 | 58 |
|  |  | somatic | 68 | 58 | 59 | 63 | 51 | 50 | 49 | 55 | 62 | 61 |
| **VISl** | U1 | coupled | 56 | 60 | 53 | 65 | 46 | 54 | 72 | 69 | 76 | 69 |
|  |  | dendritic | 45 | 46 | 33 | 45 | 41 | 31 | 49 | 36 | 43 | 51 |
|  |  | somatic | 33 | 41 | 24 | 36 | 23 | 29 | 37 | 35 | 42 | 45 |
|  | U2 | coupled | 42 | 48 | 52 | 48 | 34 | 41 | 43 | 53 | 43 | 43 |
|  |  | dendritic | 23 | 34 | 32 | 32 | 25 | 22 | 33 | 31 | 27 | 21 |
|  |  | somatic | 29 | 29 | 29 | 25 | 21 | 20 | 34 | 34 | 30 | 32 |
|  | U3 | coupled | 41 | 37 | 43 | 47 | 40 | 47 | 45 | 50 | 53 | 51 |
|  |  | dendritic | 26 | 28 | 33 | 37 | 25 | 25 | 38 | 43 | 26 | 27 |
|  |  | somatic | 26 | 16 | 27 | 24 | 20 | 25 | 25 | 33 | 26 | 32 |
| **VISpm** | U1 | coupled | 111 | 109 | 108 | 101 | 89 | 94 | 142 | 139 | 133 | 129 |
|  |  | dendritic | 61 | 67 | 63 | 63 | 44 | 66 | 81 | 71 | 86 | 80 |
|  |  | somatic | 70 | 59 | 67 | 66 | 72 | 55 | 91 | 82 | 86 | 90 |
|  | U2 | coupled | 82 | 77 | 61 | 76 | 81 | 79 | 99 | 86 | 84 | 87 |
|  |  | dendritic | 52 | 50 | 46 | 46 | 49 | 36 | 60 | 59 | 46 | 41 |
|  |  | somatic | 47 | 44 | 42 | 49 | 47 | 44 | 68 | 63 | 59 | 62 |
|  | U3 | coupled | 93 | 88 | 85 | 93 | 89 | 102 | 114 | 93 | 103 | 96 |
|  |  | dendritic | 76 | 69 | 70 | 57 | 62 | 66 | 79 | 87 | 80 | 62 |
|  |  | somatic | 58 | 66 | 45 | 54 | 62 | 55 | 64 | 74 | 78 | 71 |
| **VISam** | U1 | coupled | 48 | 57 | 64 | 70 | 54 | 57 | 79 | 71 | 83 | 70 |
|  |  | dendritic | 26 | 30 | 31 | 33 | 28 | 26 | 43 | 30 | 41 | 31 |
|  |  | somatic | 36 | 30 | 38 | 47 | 41 | 37 | 62 | 55 | 56 | 60 |
|  | U2 | coupled | 38 | 42 | 49 | 50 | 45 | 46 | 60 | 67 | 58 | 63 |
|  |  | dendritic | 30 | 26 | 27 | 27 | 21 | 24 | 43 | 32 | 27 | 32 |
|  |  | somatic | 33 | 28 | 33 | 33 | 26 | 26 | 41 | 45 | 51 | 43 |
|  | U3 | coupled | 47 | 45 | 59 | 59 | 47 | 45 | 72 | 73 | 76 | 79 |
|  |  | dendritic | 37 | 28 | 38 | 37 | 39 | 34 | 59 | 44 | 36 | 50 |
|  |  | somatic | 23 | 27 | 33 | 43 | 30 | 34 | 51 | 51 | 51 | 50 |

**Table S9:** L5 pseudo-matching, number of large calcium transients per area included in each comparison (Fig. S13 C, D, Fig. S14 C, D, Fig. S15 C, D) (i.e., with transients of the coupled (coup.), somatic (soma.) or dendritic (dend.) class in each session, for stimulus and condition (mismatch (mm) / control (ct)).

| area | day | category | flower |  | worms |  | touch of evil |  | fixed Gabors |  | rotate Gabors |  |
|---|---|---|---|---|---|---|---|---|---|---|---|---|
|  |  |  | ct | mm | ct | mm | ct | mm | ct | mm | ct | mm |
| **VISp** | U1 | coupled | 221 | 328 | 154 | 240 | 117 | 173 | 242 | 262 | 276 | 276 |
|  |  | dendritic | 123 | 146 | 74 | 103 | 75 | 70 | 144 | 119 | 110 | 109 |
|  |  | somatic | 115 | 135 | 114 | 86 | 91 | 96 | 143 | 150 | 156 | 141 |
|  | U2 | coupled | 241 | 161 | 194 | 196 | 143 | 126 | 209 | 171 | 143 | 118 |
|  |  | dendritic | 77 | 67 | 79 | 79 | 49 | 36 | 87 | 79 | 50 | 58 |
|  |  | somatic | 119 | 111 | 104 | 88 | 85 | 67 | 114 | 108 | 98 | 79 |
|  | U3 | coupled | 294 | 237 | 304 | 285 | 149 | 176 | 183 | 211 | 214 | 171 |
|  |  | dendritic | 128 | 108 | 155 | 146 | 74 | 74 | 107 | 113 | 99 | 93 |
|  |  | somatic | 138 | 128 | 119 | 120 | 87 | 93 | 85 | 101 | 107 | 89 |
| **VISl** | U1 | coupled | 170 | 195 | 126 | 144 | 124 | 159 | 153 | 149 | 201 | 157 |
|  |  | dendritic | 73 | 77 | 53 | 77 | 58 | 46 | 79 | 57 | 66 | 91 |
|  |  | somatic | 63 | 65 | 49 | 59 | 57 | 49 | 75 | 71 | 75 | 80 |
|  | U2 | coupled | 173 | 150 | 205 | 205 | 103 | 120 | 165 | 190 | 144 | 138 |
|  |  | dendritic | 38 | 57 | 71 | 69 | 48 | 41 | 68 | 64 | 57 | 58 |
|  |  | somatic | 60 | 61 | 45 | 46 | 41 | 39 | 96 | 90 | 68 | 82 |
|  | U3 | coupled | 85 | 85 | 161 | 189 | 102 | 136 | 120 | 145 | 124 | 119 |
|  |  | dendritic | 40 | 40 | 54 | 64 | 37 | 38 | 72 | 82 | 50 | 47 |
|  |  | somatic | 38 | 38 | 53 | 58 | 71 | 77 | 43 | 55 | 52 | 56 |
| **VISpm** | U1 | coupled | 314 | 285 | 213 | 246 | 212 | 208 | 323 | 400 | 344 | 336 |
|  |  | dendritic | 109 | 119 | 94 | 102 | 62 | 87 | 126 | 147 | 155 | 148 |
|  |  | somatic | 168 | 122 | 137 | 148 | 136 | 118 | 238 | 218 | 235 | 236 |
|  | U2 | coupled | 194 | 196 | 110 | 155 | 195 | 186 | 289 | 243 | 175 | 186 |
|  |  | dendritic | 76 | 88 | 63 | 67 | 81 | 46 | 105 | 86 | 68 | 60 |
|  |  | somatic | 79 | 83 | 70 | 77 | 84 | 85 | 165 | 145 | 136 | 123 |
|  | U3 | coupled | 300 | 258 | 235 | 273 | 323 | 380 | 272 | 234 | 264 | 220 |
|  |  | dendritic | 136 | 125 | 128 | 136 | 111 | 136 | 186 | 203 | 146 | 114 |
|  |  | somatic | 166 | 138 | 93 | 123 | 153 | 130 | 177 | 165 | 188 | 177 |
| **VISam** | U1 | coupled | 117 | 126 | 180 | 181 | 102 | 132 | 205 | 162 | 218 | 197 |
|  |  | dendritic | 30 | 43 | 50 | 46 | 36 | 32 | 65 | 52 | 62 | 43 |
|  |  | somatic | 66 | 47 | 121 | 143 | 84 | 74 | 137 | 121 | 147 | 132 |
|  | U2 | coupled | 106 | 126 | 123 | 122 | 96 | 104 | 157 | 167 | 124 | 158 |
|  |  | dendritic | 43 | 37 | 32 | 39 | 23 | 30 | 63 | 47 | 44 | 47 |
|  |  | somatic | 78 | 70 | 74 | 79 | 53 | 61 | 99 | 112 | 109 | 97 |
|  | U3 | coupled | 134 | 91 | 120 | 134 | 127 | 96 | 171 | 180 | 179 | 194 |
|  |  | dendritic | 71 | 48 | 49 | 45 | 57 | 56 | 99 | 70 | 60 | 75 |
|  |  | somatic | 61 | 56 | 67 | 106 | 66 | 62 | 113 | 115 | 131 | 111 |

## Notes

https://dandiarchive.org/dandiset/000336/

### Competing Interest Statement

The authors have declared no competing interest.

